# Expanding all-α-helical protein space through rational computational design

**DOI:** 10.64898/2026.07.03.736327

**Authors:** Katherine I. Albanese, Joel J. Chubb, Luis I Gutierrez-Rus, Xiyue Leng, Kathleen W. Kurgan, Bram Mylemans, Katarzyna Ożga, Rokas Petrenas, Andrey V. Romanyuk, Amanda M. Acevedo-Jake, Joel Roca-Martinez, Stephen J. Cross, J. L. Ross Anderson, Jonathan Clayden, Graham J. Leggett, Jennifer J. McManus, Thomas A. A. Oliver, Christine A. Orengo, Nigel S. Scrutton, Andrew J. Wilson, Aimee L. Boyle, Derek N. Woolfson

## Abstract

*De novo* protein design is advancing rapidly^1,2^. This is being driven by AI to generate protein backbones, sequences, and structural models^3–7^. As a result, *de novo* designed proteins are becoming larger and more complex^8–10^, and increasingly explore new protein structures^11,12^. By contrast, natural proteins have evolved structural and functional complexity by modular combination of recurring protein domains^13^. Approximately 25% of these natural domains are mostly α-helical structures^14^. Here we show how these can be expanded using rational computational design. Following the domain classification scheme CATH^15^, we build complex all-α *de novo* proteins hierarchically using sequence-to-structure relationships for helix-helix interactions, systematic rules to connect helices, computational tools to design loops, and *in silico* evaluation. The pipeline starts with a target architecture of free-standing helices. These are connected into a topology by considering local arrangements of helical bundles using understood sequence-to-structure relationships for helix packing. Single-chain sequences are completed using template- and AI-based methods. Finally, AlphaFold models are assessed to give small numbers of designs for experimental validation. We test 31 designs for 14 different architectures and 25 topologies. 75% of these express as stable, monomeric, water-soluble proteins; and >30% yield X-ray crystal structures matching the designs to atomic accuracy and with new-to-nature structures. Finally, several of the scaffolds are functionalised through one-shot designs to deliver ion, small-molecule and protein binders.

## Main

Natural proteins adopt many seemingly complex structures for a wide variety of functions, including as structural supports, catalysts, and for molecular recognition. *De novo* protein design attempts to capture and reduce this complexity by writing down completely new protein sequences from scratch. Protein design was founded with two main objectives: (i) to deepen understanding of sequence-to-structure/function relationships of natural proteins; and (ii) to deliver new-to-nature protein structures and functions with the potential to go beyond the those realised in natural proteins^1,2,16,17^.

Until recently, rational (rules based) and computational (bioinformatics and physics based) approaches dominated protein design^16–18^. Now, statistics-based, deep-learning methods are accelerating protein design and making it accessible to non-specialists^1,2^. This builds on major advances in protein-structure prediction (e.g. AlphaFold^6,7^, RoseTTAFold^19^), protein-backbone generation (RFDiffusion^20,21^), sequence generation (ProteinMPNN^4,5^), and integrated generative frameworks (Chroma and BoltzGen^22,23^). These advances are enabling rapid pipelines for structure-function design, with notable successes in designing *de novo* binders for small molecules^3,5^ and proteins^24^, as well as catalytically active proteins^25^. This is driving a paradigm shift where a target function is defined first and then used to generate protein scaffolds and sequences that satisfy the design objective^1,26^.

Nonetheless, regarding the two founding objectives of *de novo* design, current AI-based protein design has some limitations, which present new challenges. First, it is not clear to what extent rules can be extracted from generative-AI pipelines, and, thus, how they are advancing understanding of sequence-to-structure/function relationships. Second, while focussing on specified design constraints, AI methods can produce proteins that lack structural and functional nuances and complexity, such as flexibility and conformational exchange, which underpin natural protein functions such as regulation and allostery. Related to both issues, generative methods do not yet offer tight control over physicochemical properties of the design target, limiting the ability to design proteins for specific contexts. This reduces predictability and success rates and constrains applications, although progress is being made through approaches that incorporate large-scale experimental feedback to improve and refine models^27,28^.

Rational protein design offers a complementary route to address these issues. By building on fundamental principles of protein structure, rational approaches attempt to design proteins where the role of each residue is understood from the start^2,17^. The difficulty is that our understanding of sequence-to-structure/function relationships is incomplete. Thus, combining rational design (to define aspects that we do understand) with generative-AI approaches (to complete those that we do not) provides a possible path to more successful *de novo* protein design^11^. Moreover, such an integrated approach should enable increasing structural and functional complexity of design targets while retaining interpretability and predictability^2^.

α−Helical assemblies and proteins, such as coiled coils (CCs)^29^ and repeat proteins^30,31^, provide good targets for an integrated approach, as they are abundant and at least some understood sequence-to-structure relationships are available^29,32^; *i.e.*, there are a lot of data to facilitate both rational and generative design. For CCs, this has enabled the rational design of α-helical peptide assemblies with defined oligomeric states, helix orientations, and associated properties such as accessible and functionalisable central channels^33–37^. In turn, these peptide assemblies have been used as seeds to generate *de novo* single-chain coiled-coil proteins (scCCs) by combining rational, computational and AI-based design^11,38,39^. The resulting *de novo* CC peptides and proteins provide versatile scaffolds for rapidly introducing functions, including small-molecule binding^34,40^, protein binding^41^, ion-channel formation^42^, catalysis^43,44^, and for driving and intervening processes in mammalian cells^45,46^.

In nature, many proteins achieve higher levels of structural and functional complexity through the reuse and combination of domains^47^. Protein domains are generally defined as compact, semi-independent modules that fold autonomously and perform distinct functions. As a result, natural protein structures can be described hierarchically using domain-based classification systems such as CATH^15,48^ and SCOP^49,50^. For instance, CATH has four levels — <u>C</u>lass, <u>A</u>rchitecture, <u>T</u>opology, and <u>H</u>omologous superfamily — that provide a systematic description of domain structures and how they are related. Combining multiple domains into single chains leads to large, complex proteins that can harbour multiple functions. Indeed, up to 66% of prokaryotic and 80% of eukaryotic proteins have two or more domains^32^.

Here, we set out to test whether rationally designed scCCs could be combined to build more-complex structures and functions distinct from natural multidomain proteins. To do this, we introduce a rational strategy for designing multi-coiled-coil proteins (mCCs) in which scCC modules are not simply connected by linkers but merged through shared bifaceted helices that encode sequence-to-structure rules from two helical-bundle domains simultaneously. This generates continuous, modular architectures with defined structural and functional links between the constituent modules. Our pipeline builds directly on understood sequence-to-structure relationships of CC oligomers, extends scCC design principles, and integrates computational loop design, AI-assisted modelling and *in silico* validation. Through this, we deliver mCCs with diverse architectures and topologies predictably and with high experimental success rates. Structural similarity searches using FoldSeek^45^ indicate that most resulting mCCs occupy regions of all-α protein space not represented by natural proteins. Finally, we demonstrate that individual modules or subdomains can be functionalised independently or together to produce multifunctional mCCs, with examples of protein binding, small-molecule recognition and intracellular visualisation.

### Defining and designing multi-coiled-coil structures

In natural multi-domain proteins, consecutive domains are usually joined by flexible or rigid linkers, maintaining the structural independence of each module^13^. We and others have used both strategies to generate functional multi-domain coiled-coil proteins from oligomeric or single-chain helical bundles^40,46,51,52^. *De novo* protein design presents a third option: in principle, modules can be *merged* by sharing secondary structure units—for instance, bifaceted helices^53^—making the connection between them direct and defined. We sought to test this by systematically generating a series of new all-α-helical architectures and topologies.

By analogy to the CATH hierarchy^48,54^, we introduce CATS (<u>C</u>lass, <u>A</u>rchitecture, <u>T</u>opology, and <u>S</u>imilarity) as a classification system for *de novo* proteins (Figure 1A). We focussed on the all-α-helical class and specifically on coiled-coil-based helix-helix interactions^29^. This allowed rapid sequence and model generation by exploiting understood sequence-to-structure relationships and their underpinning knobs-into-holes (KIH) helix-helix interactions^29^. We define architecture as the number and overall 3D arrangement of helices, and topology as the order in which the helices are connected. Similarity replaces homology in CATH to capture the idea that whilst completely *de novo* designed proteins cannot share evolutionary homology with natural proteins, they could be structurally similar.

**Figure 1.**
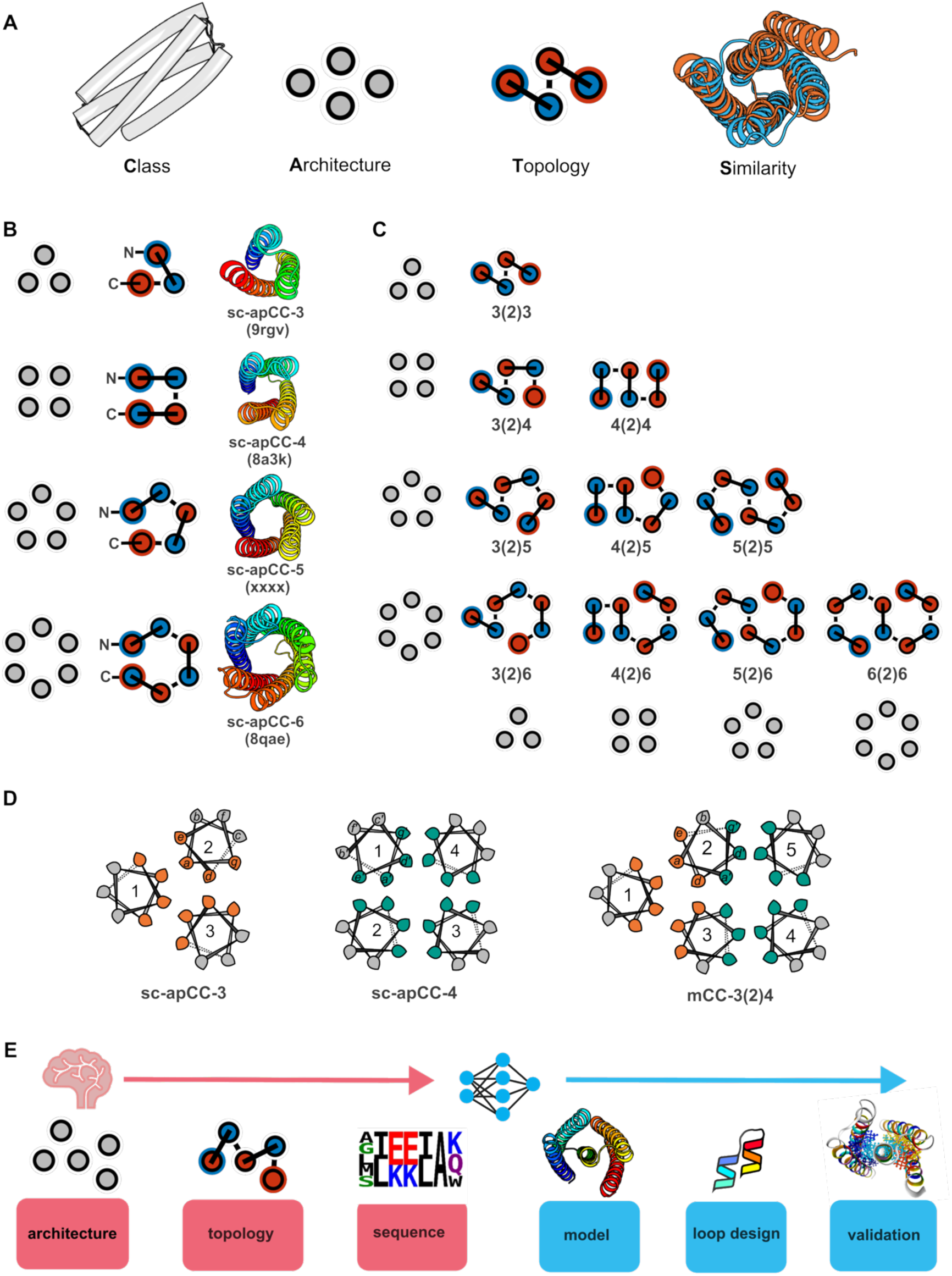
Hierarchical definition and design of multi-coiled-coil proteins (mCCs). (**A**) Designed coiled coils (CCs) can be classified using CATS (class, architecture, topology, similarity) analogous to the CATH system^48^. CCs belong to the all-α class; architecture captures helix number and their arrangement in 3D; topology defines helix orientation and connectivity; and similarity relates the *de novo* proteins to known or predicted natural folds using FoldSeek^56^. (**B**) *De novo* single-chain coiled-coil modules (scCCs)^11,38^ serve as building blocks for mCCs. (**C)** mCCs constructed by lateral blending of two scCC domains *via* bifaceted helices. The resulting architectures are defined by component modules and the number of shared helices, and alternative helix orientations and loop connectivity generate distinct topologies. (**D**) Helical-wheel representations of 3- and 4-helix bundles showing ***gade*** positions of the heptad sequence repeats in orange and green, respectively. The mCC-3(2)4 protein, formed by merging these modules, with shared bifaceted helices showing the residues that contribute to both modules. (**E**) Rational design pipeline supported by AI, progressing from architecture and topology, through sequence assignment and structural modelling, to loop optimization and final model validation. Topology diagrams use a consistent notation throughout the paper: grey discs indicate standalone helices; blue and red central discs mark the N- and C-termini of each helix, respectively; outer blue and red rings show the positions of the N- and C-terminal helices in single chains, respectively; and solid and broken lines indicate helix connection nearest to and furthest from the viewer, respectively.

Previously, using sequences and structures of *de novo* coiled-coil peptide assemblies^29^ as rational seeds for computational design, we have made scCC domains with 3, 4, and 6 antiparallel helices, Figure 1B^11,38,55^. For the present study, we added a mixed parallel/antiparallel 5-helix bundle, sc-apCC-5 (PDB ID: 31WI; Figures 1B and S1). Using these scCCs, we explored if rational seeding could be extended to deliver structures with multiple, merged coiled-coil modules (mCCs). For these, we developed a notation to specify different architectures based on the number of helices in each module and the number of helices shared between them. For example, *3(1)3* denotes two 3-helix bundles (3HBs) sharing one helix; and *3(2)4(2)3* has two terminal 3HBs, each sharing two helices with a central 4HB. Specific topologies can be described by extending this notation; e.g., *-3(1)+3* refers to an *N-*terminal 3HB with an anticlockwise arrangement of helices (-) sharing a helix with another 3HB with a clockwise arrangement (+) (Figure S2)^55^. Visualising this with topology diagrams, illustrates how, even with a small set of scCCs, complexity and diversity can be built up systematically and rapidly, Figure 1A-C. For example, for two-domain mCCs built from modules of 3 to 6 helices and sharing two helices, there are ten distinct architectures, each with various possible unique topologies depending on how the helices are connected (Figure 1C, Figure S2&S3).

Whilst sequence and structural seeds for the separate modules are available for quick sequence and model construction of target mCCs (Table S1)^11,38^, defining sequences for the shared helices is more challenging. Therefore, we took sequences for the regular helices from the relevant seeds to design bifaceted helices^53^ as follows. Heptad sequence repeats, ***gabcdef***, underpin CC folding and assembly, with the ***gade*** positions forming seams that specify helix-helix interactions^29^. To generate bifaceted helices, two sets of ***gade*** residues were offset within a sequence to encode two opposite faces on the resulting helix^57^. For example, in mCC-3(2)4, Figure 1D, helices 2 and 3 of the 3HB and helices 1 and 2 of the 4HB are shared. With the ***gabcdef*** registers of the 3HB helices as the reference, each helix can be made bifaceted by adapting their ***bef*** positions to specify new ***a’d’e’*** residues derived from the 4HB sequence. As a result, each bifaceted helix has two ***gade***s offset by one residue, ***gaa’cdd’e’***, giving two seams separated by ≈100° (Figure 1D, Table S2).

We formalised this in the following pipeline, integrating rational design, AI-based structure prediction, loop templating, and loop-sequence building (Figure 1E). For a user-defined target mCC architecture and topology, sequences for each helix were generated as described above (Figure S2 and Table S2&S3). Then, initial single-chain sequences were generated by joining the helices with Gly-Ser-based linkers. AlphaFold2^6^ (AF2) models for these were used in template-based searches using MASTER^58^ to insert loop backbones from the PDB followed by ProteinMPNN^4^ to generate loop sequences. Structure models were re-predicted with AF2 (without MSAs or templates) and ESMFold^59^ and evaluated based on consistency with the initial target (C_α_-RMSD ≤ 1 Å), confidence metrics (pLDDT ≥ 80, pTM ≥ 0.7), and structural evaluation (KIH packing using a 7 Å cutoff in Socket2^60^).

In this way, the construction of complex mCCs is reduced to rapid rules-based sequence design augmented with generative AI tools.

### *De novo* mCC proteins fold as designed

To validate the design framework experimentally, we generated 31 mCCs spanning diverse architectures (Figures 2 and S4&S5, Table S4). These included two-module constructs built from modules with 3 – 6 helices sharing two or three interfacial helices (Figure 2A), as well as three-module proteins to explore the combinatorial space further (Figure S4). Examples of the latter included mCC-3(2)4(2)3, a modular extension of mCC-3(2)4. In addition, mCC-3(1)2(1)3 and mCC-5(1)2(1)5 had terminal modules connected via a central two-helix module (see Figure S4A, rows 1-3). Notably, for most topologies, a single design was sufficient to obtain highly confident AF2 predictions consistent with the design target (Figures 2B and Table S3).

**Figure 2:**
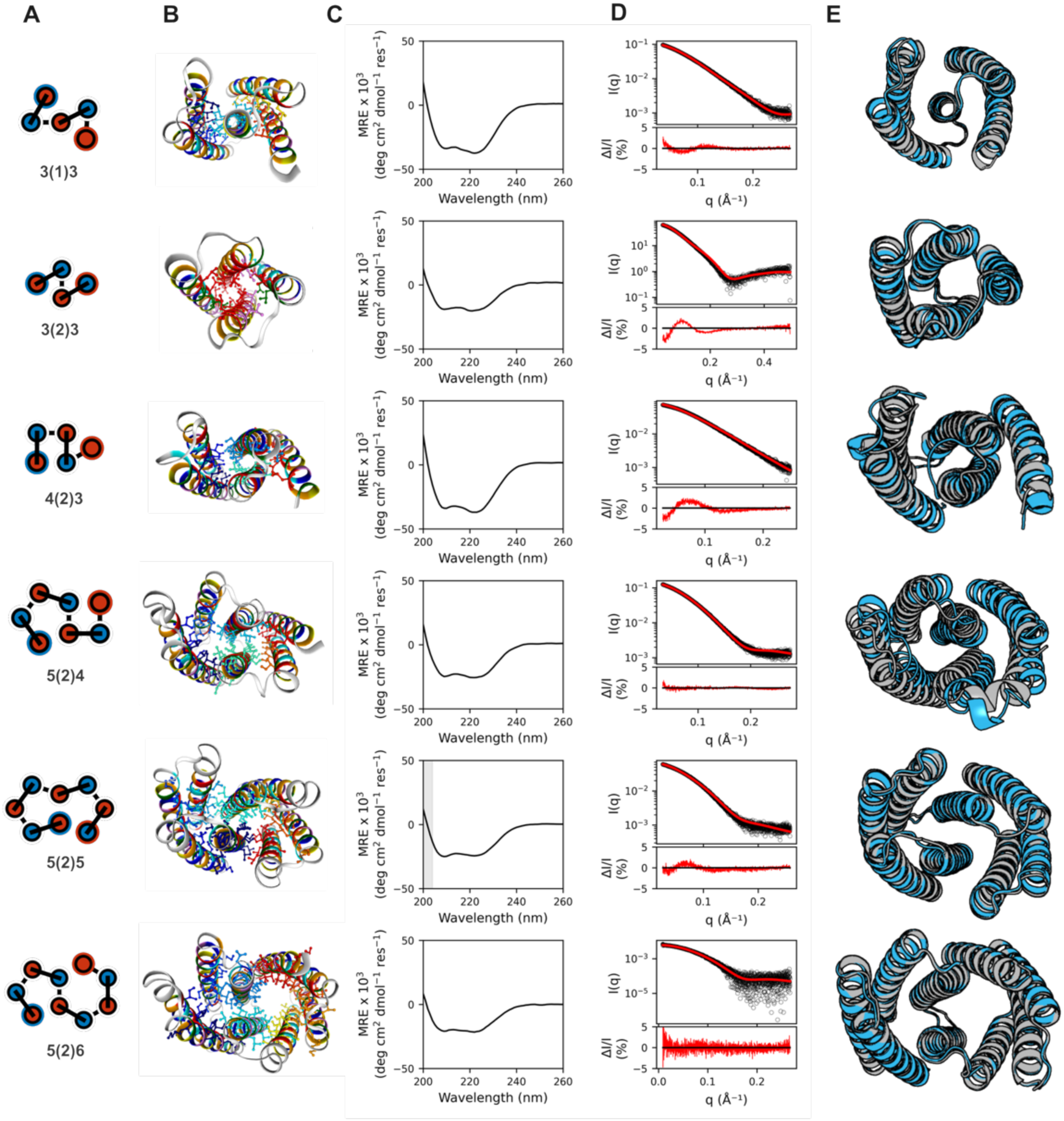
Biophysical characterisation of representative mCCs. (**A**) Topology diagrams of design targets. (**B**) AF2 models of final designs assessed by SOCKET^60^ and coloured according to heptad register assignment with ***abcdefg*** positions shown in red, orange, yellow, green, cyan, blue, and violet, respectively. Core-packing residues identified by SOCKET are shown as balls and sticks. (**C**) Circular dichroism (CD) spectra confirm α-helical structures (MRE, mean residue ellipticity). Conditions: 5 – 10 µM protein in 50 mM sodium phosphate, 150 mM NaCl, pH 7.4, at 5°C. Grey shading indicates regions where the detector high-tension (HT) voltage exceeded 600 V. **(D)** Small-angle X-ray scattering (SAXS) data collected in batch mode, with fits to AF2 models (black, experimental; red, calculated). χ² fits: 2.3, 12.4, 6.7, 1.5, 2.1, and 1.0 from top to the bottom. Conditions: 2.5 – 5 mg ml^−1^ protein in Tris-HCl, 50 mM NaCl, pH 8. **(E)** Superimposition of X-ray crystal structures (cyan) and AF2 models (grey), PDB IDs are 31WJ, 31WO, 31WP, 31WQ, 31WS and 31WT, respectively. Backbone RMSDs are 0.7, 0.6, 1.3, 3.2, 0.7, and 0.5 Å (top to bottom).

Of the 25 designs tested initially, 16 expressed as soluble proteins in *E. coli* and were monodisperse as determined by analytical size exclusion chromatography (SEC) (Figure S3B). Far-UV circular dichroism (CD) spectra confirmed α-helical proteins with high thermal stabilities (Figure 2C, Figure S3C-D). Small-angle X-ray scattering (SAXS) data were consistent with the AF2 models (Figure 2D, Figure S3E). High-resolution X-ray crystal structures were obtained for six different designs and confirmed the targeted architectures and topologies (Figure 2E, Table S5). Five of these were virtually identical to the AF2 predictions for the designs, with backbone RMSD values 0.6 – 1.3 Å. The structure of mCC-5(2)4 also confirmed the overall architecture and topology, but revealed a partially collapsed 5-helix barrel (Figure 2E, row 4). Interestingly, this suggests potential frustration between the helical supercoils of the two modules, which might be exploitable to design conformational switches^61^.

### mCC topology can be controlled precisely

To demonstrate the full programmability of mCCs, we generated the set of all four topologies for mCC-3(1)3 by combining clockwise and anticlockwise 3HB modules (Figure 3A and Figure S2)^55^. We dubbed the topologies S, W, Z and M as they form these shapes when drawn with their N termini on the left. These were specified by sequence-to-structure relationships of surface charged and buried polar interactions that drive different helix-helix interactions^55^. Two single-chain sequences for each topology were designed and expressed from synthetic genes in *E. coli* (Table S3). All proteins were monomeric, mainly α helical, and hyper-thermal stable (Figure S5). High-resolution crystal structures for all four topologies were obtained (Figure 3D). These closely matched the design models, with electron density unambiguously confiming each topology (Figure 3C and Figure S6).

**Figure 3:**
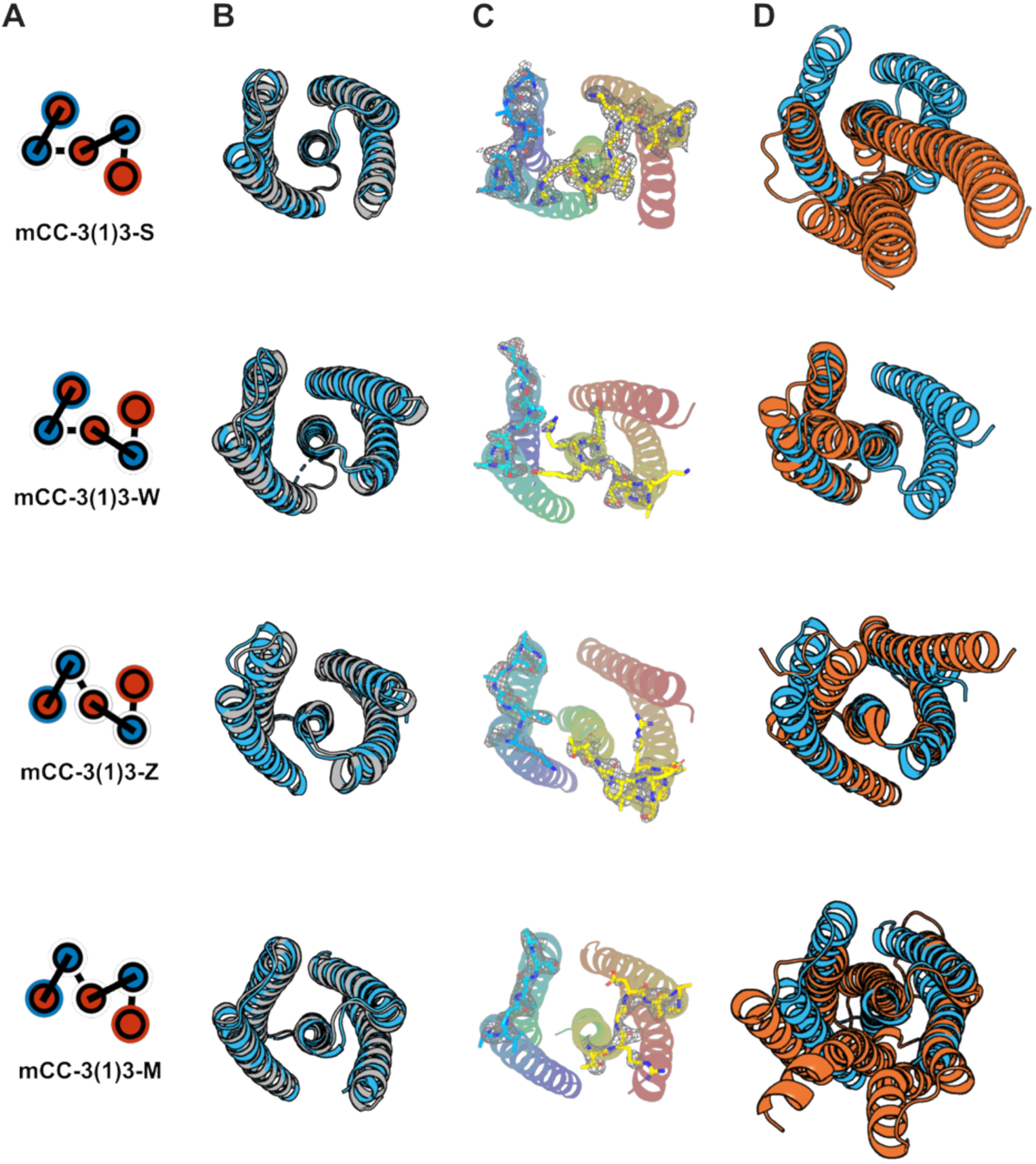
Control over mCC topology generates folds distinct from those found in nature. **(A)** Topology diagrams of the S, W, Z and M variants of the mCC-3(1)3 architecture. **(B)** Backbone superposition of X-ray crystal structures (blue) and corresponding AlphaFold2 models (grey) for the four topologies. PDB IDs are 31WJ, 31WK, 31WL and 31WN, with resolutions of 2.5, 2.2, 2.7 and 2.8 Å, respectively (top to bottom). **(C)** Electron density for the first and third loop regions, coloured by chain (chainbow). Refined loop residues are shown as sticks; maps are contoured at 1.0σ. **(D)** Closest structural matches for each designed mCC identified using a FoldSeek/TM-align workflow. The top-scoring TED domains (orange) are superimposed on the designed mCC structures (blue). TM-scores for the matches (top to bottom) are 0.56, 0.53, 0.76, and 0.66.

### *De novo* mCC proteins are distinct from natural folds

To assess the structural similarity of our designed mCCs to natural proteins, we searched the structural databases with FoldSeek (https://search.foldseek.com)^56^ starting with the experimental structures for the mCC-3(1)3 designs, Figure 3D.

Strikingly, only mCC-3(1)3-Z-1 showed a close but ambiguous match to a known fold, with a TM-score of 0.76 and high coverage for both the target and query (>0.85). mCC-3(1)3-M-1 was borderline: although its overall architecture loosely resembled a known fold, substantial helix shifts, altered packing interactions, and reduced target coverage (0.64) precluded confident assignment to any known protein fold. The two remaining topologies—mCC-3(1)3-S and mCC-3(1)3-W-1—had either low query or target coverage and TM-scores < 0.6 (Figure 3D, Figure S7). Thus, while related α-helical architectures have been accessed, the specific helix connectivities and spatial arrangements in 2 out of 4 of these mCCs seem not to have been sampled by natural evolution. This highlights the tight control afforded by our design strategy over both helix orientation and connectivity, and it underscores the ability of mCCs to explore regions of α-helical fold space beyond those observed in nature.

We extended the searches to all 18 validated mCCs using experimental structures where available and AF2 models otherwise. For this, we applied a multi-step structural similarity workflow combining high-sensitivity FoldSeek searches with high-precision TM-align comparisons. An online FoldSeek search against CATH50 identified hits from 48 CATH superfamilies, indicating that the query mCCs sampled a broad region of structural space. However, most FoldSeek hits occurred near the TM-score cutoff (0.6), consistent with the known tendency of aligned TM-scores to be inflated when only overlapping structural regions are considered. To examine this, we expanded the search to the full TED domain space (2.44 million structures and models) to give exhaustive coverage of the structural diversity within the matched superfamilies.

This second FoldSeek search gave 9,575 candidate mCC:TED pairs, which were evaluated using TM-align for high-accuracy structural comparison. Although FoldSeek identified hits for all 18 tested mCCs, 4 designs could not be successfully superimposed onto the corresponding TED structures, indicating no meaningful global similarity. The remaining 14 pairs were filtered using a ≥70% query coverage threshold, requiring that at least 70% of the query structure was aligned to the target (Figure S5). This revealed a further 11 without any convincing fold-level similarity to the 2.44 million curated TED domains, even under permissive TM-score thresholds. In these cases, any matches were limited to local or fragment correspondences and did not translate into shared global topology.

Thus, 15 out of 18 mCCs queried in the extended search are strong candidates for genuinely new-to-nature folds, expanding the known landscape of helical bundle architectures beyond those observed in nature and represented in CATH.

### mCC proteins as robust scaffolds for dual functionalisation

Finally, we tested how robust mCCs were to mutation and multi-functionalisation by designing functional sites into several scaffolds. We defined three distinct design goals to generate bifunctional proteins for: (1) binding two different natural ligands; (2) creating interaction hubs to engage multiple proteins; and (3) combining synthetic small-molecule and protein-binding modules in a single construct. Our hypothesis was that as each module is specified by multiple sequence-to-structure rules, some positions could be sacrificed to *graft on* or *drop in* constellations of functional residues.

First, we tested whether mCC scaffolds could accommodate distinct and orthogonal ligand-binding sites within separate 4HB modules. As these are proven *de novo* scaffolds for metal and haem binding^62,63^, we introduced a 3-His, Zn²⁺-binding site and a 2-His, haem-binding site into the different 4HB modules of mCC-4(2)4 to give mCC-4(2)4-ZN-HEM. The designed complex predicted with high confidence by AlphaFold3 (Figure S8). The protein expressed well in *E. coli*, and retained the biophysical properties of the parent scaffold, remaining monomeric, α helical and thermally stable (Figure S9). Spectroscopic and competition assays showed tight binding to both Zn²⁺ (sub-µM) and haem (apparent K_D_ ≈ 10 nM, at the limit of the assay) (Figure 4A; Figures S10–S13). Binding affinities and stoichiometries were largely unaffected by the presence of the second ligand, indicating that the Zn²⁺ and haem sites function independently (Figure S10 & S11). Thus, orthogonal metal- and cofactor-binding sites can be installed within different modules of an mCC scaffold.

**Figure 4.**
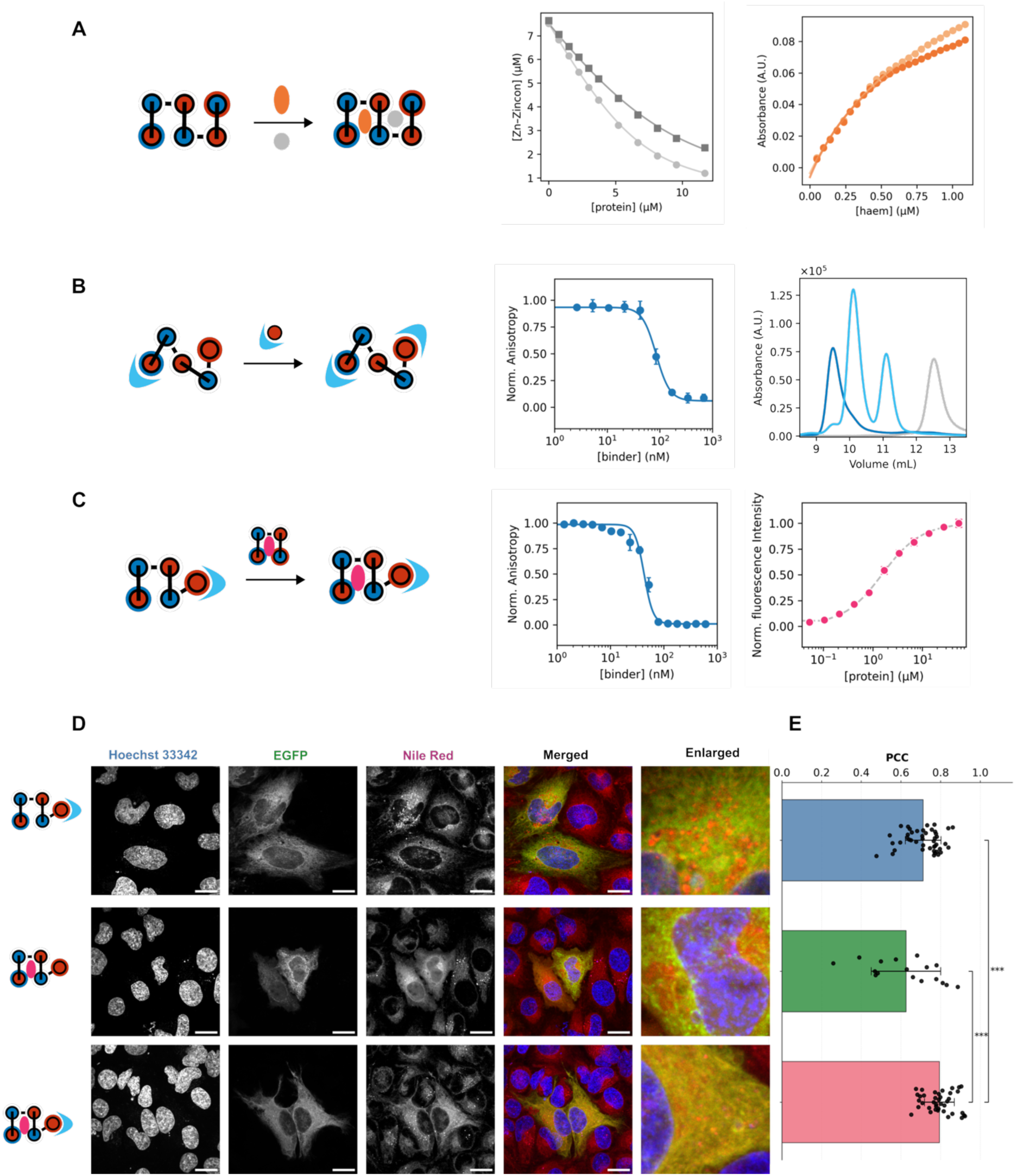
Functionalisation of mCC scaffolds. (**A**) Left: orthogonal binding sites for two ligands, Zn^2+^ and haem, designed into distinct 4HB modules of mCC-4(2)4 to give mCC-4(2)4-ZN-HEM. Middle: Zn^2+^ binding assays show reveal tight binding in the absence (grey) and presence (dark grey) of haem, with K_D_ values of 260 ± 10 nM and 400 ± 50 nM, respectively, and 1.37 ± 0.01 effective binding sites. Right: haem binding assays show tight binding in the absence (light orange) and presence (orange) of Zn^2+^ with K_D_ values of 12 ± 8 nM and 17 ± 10 nM, respectively. (**B**) Left: one or two MCL-1 binding sites grafted onto mCC-3(1)3. Middle: competitive fluorescence anisotropy assay of mCC-3(1)3-Z-1-MCL1-2 (blue) using a fluorescent MCL-1 binding peptide (BID) as tracer, showing its displacement by the designed binder with IC_50_ <100 nM). Right: size-exclusion chromatography of mCC-3(1)3-Z-1-MCL1-2 in the absence of MCL-1 (grey), and in mixtures with MCL-1 at 1:2 (blue) and 2:1 (cyan) molar ratios (total concentration of MCL-1 was kept constant at 30 µM), showing binding-dependent shifts in elution volume. (**C**) Left: mCC-3(2)4 dual-functionalised with MCL-1 and Nile red binding sites. Middle: competitive fluorescence anisotropy assay of mCC-3(2)4-MCL1-2-NR with MCL-1 (blue) with IC_50 <_100 nM. Right: saturation-binding curve measured by fluorescence intensity for Nile red binding (pink), K_D(Nile Red)_ = 1.3 ± 0.1 μM. (**D**) Confocal fluorescence images of live HeLa cells expressing an eGFP-MCL-1 fusion together with mCC-3(2)4-MCL1-2, mCC-3(2)4-NR, and mCC-3(2)4-MCL1-2-NR (top to bottom). Channels from left to right: Hoechst 33342 nuclear stain, eGFP-MCL-1, Nile red, merged signal, and zoomed-in regions. The scale bar and the zoomed-in regions are 20 µm. (**E**) Co-localization efficiency (expressed as Pearson correlation coefficient (PCC)) between Nile red and eGFP in the images from (C). Brackets show pairwise comparisons using two-sided Mann-Whitney U test with Holm correction for multiple comparisons, p < 0.001 (***).

Second, to install multiple protein-binding sites, we selected mCC-3(1)3, which has four highly exposed helices, reasoning that they should be good hosts for binding sites (Figure 4B). For the target natural protein, we chose MCL-1^64^, a well-established cancer target with validated α-helical binding partners^41,64,65^. We grafted one or two copies of such a binding motif onto the various 3(1)3 topologies (Table S8 and Figure S14), and assessed their potential to bind MCL-1 with AF2 (Table S8). We tested three each of single and double binding-site designs experimentally (Figure S14). All six proteins were monomeric and highly stable and, for both the single- and double-binder designs, 2/3 bound MCL-1 with IC_50_ < 500 nM in a competitive fluorescence anisotropy assay (Figure 4B, Figures S14 & S15). A crystal structure of a binder:MCL- 1 complex confirmed the designed binding mode (Figure S16, PDB ID 31WU). Analytical SEC showed that, only the bi-functionalised mCC-3(1)3-Z-2-MCL1-2 bound two copies of MCL-1 with high affinity (Figure 4B, Figure S17).

Third, we designed an mCC protein for intracellular visualisation by introducing binding sites for (i) MCL-1 as a subcellular target, and (ii) a fluorescent reporter dye into mCC-3(2)4 (Figure 4C). For protein binding, we adapted the above approach, using the 3HB module and a grafting pipeline^41^. Reasoning that dye binding would require a larger core, we installed a Nile red-binding site^40^ into the 4HB. We made three single-binder controls and two dual-binder designs (Figure 4C, Table S9 and Figure S18). All 5 constructs expressed in *E. coli* to give monomeric, helical, hyperstable proteins (Figure S18). Consistent with our graft-on/drop-in design pipelines, the controls bound only their intended targets, MCL-1 or Nile red (Table S9, Figure S18), and the bifunctional designs bound both in the low nM and low µM ranges, respectively (Figure 4C, Figure S18).

Next, we tested the best dual binder in eukaryotic cells. HeLa cells were transfected with an eGFP-MCL-1 fusion construct together with either the MCL-1/Nile red dual binder or the monofunctional MCL-1 or Nile red binding controls. The resulting live cells were stained with 0.1 µM of Nile red and imaged for the nucleus, the MCL-1 fusion, and the mCCs (Figure 4D). Consistent with MCL-1 being primarily located at the mitochondrial outer-membrane^66^, mitochondrial networks were visible in the eGFP channel (Figure S19). This signal correlated significantly with that for Nile red with the bi-functional MCL-1/Nile red-binding protein (Figure 4E). However, for the controls, the Nile-red channel showed puncta with the MCL-1-only binder as expected for staining lipid droplets^67^, and a more-diffuse cytoplasmic signal with the Nile-red-only binder consistent with sequestration of the dye; and both had weaker correlations to the eGFP channel (Figure 4E). Thus, the graft-on/drop-in design strategy can quickly render dual-functional mCC proteins of relatively low molecular weight (≈20 kDa) for direct visualisation of sub-cellular proteins.

## Discussion

In summary, we have expanded the structural space of all-α-helical proteins by developing an integrated rational and computational design framework for building multi-coiled-coil proteins (mCCs). Our approach is modular, rapid, scalable, and interpretable. It combines understood sequence-to-structure relationships (design rules) for helix-helix interactions and strategies for connecting helices, with AI-based methods for (i) generating loop sequences to produce single-chain proteins, and (ii) structure prediction and model evaluation. By systematically defining target architectures and topologies inspired by the CATH hierarchy^15,48,54^, we show that complex, multi-helix proteins can be designed hierarchically with high predictability. A key element to this success is the rational introduction of bifaceted helices that bridge two helical-bundle modules by satisfying sequence constraints from each.

The strategy is also robust: we deliver *de novo* proteins spanning 14 different overall architectures and 25 different topologies. In total, 31 designs are tested, 75% of which yield soluble, monomeric, and thermostable proteins, and 30% lead to X-ray crystal structures that confirm the design models. These success rates are extremely high in the current state-of-the-art of protein design. Furthermore, we show that mCC proteins serve as robust scaffolds for single-step functionalisation to introduce protein and small-molecule binding sites. Together, these examples establish mCCs as programmable *de novo* proteins in which architecture, topology, binding-site placement, and bi-functional outputs can be encoded with atomic precision through an interpretable, sequence-based design pipeline that is augmented and accelerated by generative AI methods where needed.

Interestingly—and despite the hierarchical design based on principles inspired by nature—mCCs explore regions of all-α-helical structural space largely unpopulated by natural proteins. Our FoldSeek^56^ analysis reveals that most mCCs do not have single-chain counterparts in either experimental or predicted structural databases, highlighting how the approach accesses folds beyond those shaped by evolution. Nevertheless, like natural multidomain proteins^13,47^, structural and functional complexity in mCCs is achieved by building hierarchically from modules. However, unlike most evolutionarily derived proteins, mCCs share helices between modules. This is done by explicitly superimposing sequence rules for different modules into bifaceted helices, and, effectively, merging the modules. It is difficult to envisage how this might be achieved through the natural processes of gene duplication and fusion. Thus, our approach bridges a conceptual gap between evolutionary modularity and protein design, establishing a framework for building new classes of α-helical proteins. Using mCC proteins, we envisage moving further into the dark matter of protein structures^68,69^, and providing a foundation for designing complex proteins with emergent properties, such as catalysis, allosteric control, custom biosensors, multi-functional molecular machines, and programmable catalysts that expand on nature’s functional repertoire.

## Methods

### Computational tools

AlphaFold2 (AF2) using single-sequence mode was used to generate structural models for *de novo* protein designs. MASTER was used to search for fragments (loops) between adjacent helices using the default MASTER database (see https://grigoryanlab.org/master) and ProteinMPNN was used to optimize the sequences of the MASTER loops. Additional details and scripts used for computational design from AF2 initial models are available in GitHub (https://github.com/woolfson-group/MasterMPNN).

### Structural similarity search using FoldSeek

To evaluate whether *de novo*-designed mCCs correspond to previously known protein folds, we implemented a two-step structural similarity pipeline combining the speed of FoldSeek with the precision of TM-align. For each mCC, we used the experimentally determined structure when available and the AlphaFold2 prediction otherwise. We first performed structural searches using FoldSeek^56^ against the CATH50 reference database^54^. A conservative TM-score threshold of 0.6 was applied, as the FoldSeek web server reports scores based on the aligned region. For all the superfamilies identified in this initial search, we expanded the hits by retrieving additional matching TED domains^70^, which were compiled into a local FoldSeek database to ensure comprehensive coverage of the structural space represented by these superfamilies. The 21 mCCs were queried against this database using FoldSeek locally with default parameters. To obtain high-accuracy measures of structural similarity, all FoldSeek hits were subsequently re-aligned using TM-align^71^. TM-align outputs query-normalized TM-scores together with alignment lengths and coverage, providing an exact estimate of global structural similarity. Only alignments with ≥70% query coverage were retained for downstream interpretation. The top-ranked TM-align hits for each mCC were visually inspected using PyMOL, and representative alignments were prepared for presentation.

### Expression and purification of mCC proteins

All genes were either ordered as clonal plasmids (Twist Bioscience) or DNA strings (GeneArt, IDT DNA) and directly cloned into pET28a vectors, transformed and then expressed in *E. coli* Lemo21-DE3 or BL21 Star-DE3 (New England Biolabs). Flasks containing 1 L of Miller’s Luria Broth–kanamycin–chloramphenicol and 0.5 mM L-rhamnose in case of Lemo21 and containing Miller’s Luria Broth–kanamycin in case of BL21 Star were inoculated with 5 mL of overnight cultures and incubated to an optical density at 600 nm of ∼0.6 at 37 °C with 200 r.p.m. shaking. Expression was induced with 0.5 mM isopropyl-β-D-thiogalactoside, and cultures were incubated at 37 °C overnight with 200 r.p.m. shaking. Following expression, cultures were pelleted and resuspended in 20 mL lysis buffer (50 mM Tris, pH 7.4, 500 mM NaCl, 30 mM imidazole, 1 mg ml^−1^ lysozyme) for 30 min at 37 °C. Resuspended pellets were sonicated using a Biologics Model 3000 Ultrasonic homogenizer with settings at 50% power and 90% pulser (1 pulse per second) for 5 min and then clarified at 25,500 *g* for 30 min. The clarified lysate was heat shocked at 75 °C for 10 min and then cooled on ice for 10 min before reclarifying at 25,500 *g* for 10 min. The expressed proteins were first purified with Ni affinity chromatography at room temperature. Filtered lysate was loaded onto an ÄKTAprime plus (GE, PrimeView 5.31) equipped with a HisTrap HP 5-mL column (Cytiva). His-tagged proteins were eluted using a single step gradient from 0 to 55% buffer B (buffer A consisted of 50 mM Tris, 500 mM NaCl and 30 mM imidazole at pH 7.4; buffer B consisted of 50 mM Tris, 500 mM NaCl and 300 mM imidazole at pH 7.4). Fractions were combined and further purified by SEC using a HiLoad 16/600 Superdex 200-pg size exclusion column (Cytiva) equilibrated in buffer containing 50 mM sodium phosphate and 150 mM NaCl (pH 7.4) at room temperature. Eluted fractions were pooled, concentrated and separated using SDS–PAGE to confirm protein identities.

### MCL-1 purification

The truncated human MCL-1(172-327) was recombinantly expressed and purified from *E. coli* following the protocol described before^65^. Briefly, BL21-Gold E. coli cells were transformed with a pET28a plasmid encoding for His6-tag-SUMO-MCL-1. Bacteria cultures were grown at 37 °C till an OD600 of 0.6 was reached and subsequently protein expression was induced with 250 µM IPTG and the temperature was lowered to 18 °C during overnight incubation. The cells were collected by centrifugation, and the pellet was resuspended in lysis buffer (25 mM Tris, 500 mM NaCl, pH 8.0), protease inhibitor cocktail tablet, lysozyme and DNAse). Lysis was performed by sonication, before being cleared by centrifugation. The fusion protein was isolated by loading the supernatant on a 5 mL His-trap HP column. After four washes with increasing imidazole concentration up to 100 mM imidazole. The protein was eluted with elution buffer (25 mM Tris, 500 mM NaCl, 400 mM imidazole, pH 8.0). The protein was cleaved with His-Ulp1 SUMO protease during overnight dialysis to dialysis buffer (25 mM Tris, 500 mM NaCl, pH 8.0). To isolate the cleaved MCL-1, the dialysed solution was reapplied to the 5 mL His-trap HP column. The flowthrough, containing the protein of interest was collected. A third purification step was performed by size exclusion chromatography using a HiLoad 26/60 Superdex 75 column equilibrated with 20 mM Tris, 250 mM NaCl, 0.5 mM DTT, 2.5% (v/v) glycerol, pH 8.0). The fractions containing MCL-1 were pooled and concentrated to 200 µM before flash freezing for storage at -80°C.

### Circular dichroism spectroscopy

Circular dichroism (CD) data were collected on a JASCO J-810 or J-815 spectropolarimeter fitted with a Peltier temperature controller in the far UV region. Spectra Manager (1.55) was used for data collection. CD spectra were acquired at a 5 or 10 μM protein concentration in 50 mM sodium phosphate and 150 mM NaCl (pH 7.4) at 5 °C. Data were collected in a 1 mm quartz cuvette between wavelengths of 190 nm and 260 nm with the instrument set as follows: band width, 1 nm; data pitch, 1 nm; scanning speed, 100 nm min^−1^; response time, 1 s. Each CD spectrum was obtained by averaging eight scans and subtracting the background signal of the buffer and cuvette. For thermal response experiments, the circular dichroism signal at a 222 nm wavelength was monitored over the temperature range 5–95 °C at a ramping rate of 1 °C min^-1^ with the same settings and protein concentrations given above. The spectra were converted from ellipticities (mdeg) to mean residue ellipticities (deg cm^2^ dmol^−1^ res^−1^) by normalizing for concentration of peptide bonds and the cell path length using the equation (Eq. 1):

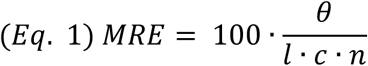

where *θ* is CD ellipticity (degrees), *c* is the molarity of the protein (M), *l* is the path length of the cuvette (cm), and *n* is the number of peptide bonds in the protein.

### Analytical size exclusion chromatography

Analytical size-exclusion chromatography (aSEC) was performed using a Superdex™ 75 Increase 10/300 GL column at a flow rate of 0.8 mL min^-1^. Protein samples (2–5 mg/mL) were prepared in 20 mM sodium phosphate buffer containing 150 mM NaCl (pH 7.4). The elution profiles are reported as max normalized absorbance.

MCL-1 binding experiments were performed as previously described^65^. UV absorbance was monitored at 230 nm using a Superdex™ 75 Increase 10/300 GL column at a flow rate of 0.8 mL min^-1^. Columns and samples were buffered with Tris (20 mM Tris, 150 mM NaCl, pH 7.5) 30 µM of MCL-1 was mixed with variable concentrations of binder proteins to achieve reported ratios.

### Small angle X-ray scattering

SAXS data was collected at the Diamond Light Source beamline B21. Batch mode samples were prepared at 0.5 - 5 mg mL^-1^ in 20 mM Tris pH 8.0, 50 mM NaCl. Size-exclusion chromatography mode samples were prepared at 10-20 mg mL^-1^ in 20 mM Tris pH 8.0, 50 mM NaCl, 1% v/v glycerol. Shodex KW403 column was equilibrated in the same buffer at 4°C. For all designs, buffer subtraction and data merging was performed with ATSAS^72,73^. MultiFoxS software^74^ was used to compare experimental scattering profiles to design models and assess quality of fit by calculating χ^2^.

### X-ray crystallography

Crystals were grown using a sitting-drop vapour-diffusion method. Proteins in 20 mM Tris pH 8.0, 50 mM NaCl were concentrated to 20 mg mL^-1^. Commercially available sparse matrix screens were used (Morpheus^®^, JCSG-plus^TM^, Structure Screen 1 and 2, Pact Premier^TM^, ProPlex^TM^; Molecular Dimensions), and the drops were dispensed using a robot (Oryx8; Douglas Instruments). For each well of an MRC 2 drop plate, 0.3 μL of protein solution and 0.3 μL of reservoir solution, in parallel with 0.4 μL of the protein solution and 0.2 μL of reservoir solution, were mixed and the plate was incubated at 20 °C. Diffraction-quality crystals were grown through multiple rounds of seeding. Crystals generally formed within a month, and after looping were soaked in reservoir solution containing 25% glycerol as a cryoprotectant. Final crystallization conditions for are provided in Table Sx. Diffraction data for the crystals were obtained at the Diamond Light Source (Didcot, UK) on beamlines I03 and I24. Images were processed with Dials^75^, autoPROC-Straniso^76,77^ and Aimless^78^. Structures were phased by molecular replacement with PHASER^79^ using the AF2 model or with *ab initio* phasing using Arcimboldo^80^-Lite. Final structures were obtained after iterative rounds of model building with COOT^81^ and refinement with Phenix Refine^82^. Solvent-exposed atoms lacking map density were deleted. Data merging and refinement statistics for all structures are provided in Table S6.

### Zn^2+^ binding assays

Zn²⁺ binding experiments were performed using freshly purified protein in 50 mM HEPES, 150 mM NaCl, pH 7.0. Before measurements, proteins were treated with EDTA to remove residual bound metals and subsequently buffer-exchanged to remove excess chelator. Stock solutions of ZnCl₂ and Zincon were prepared in water and DMSO, respectively. Zincon concentration was determined from absorbance at 488 nm using ε488nm = 26,900 M⁻¹ cm⁻¹, and ZnCl₂ stocks were standardised by titration against Zincon using ε618nm = 21,200 M⁻¹ cm⁻¹ for the Zn²⁺–Zincon complex at pH 7.0. For affinity measurements, a pre-formed Zn²⁺–Zincon complex was generated by mixing 75 μM Zincon with 10 μM ZnCl₂ and increasing concentrations of protein were added from a concentrated stock. Displacement of Zn²⁺ from Zincon was monitored by the decrease in absorbance at 618 nm using an Agilent Cary 60 UV–Vis spectrophotometer with a 10 mm pathlength cuvette. Apparent K_D_ values and effective numbers of Zn²⁺-binding sites were obtained by fitting the decrease in Zn²⁺–Zincon concentration as a function of protein concentration to a mass-action competition model, as described in the Supplementary Information. Zn²⁺-binding stoichiometry was estimated independently by titrating ZnCl₂ into samples containing 5 μM protein and 10 μM Zincon and monitoring Zn²⁺–Zincon formation at 618 nm. The Zn²⁺ concentration at the transition between the protein-buffered and Zincon-bound regimes was used to estimate the number of high-affinity Zn²⁺-binding sites per protein.

### Haem binding assays

All experiments were performed again with fresh stocks of purified protein with no traceable haem bound to it. The qualitative haem binding ability of the designed variants was determined by measuring UV-Vis spectra of a protein and hemin mixture with 2 μM of protein with 1.5 μM of hemin. Spectra in the 250–600 nm range was recorded using an AgilentCary 60 UV-Vis spectrophotometer in a 10 mm pathlength cuvette and the presence of the characteristic Soret band of haem-binding proteins was detected. Binding affinity and stoichiometry of the designed variants for haem were determined by performing a binding titration and following the absorbance of the formed Soret band at 413 nm. Briefly, a 0.4 μM solution of each protein was prepared in HEPES buffer with 0.5% w/v octyl β-glucoside to minimize aggregation of haem in the aqueous solution. Aliquots of a fresh hemin stock solution prepared in DMSO were added to the protein sample with stirring at 25°C and equilibrated for 10 minutes. The absorbance spectra were then recorded and absorbance at 413 nm was measured. Hemin aliquots were added until a 2.5-fold excess of haem was achieved. Absorbance values at 413 nm were plotted against total haem concentration and data were fitted to a one-site binding equation (Eq. 2) in IgorPro V6.37 to determine the binding affinities.

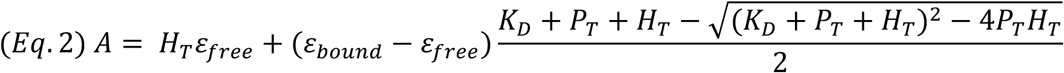

A corresponds to the total absorbance at 413 nm, *H_T_* is the total haem concentration and *P_T_* the total protein concentration. *K_D_* is the dissociation constant, while *ε_free_* and *ε_bound_* correspond to the extinction coefficient of haem when it is free in solution or bound to protein, respectively. The collected spectra corresponding to the titration curves are showed in Supplementary Fig. 12.

### Peptide synthesis of BID tracer

The tracer peptide (FAM-Ahx-EDIIRNIARHLAQVGDS[Nle]DRSIW was synthesised at 0.05 mmol scale following standard Fmoc solid-phase peptide synthesis as described before^65^. Rink amide MBHA resin was used, N,N’-diisopropylcarbodiimide (DIC) and Oxyma Pure in N,N-dimethylformamide (DMF) were used as activators and a 20% (v/v) solution of piperidine in DMF was used as deprotection solution. Each coupling step was followed by two deprotection steps and five washes with DMF. 5-Carboxyfluorescein (FAM) was coupled to the resin bound peptide with a 1:3:4.5:4.5 ratio of resin:FAM:Oxyma:DIC and the resin was subsequently washed twice with DMF. Sidechain deprotection and resin cleavage was achieved with a mixture of trifluoroacetic acid (TFA), triisopropylsilane (TIPS), 2,2’-(ethylenedioxy)diethanethiol (DoDT) and water in a 92.5:2.5:2.5:2.5 ratio. After filtration to remove the resin and evaporation of the TFA, the peptide was precipitate with -20 °C diethyl ether. After centrifugation and decantation of the diethyl ether the crude peptide was freeze dried. The pure peptide was isolated via reverse phase high performance liquid chromatography (RP-HPLC) with a C18 column using a linear gradient of 30% acetonitrile/water to 95% acetonitrile/water. Peptide quality was assessed by HPLC - mass spectroscopy using electrospray ionization.

### Fluorescence anisotropy

Fluorescence anisotropy competition assays were performed in triplicate following a well-established method previous described^65^. Briefly, MCL-1 concentration was fixed at 150 nM and BID tracer (FAM-Ahx-EDIIRNIARHLAQVGDS[Nle]DRSIW-NH_2_) at 25 nM. Measurements were taken after one hour of incubation at room temperature with a CLARIOstar plate reader (BMG Labtech). 6-Carboxyfluorescein (FAM) was excited at 482 nm and emission at 530 nm. The fluorescence anisotropy values were calculated, plotted against binder concentration and fit to the following logistic equation Eq. 2:

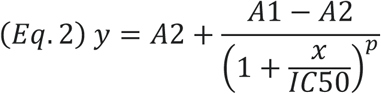

### Nile red binding

Ligand-binding experiments were performed in triplicate. The total concentration of ligand was kept constant: 0.5 μM DPH/Nile in 10% v/v MeCN. The concentration of the protein was varied from 0 – 54 μM. Fluorescence intensity data were collected on a Clariostar plate reader (BMG Labtech) using the following excitation wavelength: λ_DPH_ = 352 nm, λ_Nile red_ = 520 nm. Binding constants were extracted by fitting the data to the following Eq. 3:

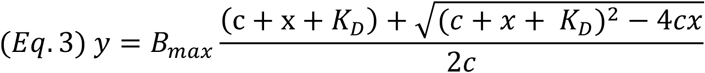

where ***c*** is the total concentration of the constant component (ligand), ***x*** is the concentration of variable component (protein), ***B_max_*** is the fluorescence intensity signal when all the constant component is bound, and ***y*** is the fraction of bound component being monitored via fluorescence signal.

### Transfection and imaging of HeLa cells

The linear DNA fragments for the designer constructs were synthesized by GeneArt or IDT and then subcloned in pTwist CMV vector (Twist Bioscience) for expression in HeLa cells. When required, PCR reactions were carried out using Q5 High Fidelity Hot Start DNA Polymerase (New England Biolabs (NEB)), following manufacturers’ instructions. PCR products were purified using Monarch® PCR & DNA Cleanup Kit (NEB). Restriction digest reactions were carried out using NEB restriction enzymes at 37 °C for 1 h. Ligation reactions were carried out using T4 DNA Ligase and Rapid ligation buffer (Promega) for 1-2 h at 16 °C. Transformation was carried out in competent (subcloning efficiency) E. coli cells (NEB® 5-alpha from NEB or DH5α from Invitrogen) for 30 min on ice, followed by heat shocking (30 s, 42 °C), and a further 5 min on ice. Cells were recovered by incubating in LB media at 37 °C for 1 h and then plated on LB-agar plates containing the ampicillin.

Plasmid DNA samples for transfection were purified using Plasmid Midi Kit according to the manufacturer’s instructions (QIAGEN) and verified by DNA sequencing using Rapid Sequencing Service provided by Source Bioscience.

HeLa cells (ECACC, UK Health Security Agency) were maintained in high glucose Dulbecco’s Modified Eagle’s Medium with 10% (v/v) foetal calf serum (Sigma-Aldrich) and 5% penicillin/streptomycin (PAA) (herein referred to as DMEM) without phenol red at 37 °C and 5% CO2. For transfection, cells were seeded on tissue culture treated 35 mm CELLview cell culture dishes with 10 mm glass bottom (Greiner) at a density of 2×10^5^ cells per well and incubated at 37 °C and 5% CO2 for 16 h prior to transfection. Cells were transfected with the total of 0.6 μg of the indicated plasmid DNA using Effectene transfection reagent according to the manufacturer’s instructions (QIAGEN). After transfection cells were incubated at 37 °C, 5% CO2 for 6 hours, washed with fresh DMEM and incubated further for 18 hours. Before imaging, cells were washed with PBS, stained with Hoechst 33342 (1 μg/mL) and MitoTracker Red FM (Thermo Fisher Scientific, M22425, 0.1 μM) or Nile red (reagent grade, Sigma-Aldrich, 0.1 μM) at 37 °C for 30 minutes, washed again, and then the medium was replaced for fresh DMEM.

Confocal images were collected using an Olympus IXplore SpinSR system with a 60x objective lens at 37 °C using the following imaging channels (λex-max λem/width λem): 405-447/60, 488-525/50, and 561-635/30 nm. Figures were assembled using the Fiji distribution of ImageJ2^83^.

### Image analysis

Co-localisation between Nile red and eGFP signals was quantified using a custom workflow for the ModularImageAnalysis (MIA) plugin for ImageJ^84–86^. The analysis was performed individually for each z-position. The images were first processed with a rolling ball filter (radius = 200 px) to remove background signal. Cells were segmented using a two-step approach using both the nuclear and eGFP component channels. For the cell and nuclear detection the image was intensity normalised and passed through a 2D median filter (radius = 2 px) before candidate cells and nuclei were detected using Cellpose-SAM^87^. Detected nuclei and cells were associated based on spatial overlap and the Pearson correlation coefficient (PCC) was calculated for each cell. Per-cell data was plotted for all samples and the pairwise differences were tested using a two-sided Mann-Whitney U test with Holm correction for multiple comparisons

## Data availability

The coordinate and structure factor files for sc-apCC-5, mCC-3(1)3-S, mCC-3(1)3-W-1, mCC-3(1)3-Z-1, mCC-3(1)3-M-1, mCC-3(2)3, mCC-3(2)4-1, mCC-4(2)5-1, mCC- 5(2)5, mCC-5(2)6-2, and complex of mCC-3(1)3-Z-2-MCL1-1 with MCL-1have been deposited in the PDB with accession codes 31WI, 31WJ, 31WK, 31WL, 31WN, 31WO, 31WP, 31WQ, 31WS, 31WT and 31WU, respectively. Data from this study are openly available in Zenodo at DOI: 10.5281/zenodo.20929603. The code to run the Master-MPNN protocol can be found in GitHub (https://github.com/woolfson-group/MasterMPNN).

## Acknowledgments

K.I.A. and K.W.K were supported by a BBSRC-NSF grant (BB/V004220/1 and 2019598) to O.D.W. and D.N.W. J.J.C. and R.P. were supported by an EPSRC program grant to G.J.L. and D.N.W. (EP/T012455/1). B.M. and A.M.A were supported by a BBSRC grants to D.N.W. and A.J.W. (BB/V006231/1 and BB/V008412/1). K.O. and X.L. were supported by a BBSRC grant to N.S.S. and D.N.W. (BB/X003027/1). R.P. was supported by a BBSRC-funded PhD studentship and by Rosa Biotech through the South West Biosciences Doctoral Training Partnership (BB/T008741/1). A.V.R. was supported by a Leverhulme Trust grant to J.J.M. and D.N.W. (RGP-2021-049) and by the Max Planck-Bristol Centre for Minimal Biology. J.R. was supported by BBSRC grant to C.A.O. (BB/X00306X/1). X. L. was supported by an Engineering and Physical Sciences Research Council (EPSRC)-funded PhD studentship through the Centre for Doctoral Training in Technology Enhanced Chemical Synthesis (EP/S024107/1). SJC and the Wolfson Bioimaging Facility thank the BBSRC for an Alert 19 equipment grant (BB/T017597/1) to purchase the SpinSR.

We thank Freya Spain for performing fluorescence anisotropy assays to test MCL-1 binding for MCL-1 and NR dual binding proteins. We thank the Diamond Light Source for access to beamlines B21, I03 and I24 (proposals mx37593 and mx31440) and The European Synchrotron Radiation Facility for access to beamline BM29 (proposal mx2551).

## Author contributions

D.N.W., K.I.A., and J.J.C conceived the project. J.J.C. and R.P. wrote the computational design code, which was tested by all equally contributing authors. Protein variants and biophysical experiments were designed by K.I.A., J.J.C., L.I.G.R., K.W.K., X.L., B.M., K.O., R.P., A.V.R., and D.N.W.. K.I.A. and K.O. screened for protein solubility and expression, and, with L.I.G.R., purified most of the proteins. Specifically: J.J.C. and L.I.G.R. performed CD experiments; L.I.G.R., X.L., and R.P. conducted most of the SAXS and crystallization trials; L.I.G.R., K.W.K., X.L., B.M., K.O., R.P., and A.V.R. solved the crystal structures; K.O. performed the Nile red-binding experiments; K.W.K. made and characterised the 3(1)3 topologies, with B.M. designing and A.M.A.J. testing the protein binders for these; L.I.G.R. devised, designed and tested small molecule binding; and K.O., J.R. and C.A.O. devised and performed the structural similarity searches. A.V.R. devised and performed cell experiments and analysed the data using programming code for automated image analysis developed by S.J.C. L.I.G.R., K.O., and D.N.W. led the writing of the manuscript, with other equally contributing authors participating in drafting and revision. R.A., J.C., G.J.L., J.J.M., T.A.A.O., C.A.O., N.S.S., A.J.W, A.L.B., and D.N.W. raised funding and/or supervised the above-listed researchers.

## References

1 Kortemme, T. De novo protein design-From new structures to programmable functions. Cell 187, 526–544 (2024). 10.1016/j.cell.2023.12.028

2 Albanese, K. I., Barbe, S., Tagami, S., Woolfson, D. N. & Schiex, T. Computational protein design. Nat Rev Method Prime 5, 13 (2025). 10.1038/s43586-025-00383-1

3 Krishna, R. et al. Generalized biomolecular modeling and design with RoseTTAFold All-Atom. Science 384, eadl2528 (2024). 10.1126/science.adl2528

4 Dauparas, J. et al. Robust deep learning-based protein sequence design using ProteinMPNN. Science 378, 49–55 (2022). 10.1126/science.add2187

5 Dauparas, J. et al. Atomic context-conditioned protein sequence design using LigandMPNN. Nat Methods 22, 717–723 (2025). 10.1038/s41592-025-02626-1

6 Jumper, J. et al. Highly accurate protein structure prediction with AlphaFold. Nature 596, 583–589 (2021). 10.1038/s41586-021-03819-2

7 Abramson, J. et al. Accurate structure prediction of biomolecular interactions with AlphaFold 3. Nature 630, 493–500 (2024). 10.1038/s41586-024-08416-7

8 Sakuma, K. et al. Design of complicated all-α protein structures. Nat Struct Mol Biol 31, 275–282 (2024). 10.1038/s41594-023-01147-9

9 Frank, C. et al. Scalable protein design using optimization in a relaxed sequence space. Science 386, 439–445 (2024). 10.1126/science.adq1741

10 Frank, C., Schiwietz, D., Fuß, L., Ovchinnikov, S. & Dietz, H. Alphafold2 refinement improves designability of large de novo proteins. bioRxiv, 2024.2011.2021.624687 (2024). 10.1101/2024.11.21.624687

11 Albanese, K. I. et al. Rationally seeded computational protein design of α-helical barrels. Nat Chem Biol 20, 991–999 (2024). 10.1038/s41589-024-01642-0

12 Harteveld, Z. et al. Exploring “dark-matter”protein folds using deep learning. Cell Syst 15 (2024). 10.1016/j.cels.2024.09.006

13 Han, J. H., Batey, S., Nickson, A. A., Teichmann, S. A. & Clarke, J. The folding and evolution of multidomain proteins. Nat Rev Mol Cell Bio 8, 319–330 (2007). 10.1038/nrm2144

14 Nepomnyachiy, S., Ben-Tal, N. & Kolodny, R. Global view of the protein universe. P Natl Acad Sci USA 111, 11691–11696 (2014). 10.1073/pnas.1403395111

15 Greene, L. H. et al. The CATH domain structure database: new protocols and classification levels give a more comprehensive resource for exploring evolution. Nucleic Acids Res 35, D291–D297 (2007). 10.1093/nar/gkl959

16 Korendovych, I. V. & DeGrado, W. F. *De novo* protein design, a retrospective. Q Rev Biophys 53, 1–33 (2020). 10.1017/S0033583519000131

17 Woolfson, D. N. A Brief History of Protein Design: Minimal, Rational, and Computational. J Mol Biol 433, 167160 (2021). 10.1016/j.jmb.2021.167160

18 Pan, X. J. & Kortemme, T. Recent advances in protein design: Principles, methods, and applications. J Biol Chem 296, 100558 (2021). 10.1016/j.jbc.2021.100558

19 Baek, M. et al. Accurate prediction of protein structures and interactions using a three-track neural network. Science 373, 871–876 (2021). 10.1126/science.abj8754

20 Butcher, J., et al. *De novo* Design of All-atom Biomolecular Interactions with RFdiffusion3. bioRxiv, 2025.2009.2018.676967 (2025). 10.1101/2025.09.18.676967

21 Watson, J. L. et al. De novo design of protein structure and function with RFdiffusion. Nature 620, 1089–1100 (2023). 10.1038/s41586-023-06415-8

22 Ingraham, J. B. et al. Illuminating protein space with a programmable generative model. Nature 623, 1070–1078 (2023). 10.1038/s41586-023-06728-8

23 Stark, H. et al. BoltzGen: Toward Universal Binder Design. bioRxiv, 2025.2011.2020.689494 (2025). 10.1101/2025.11.20.689494

24 Pacesa, M. et al. One-shot design of functional protein binders with BindCraft. Nature 646, 483–492 (2025). 10.1038/s41586-025-09429-6

25 Lauko, A. et al. Computational design of serine hydrolases. Science 388, eadu2454 (2025). 10.1126/science.adu2454

26 Chu, A. E., Lu, T. Y. & Huang, P. S. Sparks of function by de novo protein design. Nat Biotechnol 42, 203–215 (2024). 10.1038/s41587-024-02133-2

27 Cho, Y., Dauparas, J., Tsuboyama, K., Rocklin, G. J. & Ovchinnikov, S. Stable de novo protein design via joint conformational landscape and sequence optimization. Nat Commun 17, 8 (2025). 10.1038/s41467-025-66526-w

28 Li, Z. & Luo, Y. N. Generalizable and scalable protein stability prediction with rewired protein generative models. Nat Commun 17, 891 (2025). 10.1038/s41467-025-67609-4

29 Woolfson, D. N. Understanding a protein fold: The physics, chemistry, and biology of α-helical coiled coils. J Biol Chem 299, 104579 (2023). 10.1016/j.jbc.2023.104579

30 Kajava, A. V. Tandem repeats in proteins: From sequence to structure. J Struct Biol 179, 279–288 (2012). 10.1016/j.jsb.2011.08.009

31 Parmeggiani, F. & Huang, P. S. Designing repeat proteins: a modular approach to protein design. Curr Opin Struc Biol 45, 116–123 (2017). 10.1016/j.sbi.2017.02.001

32 Schweke, H. et al. An atlas of protein homo-oligomerization across domains of life. Cell 187, 999–1010 (2024). 10.1016/j.cell.2024.01.022

33 Fletcher, J. M. et al. A Basis Set of *de Novo* Coiled-Coil Peptide Oligomers for Rational Protein Design and Synthetic Biology. Acs Synth Biol 1, 240–250 (2012). 10.1021/sb300028q

34 Thomas, F., et al. *De Novo*-Designed α-Helical Barrels as Receptors for Small Molecules. Acs Synth Biol 7, 1808–1816 (2018). 10.1021/acssynbio.8b00225

35 Wood, C. W. & Woolfson, D. N. CCBuilder 2.0: Powerful and accessible coiled-coil modeling. Protein Sci 27, 103–111 (2018). 10.1002/pro.3279

36 Chubb, J. J., Albanese, K. I., Rodger, A. & Woolfson, D. N. De Novo Design of Parallel and Antiparallel A3B_3_ Heterohexameric α-Helical Barrels. Biochemistry-Us 64, 1973–1982 (2025). 10.1021/acs.biochem.4c00584

37 Thomson, A. R. et al. Computational design of water-soluble α-helical barrels. Science 346, 485–488 (2014). 10.1126/science.1257452

38 Naudin, E. A. et al. From peptides to proteins: coiled-coil tetramers to single-chain 4-helix bundles. Chem Sci 13, 11330–11340 (2022). 10.1039/d2sc04479j

39 Gradisar, H. et al. Design of a single-chain polypeptide tetrahedron assembled from coiled-coil segments. Nat Chem Biol 9, 362-+ (2013). 10.1038/nchembio.1248

40 Petrenas, R. et al. Rapid Assessment of Size, Shape, and Chemical Complementarity of Ligands for Computational Protein Design. bioRxiv, 2025.2006.2030.662286 (2025). 10.1101/2025.06.30.662286

41 Mylemans, B., et al. *De novo* designed bifunctional proteins for targeted protein degradation. bioRxiv, 2025.2012.2022.695915 (2025). 10.64898/2025.12.22.695915

42 Scott, A. J. et al. Constructing ion channels from water-soluble α-helical barrels. Nat Chem 13, 643–650 (2021). 10.1038/s41557-021-00688-0

43 Burton, A. J., Thomson, A. R., Dawson, W. M., Brady, R. L. & Woolfson, D. N. Installing hydrolytic activity into a completely *de novo* protein framework. Nat Chem 8, 837–844 (2016). 10.1038/Nchem.2555

44 Petrenas, R. et al. Confinement and Catalysis within Designed Peptide Barrels. J Am Chem Soc 147, 3796–3803 (2025). 10.1021/jacs.4c16633

45 Cross, J. A. et al. A de novo designed coiled coil-based switch regulates the microtubule motor kinesin-1. Nat Chem Biol 20, 916–923 (2024). 10.1038/s41589-024-01640-2

46 Romanyuk, A. V. et al. Biomolecular condensation using *de novo* designed globular proteins. bioRxiv, 2025.2012.2019.695468 (2025). 10.64898/2025.12.19.695468

47 Lees, J. G., Dawson, N. L., Sillitoe, I. & Orengo, C. A. Functional innovation from changes in protein domains and their combinations. Curr Opin Struc Biol 38, 44–52 (2016). 10.1016/j.sbi.2016.05.016

48 Sillitoe, I. et al. CATH: increased structural coverage of functional space. Nucleic Acids Res 49, D266–D273 (2021). 10.1093/nar/gkaa1079

49 Andreeva, A., Kulesha, E., Gough, J. & Murzin, A. G. The SCOP database in 2020: expanded classification of representative family and superfamily domains of known protein structures. Nucleic Acids Res 48, D376–D382 (2020). 10.1093/nar/gkz1064

50 Lo Conte, L., et al. SCOP: a Structural Classification of Proteins database. Nucleic Acids Res 28, 257–259 (2000). DOI 10.1093/nar/28.1.257

51 Hilditch, A. T. et al. Maturation and Conformational Switching of a Designed Phase-Separating Polypeptide. J Am Chem Soc 146, 10240–10245 (2024). 10.1021/jacs.4c00256

52 Ramsak, M. et al. Programmable de novo designed coiled coil-mediated phase separation in mammalian cells. Nat Commun 14, 7973 (2023). 10.1038/s41467-023-43742-w

53 Walshaw, J. & Woolfson, D. N. Extended knobs-into-holes packing in classical and complex coiled-coil assemblies. Journal of Structural Biology 144, 349–361 (2003).

54 Waman, V. P. et al. CATH v4.4: major expansion of CATH by experimental and predicted structural data. Nucleic Acids Res 53, D348–D355 (2024). 10.1093/nar/gkae1087

55 Leng, X. et al. De novo designed 3-helix bundle peptides and proteins with controlled topology and stability. Chem Sci 16, 18632–18641 (2025). 10.1039/D5SC05576H

56 van Kempen, M. et al. Fast and accurate protein structure search with Foldseek. Nat Biotechnol 42, 243–246 (2024). 10.1038/s41587-023-01773-0

57 Walshaw, J. & Woolfson, D. N. Open-and-shut cases in coiled-coil assembly:: α-sheets and α-cylinders. Protein Sci 10, 668–673 (2001). DOI 10.1110/ps.36901

58 Zhou, J. F. & Grigoryan, G. Rapid search for tertiary fragments reveals protein sequence-structure relationships. Protein Sci 24, 508–524 (2015). 10.1002/pro.2610

59 Lin, Z. M. et al. Evolutionary-scale prediction of atomic-level protein structure with a language model. Science 379, 1123–1130 (2023). 10.1126/science.ade2574

60 Walshaw, J. & Woolfson, D. N. SOCKET: A program for identifying and analysing coiled-coil motifs within protein structures. J Mol Biol 307, 1427–1450 (2001). 10.1006/jmbi.2001.4545

61 Dawson, W. M. et al. Structural resolution of switchable states of a de novo peptide assembly. Nat Commun 12, 1530 (2021). 10.1038/s41467-021-21851-8

62 Regan, L. & Clarke, N. D. A Tetrahedral Zinc(Ii)-Binding Site Introduced into a Designed Protein. Biochemistry-Us 29, 10878–10883 (1990). DOI 10.1021/bi00501a003

63 Zhang, S. Q. et al. Design of Tetranuclear Transition Metal Clusters Stabilized by Hydrogen-Bonded Networks in Helical Bundles. J Am Chem Soc 140, 1294–1304 (2018). 10.1021/jacs.7b08261

64 Fletcher, J. M. et al. Coiled-coil peptides as scaffolds for disrupting protein-protein interactions. Chem Sci 9, 7656–7665 (2018). 10.1039/c8sc02643b

65 Acevedo-Jake, A. M. et al. Grafted Coiled-Coil Peptides as Multivalent Scaffolds for Protein Recognition. Acs Chem Biol 20, 1309–1318 (2025). 10.1021/acschembio.5c00137

66 Perciavalle, R. M. et al. Anti-apoptotic MCL-1 localizes to the mitochondrial matrix and couples mitochondrial fusion to respiration. Nat Cell Biol 14, 575-+ (2012). 10.1038/ncb2488

67 Greenspan, P., Mayer, E. P. & Fowler, S. D. Nile Red - a Selective Fluorescent Stain for Intracellular Lipid Droplets. J Cell Biol 100, 965–973 (1985). DOI 10.1083/jcb.100.3.965

68 Taylor, W. R. Exploring Protein Fold Space. Biomolecules 10, 193 (2020). 10.3390/biom10020193

69 Taylor, W. R., Chelliah, V., Hollup, S. M., MacDonald, J. T. & Jonassen, I. Probing the “Dark Matter” of Protein Fold Space. Structure 17, 1244–1252 (2009). 10.1016/j.str.2009.07.012

70 Lau, A. M. et al. Exploring structural diversity across the protein universe with The Encyclopedia of Domains. Science 386, eadq4946 (2024). 10.1126/science.adq4946

71 Zhang, Y. & Skolnick, J. TM-align: a protein structure alignment algorithm based on the TM-score. Nucleic Acids Res 33, 2302–2309 (2005). 10.1093/nar/gki524

72 Franke, D., Gräwert, T. & Svergun, D. I. New features in ATSAS-4, a program suite for smallangle scattering data analysis. J Appl Crystallogr 58, 1027–1033 (2025). 10.1107/S1600576725002481

73 Panjkovich, A. et al. ATSAS 2.8 software for small-angle scattering from macromolecular solutions. Acta Crystallogr A 73, C647–C647 (2017). Doi 10.1107/S2053273317089264

74 Schneidman-Duhovny, D., Hammel, M., Tainer, J. A. & Sali, A. FoXS, FoXSDock and MultiFoXS: Single-state and multi-state structural modeling of proteins and their complexes based on SAXS profiles. Nucleic Acids Res 44, W424–W429 (2016). 10.1093/nar/gkw389

75 Winter, G. et al. DIALS: implementation and evaluation of a new integration package. Acta Crystallogr D Struct Biol 74, 85–97 (2018). 10.1107/S2059798317017235

76 Vonrhein, C. et al. Data processing and analysis with the autoPROC toolbox. Acta Crystallogr D Biol Crystallogr 67, 293–302 (2011). 10.1107/S0907444911007773

77 Vonrhein, C. et al. Advances in automated data analysis and processing within autoPROC, combined with improved characterisation, mitigation and visualisation of the anisotropy of diffraction limits using STARANISO. Acta Crystallogr A: Found Adv 74, a360–a360 (2018). 10.1107/S010876731809640X

78 Evans, P. Scaling and assessment of data quality. Acta Crystallogr D Biol Crystallogr 62, 72–82 (2006). 10.1107/S0907444905036693

79 Mccoy, A. J. et al. Phaser crystallographic software. J Appl Crystallogr 40, 658–674 (2007). 10.1107/S0021889807021206

80 Rodriguez, D. D. et al. Crystallographic ab initio protein structure solution below atomic resolution. Nat Methods 6, 651–653 (2009). 10.1038/nmeth.1365

81 Emsley, P. & Cowtan, K. Coot: model-building tools for molecular graphics. Acta Crystallogr D 60, 2126–2132 (2004). 10.1107/S0907444904019158

82 Afonine, P. V. et al. Towards automated crystallographic structure refinement with phenix.refine. Acta Crystallogr D 68, 352–367 (2012). 10.1107/S0907444912001308

83 Schindelin, J., et al. Fiji: an open-source platform for biological-image analysis. Nat Methods 9, 676-682 (2012). 10.1038/Nmeth.2019

84 Schindelin, J., et al. Fiji: an open-source platform for biological-image analysis. Nat Methods 9, 676-682 (2012). 10.1038/nmeth.2019

85 Schneider, C. A., Rasband, W. S. & Eliceiri, K. W. NIH Image to ImageJ: 25 years of image analysis. Nat Methods 9, 671–675 (2012). 10.1038/nmeth.2089

86 Cross, S. J., Fisher, J. D. J. R. & Jepson, M. A. ModularImageAnalysis (MIA): Assembly of modularised image and object analysis workflows in ImageJ. Journal of Microscopy 296, 173–183 (2024). 10.1111/jmi.13227

87 Pachitariu, M., Rariden, M. & Stringer, C. (bioRxiv, 2025).

