## Supplementary Information for "Expanding all-α-helical protein space through rational computational design"

#### Sequence design of sc-apCC-5

sc-apCC-5 is a chimeric sequence derived from other CC assemblies. Structurally, it comprises four antiparallel interfaces (occupying two ***e–e*** and two ***g–g*** positions of heptad repeat) and one parallel ***e–g*** interface. The ***e–e*** interfaces were retained as Ala–Ala, consistent with other antiparallel ***e–e*** interfaces[1]. For the longer ***g–g*** interfaces, one was adopted from sc-apCC-4 (Gln–Gln) and the other from sc-apCC-6 (Leu–Leu), with the goal of combining topological features from both assemblies[2]. The parallel ***e***–***g*** interface was derived from CC-Pent2 (Ala–Thr)[3]. The core-facing ***a*** and ***d*** positions were maintained as Leu and Ile, respectively, as observed in other α-helical barrels[4] (Table S1). Loop regions were designed in line with the other designs in this study.

#### Sequence design of shared helices

In the mCC architectures sharing one helix, the packing interfaces had a helical offset of *-2* (or *+2*). In the first case, ***b***, ***c***, ***f***, ***g*** (or ***e***, ***f***, ***b***, ***c***) heptad positions of the first domain align with ***g'***, ***a'***, ***d'***, ***e'*** position of the second domain. The residues at these positions are therefore substituted according to the design rules of the second domain (Table S1 and S3). In the architectures sharing two helices, the domains can be merged via the longer ***g–g*** or shorter ***e-e*** interface of the first domain, producing a helical offset of *+1* or *-1*. In the first case, the ***g***, ***c***, ***f*** position of the first domain corresponds to ***a'***, ***d'***, ***g'*** positions of the second domain. In the second case the ***b***, ***e***, ***f*** positions of the first domain correspond to ***a'***, ***d'***, ***e'*** positions of the second domain. In both cases, these positions are substituted according to the design rules of that domain. Unless otherwise specified, all remaining positions retain their identities from the first domain.

#### Loop design

Loop design was carried out in three main steps: (1) defining the order in which helices were connected, (2) initial structural modelling using flexible loops, and (3) loop refinement and filtering. Helix connectivity was constrained to produce continuous looping, such that only neighbouring helices were connected. In addition, all helices belonging to the N-terminal module were required to appear first in the design topology, followed by the helices of the second module. Helical sequences (Table S1 and S3), each consisting of approximately four heptad repeats (***g***–***g*** positions), were connected in the defined order using short, flexible linkers (primarily GSGS and related sequences). The resulting full-length sequences were modelled using AlphaFold2. Models consistent with the intended design topology were then subjected to loop refinement. Loop refinement was performed by redesigning loop regions using a combined approach. Loop backbone conformations were identified using MASTER[5] (input_length = 9, rmsd_cutoff = 0.7, extension_length = 2, max_backbones = 8), followed by loop sequence optimisation using ProteinMPNN[6] (three sequences per backbone). Candidate designs selected for experimental validation were required to pass stringent filtering criteria: maximum RMSD ≤ 1 Å, mean pLDDT ≥ 80, and pTM ≥ 0.7.

#### Introducing topology control in 3(1)3 architecture

The sc-apCC-3[7] consists primarily of leucine residues in the core ***a*** and ***d*** positions of the heptad repeat with the exception of a central polar layer consisting of Asn in the ***a*** position of the third heptad repeat of the N-terminal helix, Thr in the ***d*** position of the second heptad repeat of the second helix, and another Thr in the ***a*** position of the third heptad repeat of the C-terminal helix[7]. In addition to the polar layer, the topology of this 3-helix bundle is driven by salt bridge interactions at the ***e*** and ***g*** heptad positions. In the basic N-terminal helix **B**, these positions are occupied by Lys, while in the acidic helix **A**, they are occupied by Glu, and the C-terminal neutral helix **N** consists of acidic residues at the ***e*** and basic at the ***g*** heptad positions.

#### Double small molecule binding design

The designed protein scaffold mCC-4(2)4 consisting of two 4-helix bundles merged by sharing two bifaceted helices was used as a starting point for the design of two orthogonal binding sites for $Zn^{2+}$ and haem, one in each 4-helix bundle module.. For the design of the $Zn^{2+}$-binding site, Metal-Installer[8] was used to identify positions in mCC-4(2)4 compatible with a three-histidine coordination motif. An AlphaFold3 model of mCC-4(2)4 was used as the input structure, and candidate sites were selected using a Metal-Installer threshold value of 3. From the resulting predictions, sites located near either end of the targeted 4-helix-bundle module were prioritised, with the aim of improving accessibility to the metal centre within the bundle. A reduced set of shortlisted designs was then evaluated by AlphaFold3, and the final candidate was selected based on the confidence scores of the predictions, together with visual inspection of the models to confirm that the predicted $Zn^{2+}$ coordination geometry was structurally plausible. Some additional mutations were introduced in the loop near the designed binding site to ensure that no other side chains could interfere or coordinate the $Zn^{2+}$ ion and to increase accessibility to the binding site (H30S, H32S, D99G, L167G, P171S and E175S). For the design of the haem-binding site, a de novo designed photosynthetic reaction-centre maquette based on a 4-helix-bundle protein was used as a starting template[9]. The target mCC-4(2)4 scaffold was structurally aligned with the maquette to superimpose the first 4-helix-bundle module of the mCC scaffold onto the haem-binding bundle of the reference design. Residues in the maquette that had been designed either to coordinate the porphyrin or to create space for haem binding were then identified, and the corresponding structurally equivalent positions in mCC-4(2)4 were selected. These residues were mutated to recapitulate a similar haem-binding environment within the mCC scaffold. The haem- and $Zn^{2+}$-binding sites were then grafted onto the starting scaffold to generate a dual small-molecule-binding variant, named mCC-4(2)4-ZN-HEM.

#### $Zn^{2+}$ binding affinity determination

$Zn^{2+}$-binding affinities were estimated from Zincon competition assays by fitting the experimentally determined concentration of the $Zn^{2+}$–Zincon complex as a function of total protein concentration after each protein addition. Corrected absorbance values at 618 nm were converted to $Zn^{2+}$–Zincon concentrations using the molar absorption coefficient of the $Zn^{2+}$–Zincon complex under the assay conditions. Total $Zn^{2+}$, Zincon and protein concentrations were corrected for dilution after each addition. The data was then fitted as a function of protein concentration to a mass-action competition model.

The model included two competing equilibria:

$$Zn^{2+} + Zincon \rightleftharpoons Zn\text{-}Zincon$$

$$Zn^{2+} + Protein \rightleftharpoons Protein\text{-}Zn$$

The dissociation constant of Zn–Zincon was fixed to the literature value at pH 7.0:

$$K_d(Zn\text{-}Zincon) = 15.49\ \mu M$$

At each titration point, total Zn was described by:

$$[Zn]_{total} = [Zn]_{free} + [Zn\text{-}Zincon] + [Protein\text{-}Zn]$$

where:

$$[Zn\text{-}Zincon] = \frac{[Zincon]_{total}[Zn]_{free}}{K_d(Zn\text{-}Zincon) + [Zn]_{free}}$$

and:

$$[Protein\text{-}Zn] = \frac{n_{effective}[Protein]_{total}[Zn]_{free}}{K_d(Protein\text{-}Zn) + [Zn]_{free}}$$

For each protein concentration, $[Zn]_{free}$was solved numerically from:

$$[Zn]_{total} = [Zn]_{free} + \frac{[Zincon]_{total}[Zn]_{free}}{K_d(Zn\text{-}Zincon) + [Zn]_{free}} + \frac{n_{effective}[Protein]_{total}[Zn]_{free}}{K_d(Protein\text{-}Zn) + [Zn]_{free}}$$

The predicted Zn–Zincon concentration was then calculated as:

$$[Zn\text{-}Zincon]_{predicted} = \frac{[Zincon]_{total}[Zn]_{free}}{K_d(Zn\text{-}Zincon) + [Zn]_{free}}$$

and fitted to the experimentally determined Zn–Zincon concentrations. The fitted parameters were $K_d(Protein\text{-}Zn)$and $n_{effective}$. For numerical stability, the affinity was fitted as $\log_{10}(K_d)$, and the final dissociation constant was calculated as:

$$K_d(Protein\text{-}Zn) = 10^{\log_{10}(K_d(Protein\text{-}Zn))}$$

The resulting values are reported as apparent Zn-binding dissociation constants under the assay conditions. The effective capacity parameter was included to account for deviations from ideal one-site behaviour and is reported separately from the fitted affinity.

#### MCL-1 binder design protein interaction hubs

In a previous study, we identified a binding site for MCL-1 that can be grafted onto parallel coiled-coil dimer, trimers and tetramers[10,11]. The binding site corresponds to the sequence (L..EQ..LR.IGD.VN..Q..LN), where dots represent any amino acid. This sequence was systematically overlaid onto all exposed helices of the 3(1)3 designs, aligning the ***a*** and ***d*** positions of the coiled-coil register with the placeholder positions (dots). To generate double binders, two such binding sites were placed on distinct coiled-coil domains. Structural predictions for both the binder monomer and its complex with MCL-1 were carried out using AlphaFold2, employing an MSA for MCL-1 and a single sequence input for the binder proteins. Predicted models were filtered to retain only those with pLDDT > 85 for both the monomer and the complex. The remaining designs were then ranked according to the 5-model average pAE of the binder. The pAE and ipTM scores are shown in the Table S6. While the ipTM wasn't used in the selection, it has since emerged as one of the best predictors. The script used to run and analyse the AlphaFold2 predictions can be found on github.

#### MCL-1:Nile Red binder design for MCL-1 visualisation

We applied a previously described AI enhanced motif-grafting method[10]. Briefly, a selection of nature-inspired 6-residue binding motifs, were grafted at all possible locations of the exposed 3-helix bundle module. The resulting models were aligned onto MCL-1:effector complexes. Based on the resulting complexes, non-core residues on the 3(2)4 designs within 6 Å of MCL-1 were altered by a ProteinMPNN-RosettaRelax pipeline. The complex of the final sequence and MCL-1 was predicted as described above. Final selection was based on highest ipTM scores. The script used for the design can be found on github. The Nile Red binding site was subsequently introduced by direct transplantation of binding site residues from the best Nile Red binder designed in single 4HB previously (sc-apCC-4-SN38-1)[12].

### Supplementary Data

**Table S1. Sequence design rules for CC used in this study**

| | register position | | | | | | |
|---|---|---|---|---|---|---|---|
| number of helices | ***a*** | ***b*** | ***c*** | ***d*** | ***e*** | ***f*** | ***g*** |
| **2** | I | A | A | L | K/E | Q(W) | K/E |
| **3** | L | A | A | L | K/E | Q(W) | K/E |
| **4** | L | K/E | K/E | I | A | Q(W) | Q |
| **5** (parallel) | L | K | E | I | A | Q(W) | T |
| **6** (LLIA) | L | K/E | K/E | I | A | Q(W) | L |
| **6** (SLLA) | L | K/E | K/E | L | A | Q(W) | S |

**Table S2. Sequence and computational selection filters for *de novo* designed sc-apCC-5. pLDDT and pTM averaged over five AF2 models.**

| name | architecture | topology | sequence | pLDDT | pTM |
|---|---|---|---|---|---|
| **sc-apCC-5** | 5 | +5 | MMLEEIAQTLEEIAKTLEKIAQTLKKIAQQAEDGEIA QQLEEIAEQLEKIAWQLKKIAQQLKDNAPGEEIAQQL EEIAKQLEKIAQQLKKIAKQTEDGEIAQLLEEIAELL EKIAYLLKKIAQLLKQEKVDGKEIAQLLKEIAKLLEK IAQLLEKIAQLL | 94.9 | 0.89 |

**Table S3. Exemplary sequences of bifaceted helices, x represents solvent exposed residues**

| architecture | **domain 1** ***abcdefg*** | **domain 2** ***abcdefg*** | **bifaceted helix** ***abcdefg*** |
|---|---|---|---|
| 3(1)3 | LxxLxxx | LxxLxxx | LLxLxLx |
| 3(2)3 | LxxLxxx | LxxLxxx | LxLLxxL |
| 3(1)2(1)3 | LxxLxxx | IxxLxxx | LLxLxIx |
| 3(2)4 | LxxLxxx | LxxIAxQ | LLxLIAx |
| 3(2)6 | LxxLxxx | LxxIAxL | LLxLIAx |
| 4(1)4 | LxxIAxQ | LxxIAxQ | LIALALQ |
| 4(2)4 | LxxIAxQ | LxxIAxQ | LLxIIAQ |
| 4(2)5 | LxxIAxQ | LxxIAxQ | LLxIIAQ |
| 4(2)6 | LxxIAxQ | LxxIAxL | LLxIIAQ |
| 5(2)5 | LxxIAxQ | LxxIAxQ | LLxIIAQ |
| 5(1)2(1)5 | LxxIAxQ | IxxLxxx | LLxIIAQ |
| 5(2)6 | LxxIAxQ | LxxIAxS | LLxIIAS |
| 6(2)6 | LxxIAxS | LxxIAxS | LLxIIAS |

**Table S4. Protein sequences passing computational selection filters and chosen for experimental validation. pLDDT and pTM averaged over five AF2 models.**

| name | architecture | topology | sequence | pLDDT | pTM_ |
|---|---|---|---|---|---|
| **mCC-3(1)3-S** | 3(1)3 | -3(1)+3 | MGSSHHHHHHSSGLVPRGSHMMELAALEEELAALEWETAALEEELAALENGSPSAAGLAALKEKLAATKEKLAALKEKLAALKQKQPNSLGLKLLELKLKLLENKNKLLELKLKLLELQRRVGRTPGLAALEEKLAALEEKTAALEEKLAALEASYDPAGLAALEEELAATEYELAALEEELAALEEG | 92.5 | 0.85 |
| **mCC-3(1)3-W-1** | 3(1)3 | -3(1)-3 | MGSSHHHHHHSSGLVPRGSHMMKLAALKEKLAALKWKNAALKEKLAALKNGTPSDAGLAALEEELAATEEELAALEEELAALEKEQPGNVGLELLKLELELLKNETELLKLELELLKAERYGDKAGLAALKEKLAALKEKTAALKEKLAALKQKQGASAGLAALEEKLAATEYKLAALEEKLAALEEG | 92.0 | 0.85 |
| **mCC-3(1)3-W-2** | 3(1)3 | -3(1)-3 | MGSSHHHHHHSSGLVPRGSHMMELAALEEELAALEWENAALEEELAALEEAAEEGEDVSEGLAALKEKLAATKEKLAALKEKLAALKEKQPDSVGLKLLELKLKLLENKTKLLELKLKLLELKYYGDEEEKEKGLAALEEELAALEEETAALEEELAALEKEQPNSPGLAALKEELAATKYELAALKEELAALKEG | 93.0 | 0.85 |
| **mCC-3(1)3-Z-1** | 3(1)3 | +3(1)-3 | MGSSHHHHHHSSGLVPRGSHMMKLAALKEKLAALKWKNAALKEKLAALKKKLGASAGLAALEEELAATEEELAALEEELAALEKAGLSEKEKELGLKLLELKLKLLELKTKNLELKLKLLELKTAPESVRVPGLAALEEELAATEEELAALEEELAALEKEVPEKYKAGLAALKEELAALKYETAALKEELAALKEG | 92.0 | 0.84 |
| **mCC-3(1)3-Z-2** | 3(1)3 | +3(1)-3 | MGSSHHHHHHSSGLVPRGSHMMKLAALKEKLAALKWKNAALKEKLAALKKKLGASAGLAALEEELAATEEELAALEEELAALEKAGLSEKEKELGLKLLELKLKLLETKTKLLELKLKLLELKTAPESVRVPGLAALEEELAALEEETAALEEELAALEKEVPEKYKAGLAALKEELAANKYELAALKEELAALKEG | 92.2 | 0.84 |
| **mCC-3(1)3-M-1** | 3(1)3 | +3(1)+3 | MGSSHHHHHHSSGLVPRGSHMMKLAALKEKLAALKWKTAALKEKLAALKNVPLTPEEKAGLAALEEELAATEEELAALEEELAALEEVSGDPAERERGLKLLELKLKLLETKNKLLELKLKLLELRLTRGDSEGLAALEEKLAALEEKTAALEEKLAALERAGLSDEEAAAGLAALEEELAANEYELAALEEELAALEEG | 91.0 | 0.84 |
| **mCC-3(1)3-M-2** | 3(1)3 | +3(1)+3 | MGSSHHHHHHSSGLVPRGSHMMKLAALKEKLAALKWKNAALKEKLAALKNVPLTPEEKAGLAALEEELAATEEELAALEEELAALEEVSGDPAERERGLKLLELKLKLLELKTKNLELKLKLLELRLTRGDSEGLAALEEKLAATEEKLAALEEKLAALERAGLSDEEAAAGLAALEEELAALEYETAALEEELAALEEG | 91.3 | 0.85 |
| **mCC-3(2)3** | 3(2)3 | +3(2)+3 | MGSSHHHHHHSSGLVPRGSHMMLAALKEKLAALKEKNAALKYKLAALKKKKGKSKGLAKLEELLAKTEETLAELEWLLAEQEKNPDAAGLAELEETLAELEELTAKLEELLAKLEAEELDGAALEEELAALEEELAANKKKLAALK | 92.5 | 0.82 |
| **mCC-3(1)2(1)3** | 3(1)2(1)3 | +3(1)2(1)+3 | MGSSHHHHHHSSGENLYFQSGSLAALKEKLAALKEKNAALKYKLAALKEKKPGNPEGAALEKELAATEKELAALEWELAALESGEKSSKKNGELLELKLEILELKTELLELKLEILELDPSLPELERRGRLLKLKLRILKLKNRLLKLKLRILKLETGWTPEGAALEKELAATEKELAALEWELAALENGDPTAEQGAALEEKLAALEEKTAALEYKLAALEAD | 91.3 | 0.86 |
| **mCC-3(2)4-1** | 3(2)4 | -4(2)-3 | MGSSHHHHHHSSGLVPRGSHMMQLEEIAQQLEEIAEQLKKIAEQLKKIAKGHPNGKGLEEIAQQLEEIAEQLKKIAWQLKKIATAPEEERVPGLLEIAEQLLEIAEQLTKIAEQLLKIAEQGNDKGLLEIAKQLTEIAKQLLKIAKQLLKIAREGGVDEEGLAALEKKLAALEQKNAALEYKLAALE | 93.4 | 0.88 |
| **mCC-3(2)4-2** | 3(2)4 | +4(2)-3 | MGSSHHHHHHSSGLVPRGSHMMQLLEIAQQLLEIAEQLTKIAEQLLKISVEEPEDSEGLEEIAQQLEEIAEQLKKIAWQLKKIAEGHPHGKGLEEIAQQLEEIAKQLKKIAEQLKKIATAPPEVRSPGLLEIAQQLTEIAKQLLKIAEQLLKIAKEEGVDEEGLAALEKKLAALEKKNAALEYKLAALE | 91.7 | 0.86 |
| **mCC-3(2)4(2)3** | 3(2)4(2)3 | +3(2)+4(2)-3 | MGSSHHHHHHSSGENLYFQSGSLAALEKKLAALEQKNAALEQKLAALEKEGKQSGLLEIAKQLTEIAKQLLKIAKQLLKIAKGKGDAGLLEIAEQLLEIAEQLTKIAEQLLKIGVEKPDDYEGLLEIAEQLLEIAEQLTKIAEQ | 93.2 | 0.88 |

| | | | | | |
|---|---|---|---|---|---|
| | | | LLKIAEGKGDKGLLEIAKQLTEIAKQLLKIAWQLLK<br>IAKENGVSGAALEKKLAALEQKNAALEYKLAALEK | | |
| **mCC-3(2)6-1** | 3(2)6 | +3(2)+6 | MGSSHHHHHHSSGLVPRGSHMMLAALKEKLAALKEK<br>NAALKYKLAALKEKLGLTPELEILELLLEITELLLK<br>ILELLLKILEEADPNPDPAKLEILELLLEILELLTK<br>ILELLLKILEGAQPLEMLLEEIAQLLEEIAKLLKKI<br>AELLKKIAETTETKKQGDLLEEIAQLLEEIAKLLKK<br>IAELLKKIAEGYGDKRTLLEEIAQLLEEIAKLLKKI<br>AELLKKIAEVAPTQRHRYLLEEIAQLLEEIAKLLKK<br>IAWLLKKIAEG | 92.3 | 0.88 |
| **mCC-3(2)6-2** | 3(2)6 | -3(2)+6 | MGSSHHHHHHSSGLVPRGSHMMLAALKEKLAALKEK<br>NAALKYKLAALKEKLGLTPELLALIAELLATIAELL<br>ALIAELLALIADPNPDPAELLALIAELLALIAETLA<br>LIAELLALIAETTETKKQGDLLEEIAQLLEEIAKLL<br>KKIAELLKKIAEGYGDKRTLLEEIAQLLEEIAKLLK<br>KIAELLKKIAEVAPTQRHRYLLEEIAQLLEEIAKLL<br>KKIAELLKKIAEGTNSDSDLKSLLEEIAQLLEEIAK<br>LLKKIAWLLKKIAEG | 90.3 | 0.86 |
| **mCC-4(1)4** | 4(1)4 | +4(1)-4 | MGSSHHHHHHSSGENLYFQSHMLEEIAEQLEEIAQQ<br>LEEIAQQLKKIAQQTEDGEIAQQLEEIAYQLKKIAE<br>QLKKIAQQLKQGEGEEIAQQLEEIAKQLKKIAQQLK<br>KIAQQTEDGEIALQLIEIALQLIKIALQLIKISTSN<br>GEIAQQLEEIAEQLKKIAEQLKKIAQQLSTSNGEEI<br>AEQLEEIAWQLKKIAEQLKKIAQEEENGEIAQQLEE<br>IAEQLKKIAEQLKKIAQQLKK | 90.9 | 0.86 |
| **mCC-4(2)4** | 4(2)4 | +4(2)-4 | MGSSHHHHHHSSGLVPRGSHMMQLEEIAQQLEEIAK<br>QLKKIAWQLKKIAEGHPHGKGLEEIAQQLEEIAKQL<br>KKIAEQLKKISTAPEEVRNPGLIEILQQLIEILQQL<br>IKILQQLIKILSDGEAKPIFEGLIEILQQLIEILQQ<br>LIKILQQLIKILEDVVPQEGIEELQQAIEELQQAIK<br>KLQEAIKKLQENPGAPEGIEELQQAIEELQQAIKKL<br>QYAIKKLQQ | 93.5 | 0.90 |
| **mCC-4(2)5-1** | 4(2)5 | +5(2)+4 | MGSSHHHHHHSSGLVPRGSHMMAEEIAQTLEEIAQT<br>LKKIAWTLKKIAEKIANPEIEQELEAIAEELKAIAE<br>KLKAIAQKLKAGGSPEEIAQELEAIAQELKAIAEKL<br>KAIAQKIADPIIEQLLEIIEQLLKIIKQLLKIIKQL<br>LKAGGSPEIIEQLLEIIEQLLKIIKQLLKIIYEPEL<br>AQEGLEEIAQQLEEIAKQLKKIAEQLKKIYEETGNK<br>GLEEIAQQLEEIAKQLKKIAYQLKKIAQ | 93.0 | 0.90 |
| **mCC-4(2)5-2** | 4(2)5 | +4(2)+5 | MGSSHHHHHHSSGLVPRGSHMMQLEEIAEQLEEIAK<br>QLKKIAWQLKKLAKGNKYGVQLEEIAEQLEEIAKQL<br>KKIAEQLKSGKLDVQEGIEQLLEIIEQLLKIIKQLL<br>KLILDGAKNGEQLLEIIEQLLKIIKQLLKIIKQLLE<br>QGNLGEIAEELEAIAEELKAIAEKLKAIANVKNPGK<br>GIEQELEAIAEELKAIAEKLKAIAEKLKELGLGEIA<br>QTLEEIAETLKKIAETLKKIAYK | 92.3 | 0.89 |
| **mCC-4(2)6-1** | 4(2)6 | +4(2)-6 | MGSSHHHHHHSSGLVPRGSHMMLLEEIAQLLEEIAQ<br>LLKKIAELLKKIAQGAQPLEMLLEEIAQLLEEIAWL<br>LKKIAELLKKIAQTTETKKQGDLLEEIAQLLEEIAQ<br>LLKKIAELLKKIAQGYGDKRTLLEEIAQLLEEIAEL<br>LKKIAELLKKIAQVAPTQRHRYLLEIIAQLLEIIAQ<br>LLKIIAQLLKIIAQGTNSDSDLKSLLEIIAQLLEII<br>AQLLKIIAQLLKIIASDDEEAEIKGLEEIAQQLEEI<br>AYQLEKIAQQLKKLAKGNKHGKGLEEIAQQLEEIAQ<br>QLKKIAQQLKKIAQG | 94.0 | 0.92 |
| **mCC-4(2)6-2** | 4(2)6 | -6(2)+4 | MGSSHHHHHHSSGLVPRGSHMMLLEEIAQLLEEIAK<br>LLKKIAELLKKIAQGAQPLEMLLEEIAQLLEEIAYL<br>LKKIAELLKKIAQTTETKKQGDLLEIIAQLLEIIAQ<br>LLKIIAQLLKIIASSDEEAQKKGLEEIAQQLEEIAQ<br>QLEKIAQQLKQLGEGEIAQQLEEIAWQLKKIAQQLK<br>KLATGDYGDNFGRTLLEIIAQLLEIIAQLLKIIAQL<br>LKIIAQVAPTQRHRYLLEEIAQLLEEIAKLLKKIAE<br>LLKKIAQGTNSDSDLKSLLEEIAQLLEEIAQLLKKI<br>AELLKKIAQG | 93.5 | 0.92 |
| **mCC-4(2)6-3** | 4(2)6 | -6(2)-4 | MGSSHHHHHHSSGLVPRGSHMMGQLEEIAQQLEEIA<br>QQLKKIAQQLKKLAEGQGDKGLEEIAQQLEEIAYQL<br>EKIAQQLKEGKDGLEIIAQLLEIIAQLLKIIAQLLK<br>IIAQGAQPLEMLLEIIAQLLEIIAQLLKIIAQLLKI<br>IAQTTETKKQGDLLEEIAQLLEEIAKLLKKIAELLK<br>KIAQGYGDKRTLLEEIAQLLEEIAQLLKKIAELLKK<br>IAQVAPTQRHRYLLEEIAQLLEEIAWLLKKIAELLK<br>KIAQGTNSDSDLKSLLEEIAQLLEEIAELLKKIAEL<br>LKKIAQG | 94.6 | 0.92 |
| **mCC-4(2)6-4** | 4(2)6 | +4(2)-6 | MGSSHHHHHHSSGLVPRGSHMQLEEIAQQLEEIAQQ<br>LKKIAQQLKKLAEPQGDKGLEEIAQQLEEIAYQLEK<br>IAQQLKEGKDGLLEILAQLLEILAQLLKILAQLLKI | 79.0 | 0.75 |

| | | | | | |
|---|---|---|---|---|---|
| | | | LAQGAQPLEMLLEILAQLLEILAQLLKILAQLLKIL<br>AQTTETKKQGDSLEELAQSLEELAKSLKKLAESLKK<br>LAQGYGDKRTSLEELAQSLEELAKSLKKLAESLKKL<br>AQVAPTQRHRYSLEELAQSLEELAWSLKKLAESLKK<br>LAQGTNSDSDLKSSLEELAQSLEELAESLKKLAESL<br>KKLAQG | | |
| **mCC-5(2)5** | 5(2)5 | +5(2)+5 | MGSSHHHHHHSSGLVPRGSHMMLEIIALLLEIIALL<br>LEIIALLLKIIALLAEDGEIAQQLEEIAKQLEKIAQ<br>QLKKIAQQLKDNAPGEEIAQQLEEIAKQLEKIAWQL<br>KKIAQQTEDGEIAQLLEEIAKLLEKIAQLLKKIAQL<br>LKQEKVDGLEIIALLLEIIALLLKIIALLLKIIALK<br>AETGEIAQQLEEIAKQLEKIAQQLKKIAQQLKDNAP<br>GEEIAQQLEEIAKQLEKIAYQLKKIAQQTEDGEIAQ<br>LLEEIAKLLEKIAQLLKKIAQLLKQ | 92.6 | 0.85 |
| **mCC-5(1)2(1)5** | 5(1)2(1)5 | +5(1)2(1)+5 | MGSSHHHHHHSSGENLYFQSHMEKIAQTLEKIAKTL<br>EKIAQTLEKIAQQYEEQGQGEIAQQLKEIAKQLKKI<br>AQQLKKIAQLVKDNADPGEIAQQLEEIAKQLEKIAW<br>QLKKIAQQFKEQGQGEIAQQLEEIAKQLEEIAQQLK<br>EIAQLLKKEGNEGLIALQLEIIALQLELIALQLEII<br>ALILNNDEQGLIALTLRIIALTLRLIALTLRIIALK<br>AKEKGLGEIAQQLEEIAKQLEKIAQQLEKIAQLVEN<br>NADPGEIAQQLKEIAKQLKKIAYQLKKIAQQAKELG<br>LGEIAQQLEEIAKQLEKIAQQLEKIAQLLEQPEVDW<br>GEIAQQLKEIAKQLKKIAQQLKKIAQQL | 94.3 | 0.89 |
| **mCC-5(2)6-1** | 5(2)6 | +5(2)+6 | MGSSHHHHHHSSGLVPRGSHMMLEEIAQTLEEIAQT<br>LEKIAYTLKKIAQTAPTGEIAQQLEEIAQQLEKIAE<br>QLKKIAQQLRSGQAQPGEIAQQLEEIAQQLEKIAEQ<br>LKKIAQQYRELGDELGALLLEIIALLLEIIALLLKI<br>IAILSVSLGIDHGIIALLLKIIALLLEIIALLLEII<br>AAKNEEEIKKGLEEIAQLLEEIAQLLKKIAELLKKI<br>AEGHPVGKGLEEIAQLLEEIAQLLKKIAELLKKIAE<br>NPNDKKGLEEIAQLLEEIAQLLKKIAELLKKIAEQE<br>KDEKKKKGLEEIAQLLEEIAQLLKKIAWLLKKIAQ | 92.9 | 0.89 |
| **mCC-5(2)6-2** | 5(2)6 | +5(2)+6 | MGSSHHHHHHSSGENLYFQSHMLEEIAQTLEEIAKT<br>LEKIAQTLKKIAQQAEDGEIAQQLEEIAEQLEKIAW<br>QLKKIAQQLKDNAPGEEIAQQLEEIAKQLEKIAQQL<br>KKIAKQTEDGLIASLLELIASLLELIASLLKLIASL<br>LKQEKVDGKLIASLLKLIASLLELIASLLELILNPE<br>GLEELAQSLEELAKSLKKLAWSLKKLGGKGLEELAQ<br>SLEELAKSLKKLAWSLKQEEVGLEELAQSLEELAKS<br>LKKLAWSLKKLGGEGLEELAQSLEELAKSLKKLAWS<br>LKK | 91.4 | 0.89 |
| **mCC-6(2)6-1** | 6(2)6 | -6(2)+6 | MGSSHHHHHHSSGLVPRGSHMMLLEEIAELLEEIAK<br>LLKKIAWLLKKIAKKPEEGSKEGLEEIAQLLEEIAE<br>LLKKIAELLKKIAEGHPAGKGLEEIAELLEEIAKLL<br>KKIAELLKKIAEKPEEGSKEGLEEIAQLLEEIAELL<br>KKIAELLKKIAKELNVPGLEIIALLLEIIALLLKII<br>ALLLKIIAKGGFNEKIGLEIIALLLEIIALLLKIIA<br>LLLKIIANPEYKKKGLEEIAQLLEEIAKLLKKIAYL<br>LKKIAEGHPYGKGLEEIAELLEEIAELLKKIAELLK<br>KIAEGEEDEKKGLEEIAQLLEEIAKLLKKIAELLKK<br>IAEGHPFGEGLEEIAELLEEIAELLKKIAELLKKIA<br>QG | 92.9 | 0.92 |
| **mCC-6(2)6-2** | 6(2)6 | +6(2)+6 | MGSSHHHHHHSSGENLYFQSGSLLEEIAELLEEIAK<br>LLKKIAWLLKKIAENSEEFDVGKGLEEIAQLLEEIA<br>ELLEKIAELLKKIAEEAETEEEAKGLEEIAELLEEI<br>AKLLKKIAELLKKIAENSEEFDVKKGLEEIAQLLEE<br>IAELLKKIAELLKKIAEKEGNSGLELIASLLELIAS<br>LLKLIASLLKLIAENLGEGSGSGLELIASLLELIAS<br>LLKLIASLLKLIANPGDKEGLEELAQSLEELAKSLK<br>KLAYSLKKLAESVEGEELKEGLEELAESLEELAESL<br>EKLAESLKKLAKGEATQEEVNKGLEELAQSLEELAK<br>SLKKLAESLKKLAEEGEDEGLEELAESLEELAESLE<br>KLAESLKKLAG | 91.2 | 0.90 |
| **mCC-6(2)6-3** | 6(2)6 | +6(2)-6 | MGSSHHHHHHSSGENLYFQSGSSLEELAQSLEELAK<br>SLKKLAYSLKKLAESVEGEELKEGLEELAESLEELA<br>ESLEKLAESLKKLAETGDEELGGKGLEELAQSLEEL<br>AKSLKKLAESLKKLAEEGEDEGLEELAESLEELAES<br>LEKLAESLKKLAEKSGGPTGGLELIASLLELIASLL<br>KLIASLLKLIAENPEEGSGSGLELIASLLELIASLL<br>KLIASLLKLIASKAEKEEVAKGLEEIAELLEEIAKL<br>LKKIAWLLKKIAEDSEEFDKKKGLEEIAQLLEEIAE<br>LLEKIAELLKKIAETAEKEEVAKGLEEIAELLEEIA<br>KLLKKIAELLKKIAEDSEEFDKKKGLEEIAQLLEEI<br>AELLKKIAELLKKIAK | 80.3 | 0.75 |

| | | | | | |
|---|---|---|---|---|---|
| **mCC-6(2)6-4** | 6(2)6 | -6(2)+6 | MGSSHHHHHHSSGENLYFQSGSSLEELAQSLEELAK SLKKLAYSLKKLAKGETEAKEGLEELAESLEELAWS LEKLAESLKKLAEKGGDEGLEELAQSLEELAKSLKK LAESLKKLGEEPNSKEGLEELAESLEELAESLEKLA ESLKKLAEEEKDEVKKQGLELLASLLELLASLLKLL ASLLKLLATTAPENQVPAGLELLASLLELLASLLKL LASLLKLLSNKDESKRKEGLEELAESLEELAESLKK LAESLKKLAETVEKEEHAKGLEELAESLEELAESLE KLAESLKKLAEDLNDKKGLEELAESLEELAESLKKL AESLKKLAELGVFPEGLEELAESLEELAESLKKLAE SLKKLAE | 82.3 | 0.75 |
| **mCC-6(2)6-5** | 6(2)6 | +6(2)+6 | MGSSHHHHHHSSGENLYFQSGSLLELLASLLELLAS LLKLLASLLKLLADEGEDEGLEELAQSLEELAKSLE KLAYSLKKLAEGETEAKEGLEELAESLEELAESLKK LAWSLKKLAEEGGDEGLEELAQSLEELAKSLEKLAE SLKKLAEDPESKEGLEELAESLEELAESLEKLAESL KKLAEEEEDEERRQGLELLASLLELLASLLKLLASL LKLLGGEPSEEELKEGLEELAESLEELAESLKKLAE SLKKLAEEEKDEGLEELAESLEELAESLKKLAESLK KLGENPEDKEGLEELAESLEELAESLKKLAESLKKL AEEEKDEGLEELAESLEELAESLEKLAESLKKLAE | 79.3 | 0.69 |

**Table S5. Proteins passing experimental validation**

| name | expression | CD | SAXS* | xtal |
|---|---|---|---|---|
| **mCC-3(1)3-S** | 1 | 1 | 1 | 1 |
| **mCC-3(1)3-W-1** | 1 | 1 | 1 | 1 |
| **mCC-3(1)3-W-2** | 1 | 1 | 0 | 0 |
| **mCC-3(1)3-Z-1** | 1 | 1 | 1 | 1 |
| **mCC-3(1)3-Z-2** | 1 | 1 | 1 | 0 |
| **mCC-3(1)3-M-1** | 1 | 1 | n/a | 1 |
| **mCC-3(1)3-M-2** | 1 | 1 | 0 | 0 |
| **mCC-3(2)3** | 1 | 1 | 0 | 1 |
| **mCC-3(1)2(1)3** | 1 | 1 | 1 | 0 |
| **mCC-3(2)4-1** | 1 | 1 | n/a | 1 |
| **mCC-3(2)4-2** | 1 | 1 | 0 | 0 |
| **mCC-3(2)4(2)3** | 1 | 1 | 1 | 0 |
| **mCC-3(2)6-1** | 1 | 1 | 1 | 0 |
| **mCC-3(2)6-2** | 0 | 0 | 0 | 0 |
| **mCC-4(1)4** | 1 | 1 | 1 | 0 |
| **mCC-4(2)4** | 1 | 1 | 1 | 0 |
| **mCC-4(2)5-1** | 1 | 1 | 1 | 1 |
| **mCC-4(2)5-2** | 0 | 0 | 0 | 0 |
| **mCC-4(2)6-1** | 0 | 0 | 0 | 0 |
| **mCC-4(2)6-2** | 0 | 0 | 0 | 0 |
| **mCC-4(2)6-3** | 0 | 0 | 0 | 0 |
| **mCC-4(2)6-4** | 1 | 1 | 1 | 0 |
| **mCC-5(2)5** | 1 | 1 | 1 | 1 |
| **mCC-5(1)2(1)5** | 1 | 1 | 1 | 0 |
| **mCC-5(2)6-1** | 0 | 0 | 0 | 0 |
| **mCC-5(2)6-2** | 1 | 1 | 1 | 1 |
| **mCC-6(2)6-1** | 1 | 1 | 1 | 0 |
| **mCC-6(2)6-2** | 1 | 1 | 1 | 0 |
| **mCC-6(2)6-3** | 0 | 0 | 0 | 0 |
| **mCC-6(2)6-4** | 0 | 0 | 0 | 0 |
| **mCC-6(2)6-5** | 0 | 0 | 0 | 0 |
| total | 23/31 | 23/31 | 17/31 | 10/31 |

*passed when $\chi^2 < 3$

**Table S6. Merging and refinement statistics for crystal structures in this study.**

| | sc-apCC-5 | mCC-3(1)3-S | mCC-3(1)3-W-1 | mCC-3(1)3-Z-1 |
|---|---|---|---|---|
| PDB id | 31WI | 31WJ | 31WK | 31WL |
| Wavelength | 0.95 Å | 0.62 Å | 0.62 Å | 0.62 Å |
| Resolution range | 50.94 - 2.05 (2.14 - 2.05) | 19.94-2.4 (2.52-2.4) | 83.82-2.2 (2.24-2.20) | 69.22-2.17 (2.48-2.17) |
| Space group | $P2_12_12_1$ | $P2_12_12_1$ | $P6_222$ | $P2_12_12_1$ |
| Unit cell | 62.29 Å<br>63.76 Å<br>88.52 Å<br>90º 90º 90º | 44.579 Å<br>63.083 Å<br>71.473 Å<br>90º 90º 90º | 96.791 Å<br>96.791 Å<br>103.672 Å<br>90º 90º 120º | 47.311 Å<br>87.210 Å<br>113.791 Å<br>90º 90º 90º |
| Total reflections | 372457 (48682) | 44582 (2409) | 547326 (16263) | 94523 (4169) |
| Unique reflections | 28042 (3470) | 5694 (285) | 15045 (752) | 12508 (625) |
| Multiplicity | 13.3 (14.0) | 7.8 (8.5) | 36.4 (21.6) | 7.6 (6.7) |
| Completeness (%) | 99.97 (99.89) | 91.1 (92.3) | 99.0 (92.0) | 88.3 (67.4) |
| Mean I/sigma(I) | 12.18 (0.66) | 9.4 (2.4) | 16.9 (2.8) | 4.6 (1.8) |
| Wilson B-factor | 50.32 | 16.53 | 19.63 | 28.05 |
| R-pim | 0.02244 (1.203) | 0.083 (0.501) | 0.049 (0.44) | 0.135 (0.591) |
| CC1/2 | 1 (0.38) | 0.99 (0.85) | 1 (0.58) | 1 (0.69) |
| Reflections used in refinement | 22757 (2798) | 5656 (1616) | 15009 (2849) | 12488 (182) |
| Reflections used for R-free | 1144 (101) | 295 (76) | 722 (144) | 638 (7) |
| R-work | 0.2161 (0.3372) | 0.2539 (0.2842) | 0.2278 (0.2583) | 0.2339 (0.3587) |
| R-free | 0.2525 (0.3759) | 0.2951 (0.3158) | 0.2498 (0.2730) | 0.2732 (0.3530) |
| Number of non-hydrogen atoms | 2371 | 1275 | 1373 | 2638 |
| macromolecules | 2371 | 1208 | 1171 | 2551 |
| ligands | 0 | 0 | 0 | 0 |
| solvent | 0 | 67 | 202 | 87 |
| Protein residues | 324 | 164 | 160 | 353 |
| RMS(bonds) | 0.001 | 0.003 | 0.002 | 0.002 |
| RMS(angles) | 0.31 | 0.6 | 0.55 | 0.44 |
| Ramachandran favored (%) | 98.44 | 97.53 | 99.36 | 99.14 |
| Ramachandran allowed (%) | 1.56 | 1.85 | 0.64 | 0.86 |
| Ramachandran outliers (%) | 0 | 0.62 | 0 | 0 |
| Rotamer outliers (%) | 0.93 | 0 | 0 | 0.43 |
| Clashscore | 0.63 | 4.08 | 3.71 | 4.23 |
| Average B-factor | 66.9 | 22.65 | 27.27 | 37.92 |
| macromolecules | 66.9 | 22.71 | 26.2 | 38.03 |
| ligands | / | / | / | / |
| solvent | / | 21.55 | 33.45 | 34.5 |

**Table S6 cont. Merging and refinement statistics for crystal structures in this study.**

| | mCC-3(1)3-M-1 | mCC-3(2)3 | mCC-3(2)4-1 | mCC-4(2)5-1 |
|---|---|---|---|---|
| PDB id | 31WN | 31WO | 31WP | 31WQ |
| Wavelength | 1.0 Å | 0.98 Å | 0.62 Å | 0.954 Å |
| Resolution range | 77.16 – 2.45 (2.48-2.45) | 9.85 - 2.4 (2.64 - 2.4) | 27.67 - 1.65 (1.75 - 1.65) | 52.17 - 2.2 (2.42 - 2.2) |
| Space group | C2 | R $3_2$:H | C2 | $P2_1$ |
| Unit cell | 118.468 Å 43.472 Å 90.81 Å 90° 121.87° 90° | 111.55 Å 111.55 Å 125.56 Å 90° 90° 120° | 99.944 Å 26.57 Å 56.835 Å 90° 103.13° 90° | 45.825 Å 44.188 Å 54.861 Å 90° 108.01° 90° |
| Total reflections | 70866 ((3319) | 252889 (45180) | 215827 (26482) | 73123 (18394) |
| Unique reflections | 11925 (596) | 13570 (3346) | 17913 (2917) | 10764 (2657) |
| Multiplicity | 5.9 (5.6) | 18.6 (13.5) | 12.0 (9.1) | 6.8 (6.9) |
| Completeness (%) | 85.1 (44.0) | 99.92 (99.83) | 95.63 (87.22) | 99.59 (98.98) |
| Mean I/sigma(I) | 6.6 (1.6) | 8.11 (1.06) | 11.96 (2.25) | 9.84 (1.59) |
| Wilson B-factor | 45.83 | 46.29 | 20.78 | 48.86 |
| R-pim | 0.088 (0.802) | 0.2634 (2.608) | 0.02767 (0.3355) | 0.02985 (0.3753) |
| CC1/2 | 1 (0.36) | 0.998 (0.47) | 0.999 (0.855) | 0.999 (0.761) |
| Reflections used in refinement | 11915 (536) | 11933 (2939) | 17131 (2545) | 10720 (2629) |
| Reflections used for R-free | 581 (25) | 628 (188) | 866 (124) | 514 (111) |
| R-work | 0.2426 (0.3329) | 0.2387 (0.2989) | 0.1874 (0.2299) | 0.2275 (0.3077) |
| R-free | 0.2897 (0.3624) | 0.2575 (0.3264) | 0.2268 (0.2670) | 0.2650 (0.3488) |
| Number of non-hydrogen atoms | 2359 | 1846 | 1408 | 1620 |
| macromolecules | 2341 | 1812 | 1315 | 1618 |
| ligands | 0 | 4 | 23 | 0 |
| solvent | 18 | 30 | 70 | 2 |
| Protein residues | 347 | 260 | 169 | 224 |
| RMS(bonds) | 0.002 | 0.002 | 0.004 | 0.005 |
| RMS(angles) | 0.4 | 0.36 | 0.33 | 0.65 |
| Ramachandran favored (%) | 98.23 | 100.00 | 98.2 | 99.09 |
| Ramachandran allowed (%) | 1.77 | 0.00 | 1.8 | 0.91 |
| Ramachandran outliers (%) | 0 | 0.00 | 0 | 0 |
| Rotamer outliers (%) | 0.55 | 0.70 | 0 | 1.41 |
| Clashscore | 3.43 | 3.06 | 2.24 | 5.84 |
| Average B-factor | 51.67 | 59.74 | 29.38 | 62.48 |
| macromolecules | 51.75 | 59.81 | 28.52 | 62.49 |
| ligands | / | 56.07 | 48.88 | / |
| solvent | 40.89 | 56.38 | 39.12 | 59.12 |

**Table S6 cont. Merging and refinement statistics for crystal structures in this study.**

| | **mCC-5(2)5** | **mCC-5(2)6-2** | **mCC-3(1)3-Z-2-MCL1-1:MCL-1** |
|---|---|---|---|
| PDB id | 31WS | 31WT | 31WU |
| Wavelength | 0.62 Å | 0.954 Å | 0.69 Å |
| Resolution range | 39.69 - 2.16<br>(2.3 - 2.16) | 50.21 - 3.491<br>(3.84 - 3.49) | 47.22 - 3.0<br>(3.19 - 3.0) |
| Space group | $P2_12_12_1$ | $P2_12_12_1$ | $P2_12_12_1$ |
| Unit cell | 42.447 Å<br>56.244 Å<br>111.888 Å<br>90º 90º 90º | 65.117 Å<br>69.826 Å<br>72.255 Å<br>90º 90º 90º | 56.19 Å<br>78.7 Å<br>174.18 Å<br>90º 90º 90º |
| Total reflections | 188666 (23815) | 59400 (15464) | 209062 (35769) |
| Unique reflections | 14988 (2432) | 4486 (1098) | 16121 (2628) |
| Multiplicity | 12.6 (9.8) | 13.2 (14.1) | 13.0 (13.6) |
| Completeness (%) | 99.73 (98.85) | 99.67 (99.27) | 99.88 (99.85) |
| Mean I/sigma(I) | 10.49 (1.96) | 10.88 (0.49) | 7.19 (1.11) |
| Wilson B-factor | 41.84 | 164.88 | 87.04 |
| R-pim | 0.03327 (0.3208) | 0.02711 (0.8836) | 0.07217 (0.7154) |
| CC1/2 | 0.976 (0.84) | 0.996 (0.434) | 0.998 (0.749) |
| Reflections used in refinement | 14946 (2404) | 4472 (1091) | 16110 (2627) |
| Reflections used for R-free | 773 (125) | 506 (118) | 806 (159) |
| R-work | 0.2161 (0.2878) | 0.2624 (0.3404) | 0.2782 (0.3714) |
| R-free | 0.2485 (0.3555) | 0.2994 (0.3640) | 0.2991 (0.3625) |
| Number of non-hydrogen atoms | 1894 | 2106 | 4711 |
| macromolecules | 1868 | 2106 | 4711 |
| ligands | 8 | 0 | 0 |
| solvent | 18 | 0 | 0 |
| Protein residues | 255 | 271 | 647 |
| RMS(bonds) | 0.003 | 0.002 | 0.002 |
| RMS(angles) | 0.4 | 0.39 | 0.46 |
| Ramachandran favored (%) | 99.6 | 97.03 | 95.62 |
| Ramachandran allowed (%) | 0.4 | 2.6 | 3.6 |
| Ramachandran outliers (%) | 0 | 0.37 | 0.78 |
| Rotamer outliers (%) | 0.57 | 0.44 | 0 |
| Clashscore | 2.59 | 5.01 | 5.19 |
| Average B-factor | 52.91 | 162.52 | 82.66 |
| macromolecules | 52.86 | 162.52 | 82.66 |
| ligands | 66.75 | / | / |
| solvent | 52.57 | / | / |

**Table S7. Protein sequences for haem and zinc double-binder passing in silico filters and experimental validation.**

| name | architecture | topology | sequence |
|---|---|---|---|
| mCC-4(2)4-ZN-HEM | 4(2)4 | +4(2)-4 | QLEEHAQQAEEIAKQLKKIAWQLKKIAEGSPSGKGLEEIAQQLE EIAKQLKKLAETGKKLSTAPEEVRNPGLIEHLQQAIEILQQLIK ILQQLIKIHSGGEAKPIHEGLHEILQQLIEILQQLIKLLQTGIK LLEDVVPQEGIEELQQAIEELQQAIKKLQEAIKKGQENSGAPSG IEELQQAIEELQQAIKKLQYAIKKLQQ |

**Table S8. Protein sequences for MCL-1 double-binder passing in silico filters and experimental validation.**

| name | architecture | topology | # binding sites | sequence | pAE | ipTM | $IC_{50}$ (nM) |
|---|---|---|---|---|---|---|---|
| mCC-3(1)3-W-2-MCL1-1 | 3(1)3 | -3(1)-3 | 1 | SHMELAALEEELAALEWENAALE EELAALEEAAEEGEDVSEGLAAL KEKLAATKEKLAALKEKLAALKE KQPDSVGLKLLELKLKLLENKTK LLELKLKLLELKYYGDEEEKEKG LAALEEELAALEEETAALEEELA ALEKEQPNSPGLLALEQELLRTI GDLVNLKQELLNLKEG | 3.67 | 0.91 | 380 ± 30 |
| mCC-3(1)3-W-2-MCL1-2 | 3(1)3 | -3(1)-3 | 2 | SHMELLALEEELLRLIGDNVNLE QELLNLEEAAEEGEDVSEGLAAL KEKLAATKEKLAALKEKLAALKE KQPDSVGLKLLELKLKLLENKTK LLELKLKLLELKYYGDEEEKEKG LLALEQELLRLIGDTVNLEQELL NLEKEQPNSPGLAALKEELAATK YELAALKEELAALKEG | 5.12 | 0.87 | 17000 ± 1000 |
| mCC-3(1)3-Z-1-MCL1-1 | 3(1)3 | +3(1)-3 | 1 | SHMKLLALEQKLLRLIGDNVNLK QKLLNLKKKLGASAGLAALEEEL AATEEELAALEEELAALEKAGLS EKEKELGLKLLELKLKLLELKTK NLELKLKLLELKTAPESVRVPGL AALEEELAATEEELAALEEELAA LEKEVPEKYKAGLAALKEELAAL KYETAALKEELAALKEG | 4.12 | 0.91 | 200 ± 16 |
| mCC-3(1)3-Z-1-MCL1-2 | 3(1)3 | +3(1)-3 | 2 | SHMMKLLALEQKLLRLIGDNVNL KQKLLNLKKKLGASAGLAALEWE LAATEEELAALEEELAALEKAGL SEKEKELGLKLLELKLKLLELKT KNLELKLKLLELKTAPESVRVPG LAALEEELAATEEELAALEEELA ALEKEVPEKYKAGLLALEQELLR LIGDTVNLKQELLNLKEG | 5.15 | 0.86 | 85 ± 4 |
| mCC-3(1)3-Z-2-MCL1-1 | 3(1)3 | +3(1)-3 | 2 | SHMMKLLALEQKLLRLIGDNVNL KQKLLNLKKKLGASAGLAALEWE LAATEEELAALEEELAALEKAGL SEKEKELGLKLLELKLKLLETKT KLLELKLKLLELKTAPESVRVPG LLALEQELLRLIGDTVNLEQELL NLEKEVPEKYKAGLAALKEELAA NKYELAALKEELAALKEG | 5.20 | 0.87 | 98 ± 3 |
| mCC-3(1)3-Z-2-MCL1-2 | 3(1)3 | +3(1)-3 | 1 | SHMKLAALKEKLAALKWKNAALK EKLAALKKKLGASAGLAALEWEL AATEEELAALEEELAALEKAGLS EKEKELGLKLLELKLKLLETKTK LLELKLKLLELKTAPESVRVPGL LALEQELLRLIGDTVNLEQELLN LEKEVPEKYKAGLAALKEELAAN KYELAALKEELAALKEG | 7.04 | 0.74 | 12900 ± 700 |

**Table S9. Protein sequences for ligand-protein dual binder and experimental validation.**

| name | architecture | topology | sequence | $K_{D(Nile Red)}$ (μM) | $IC_{50(MCL1)}$ (nM) |
|---|---|---|---|---|---|
| mCC-3(2)4-MCL1-1 | 3(2)4 | +4(2)-3 | MGSSHHHHHHSSGENLYFQSGSGTMQLEEI AQQLEEIAEQLKKIAEQLKKIAKGHPNGKG LEEIAQQLEEIAEQLKKIAWQLKKIATAPE EERVPGLLEIAEQLLEIAEQLTKIAEQLLK IAEQGNDKGLLEIAKQLTEIATQLLNIAGQ | not binding | 33 ± 2 |

| | | | | | |
|---|---|---|---|---|---|
| | | | LLEIASEGGVDEEGLEALISKLLSLIGDNV<br>YLEQKLKKLE | | |
| mCC-3(2)4-MCL1-2 | 3(2)4 | +4(2)-3 | MGSSHHHHHHSSGENLYFQSGSGTMQLEEI<br>AQQLEEIAEQLKKIAEQLKKIAKGHPNGKG<br>LEEIAQQLEEIAEQLKKIAWQLKKIATAPE<br>EERVPGLLEIAEQLLEIAEQLTKIAEQLLK<br>IAEQGNDKGLLEIAKQLTEIAKQLLKIAGQ<br>LLNIAAEGEKDIVGLLKLIGDLLYLEQYNA<br>RLEYKLAALE | not binding | 42 ± 2 |
| mCC-3(2)4-SN38-1 | 3(2)4 | +4(2)-3 | MGSSHHHHHHSSGENLYFQSGSGTMQLEEI<br>AQQLEEGAEQTKKNAEQLKKIAKGHPNGKG<br>LEEIAQQAEEIAEQSKKIAWQLKKIATAPE<br>EERVPGLLEIAEQLLEIAEQATKGAEQLLK<br>IAEQGNDKGLLEIAKQATEGAKQALKIAKQ<br>LLKIAREGGVDEEGLAALEKKLAALEQKNA<br>ALEYKLAALE | 0.48 ± 0.04 | not binding |
| mCC-3(2)4-MCL1-1-SN38-1 | 3(2)4 | +4(2)-3 | MGSSHHHHHHSSGENLYFQSGSGTMQLEEI<br>AQQLEEGAEQTKKNAEQLKKIAKGHPNGKG<br>LEEIAQQAEEIAEQSKKIAWQLKKIATAPE<br>EERVPGLLEIAEQLLEIAEQATKGAEQLLK<br>IAEQGNDKGLLEIAKQATEGATQALNIAGQ<br>LLEIASEGGVDEEGLEALISKLLSLIGDNV<br>YLEQKLKKLE | 1.29 ± 0.06 | 43 ± 2 |
| mCC-3(2)4-MCL1-2-SN38-1 | 3(2)4 | +4(2)-3 | MGSSHHHHHHSSGENLYFQSGSGTMQLEEI<br>AQQLEEGAEQTKKNAEQLKKIAKGHPNGKG<br>LEEIAQQAEEIAEQAKKIAWQLKKIATAPE<br>EERVPGLLEIAEQLLEIAEQATKGAEQLLK<br>IAEQGNDKGLLEIAKQATEGAKQALKIAGQ<br>LLNIAAEGEKDIVGLLKLIGDLLYLEQYNA<br>RLEYKLAALE | 1.77 ± 0.05 | 44 ± 4 |

**Table S10. Protein sequences to test dual binder in cells.**

| name | architecture | topology | sequence |
|---|---|---|---|
| mCC-3(2)4-MCL1-2 | 3(2)4 | +4(2)-3 | MQLEEIAQQLEEIAEQLKKIAEQLKKIAKGHPNGKGLEEIAQQLEEIAEQLKKIA<br>WQLKKIATAPEEERVPGLLEIAEQLLEIAEQLTKIAEQLLKIAEQGNDKGLLEIA<br>KQLTEIAKQLLKIAGQLLNIAAEGEKDIVGLLKLIGDLLYLEQYNARLEYKLAAL<br>E |
| mCC-3(2)4-SN38-1 | 3(2)4 | +4(2)-3 | MQLEEIAQQLEEGAEQTKKNAEQLKKIAKGHPNGKGLEEIAQQAEEIAEQSKKIA<br>WQLKKIATAPEEERVPGLLEIAEQLLEIAEQATKGAEQLLKIAEQGNDKGLLEIA<br>KQATEGAKQALKIAKQLLKIAREGGVDEEGLAALEKKLAALEQKNAALEYKLAAL<br>E |
| mCC-3(2)4-MCL1-2-SN38-1 | 3(2)4 | +4(2)-3 | MQLEEIAQQLEEGAEQTKKNAEQLKKIAKGHPNGKGLEEIAQQAEEIAEQAKKIA<br>WQLKKIATAPEEERVPGLLEIAEQLLEIAEQATKGAEQLLKIAEQGNDKGLLEIA<br>KQATEGAKQALKIAGQLLNIAAEGEKDIVGLLKLIGDLLYLEQYNARLEYKLAAL<br>E |
| eGFP-MCL1(424-578) | - | - | MVSKGEELFTGVVPILVELDGDVNGHKFSVSGEGEGDATYGKLTLKFICTTGKLP<br>VPWPTLVTTLTYGVQCFSRYPDHMKQHDFFKSAMPEGYVQERTIFFKDDGNYKTR<br>AEVKFEGDTLVNRIELKGIDFKEDGNILGHKLEYNYNSHNVYIMADKQKNGIKVN<br>FKIRHNIEDGSVQLADHYQQNTPIGDGPVLLPDNHYLSTQSALSKDPNEKRDHMV<br>LLEFVTAAGITLGMDELYKGSGSGSGSMELYRQSLEIISRYLREQATGAKDTKPM<br>GRSGATSRKALETLRRVGDGVQRNHETAFQGMLRKLDIKNEDDVKSLSRVMIHVF<br>SDGVTNWGRIVTLISFGAFVAKHLKTINQESCIEPLAESITDVLVRTKRDWLVKQ<br>RGWDGFVEFFHVEDLEGGIRNVLLAFAGVAGVGAGLAYLIR |

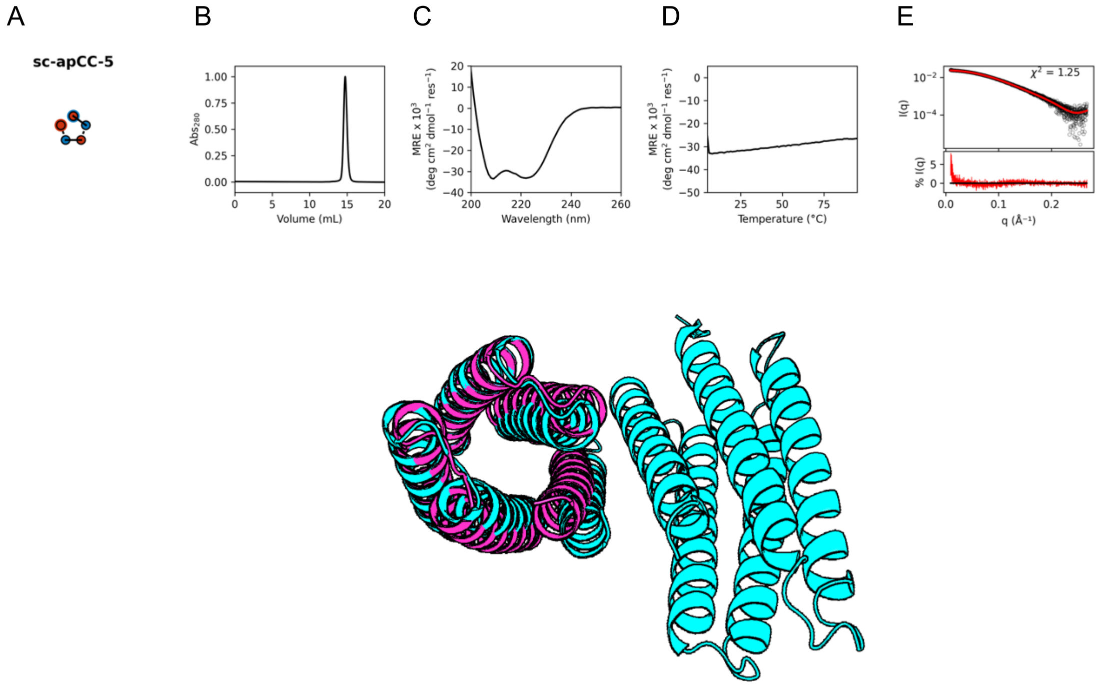


**Figure S1. Experimental characterization of sc-apCC-5.** Topology diagram showing helical interconnectivity. **(B)** Normalized analytical size exclusion chromatograms. Conditions: Superdex™ 75 Increase 10/300 GL, 50 mM phosphate buffer pH 7.4, 150mM NaCl **(C)** CD spectra recordered at 5 °C. Filled grey regions denote hight tension voltage ≥ 600 V. MRE, mean residue ellipticity (deg $cm^2$ $dmol^{-1}$ $res^{-1}$). Conditions: 5-10 µM protein, 50 mM sodium phosphate and 150 mM NaCl (pH 7.4). **(D)** Temperature dependent CD signal, monitored at 222 nm. Scans were collected heating to (solid lines) and cooling from (dashed lines) 95 °C. MRE, mean residue ellipticity (deg $cm^2$ $dmol^{-1}$ $res^{-1}$). **Conditions:** 10 µM protein, 50 mM sodium phosphate and 150 mM NaCl (pH 7.4). (E) Small-angle X-ray scattering profiles fitted to the AF2 models returning using MultiFoXS[13]. Residuals are shown as percentage of maximum scattering intensity. (F) Crystal structure (cyan) overlaid with the AF2 model (magenta). Resolution 2.1 Å; R = 0.21; $R_{free}$ = 0.24, PDB ID is 31WI.

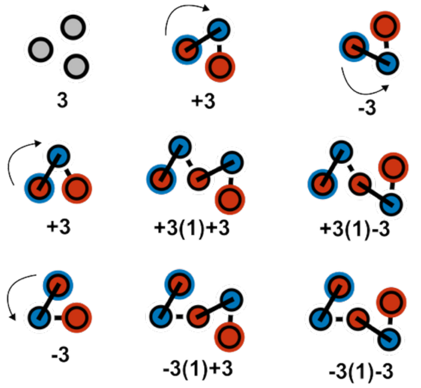


**Figure S2. Extended topology notations illustrated using sc-apCC-3 based systems.** Two different topologies of sc-apCC-3 (3): clockwise (+3) and anticlockwise (-3) gives four combinations, i.e. there are four topologies of 3(1)3 architecture. In the topology diagrams grey discs indicate standalone helices, blue and red central discs mark the N- and C-termini of each helix, and outer blue and red rings show the positions of the N- and C-terminal helices in single chains, respectively.

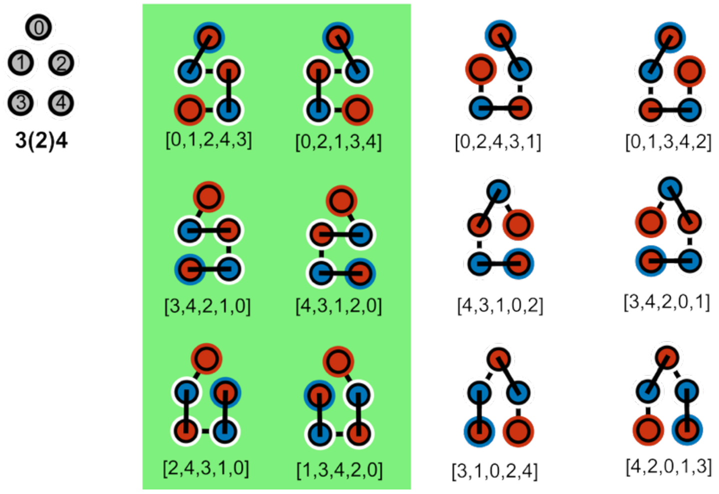


**Figure S3. Topological constraints considered during design.** Of the 120 permutations of five helices [0,1,2,3,4] of 3(2)4 architecture, only 42 produce continuous looping, in which neighbouring helices are connected (e.g., not 2–3 or 0–4; only selected examples are shown). Among these, only six permutations ensure that all the helices of the N-terminal module appear in the design topology first, followed by the remaining helices of the second module (green background).

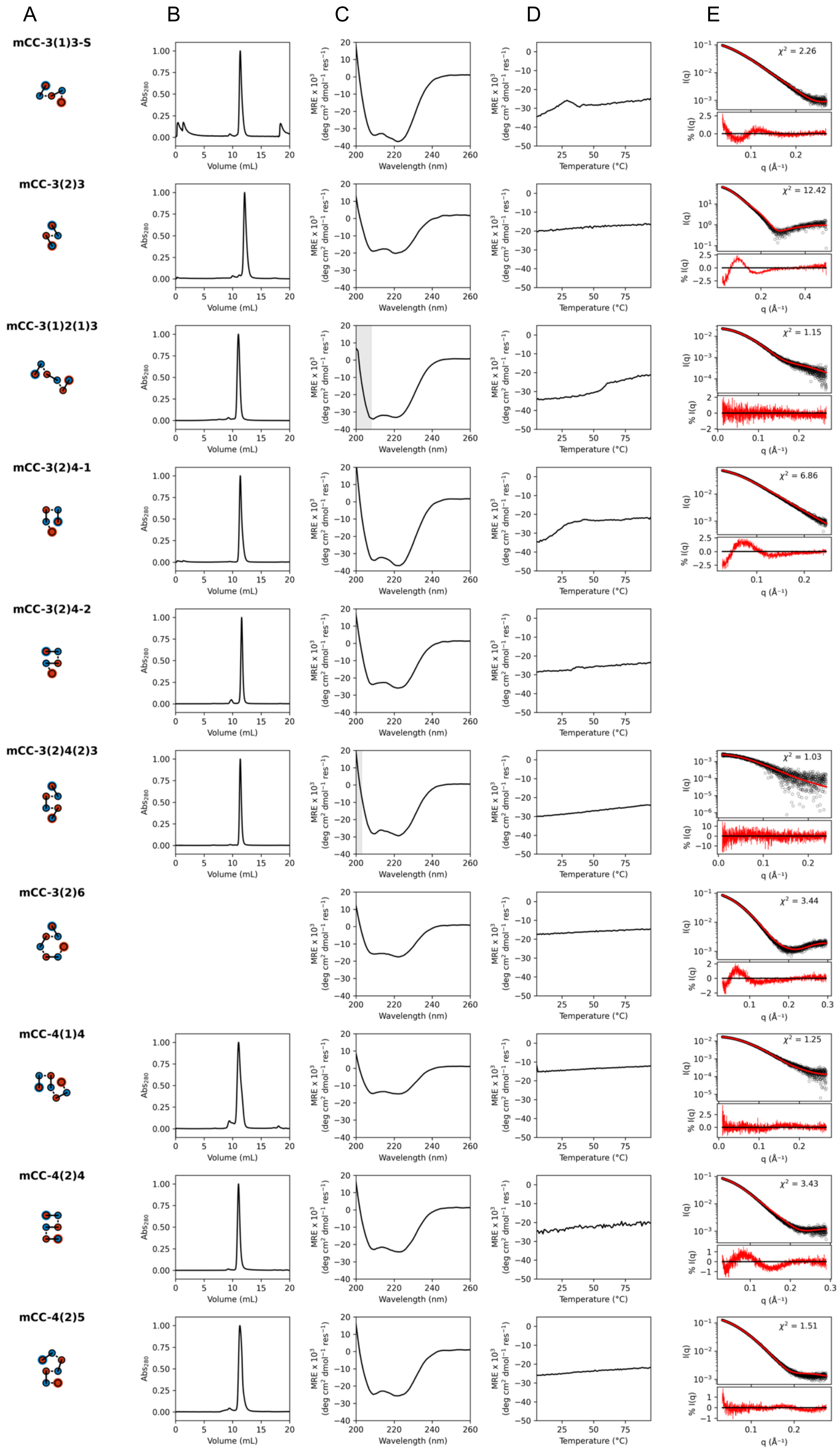
A
B
C
D
E
mCC-3(1)3-S
mCC-3(2)3
mCC-3(1)2(1)3
mCC-3(2)4-1
mCC-3(2)4-2
mCC-3(2)4(2)3
mCC-3(2)6
mCC-4(1)4
mCC-4(2)4
mCC-4(2)5
Abs280
Volume (mL)
MRE x 10³ (deg cm² dmol⁻¹ res⁻¹)
Wavelength (nm)
Temperature (°C)
I(q)
% I(q)
q (Å⁻¹)
χ² = 2.26
χ² = 12.42
χ² = 1.15
χ² = 6.86
χ² = 1.03
χ² = 3.44
χ² = 1.25
χ² = 3.43
χ² = 1.51

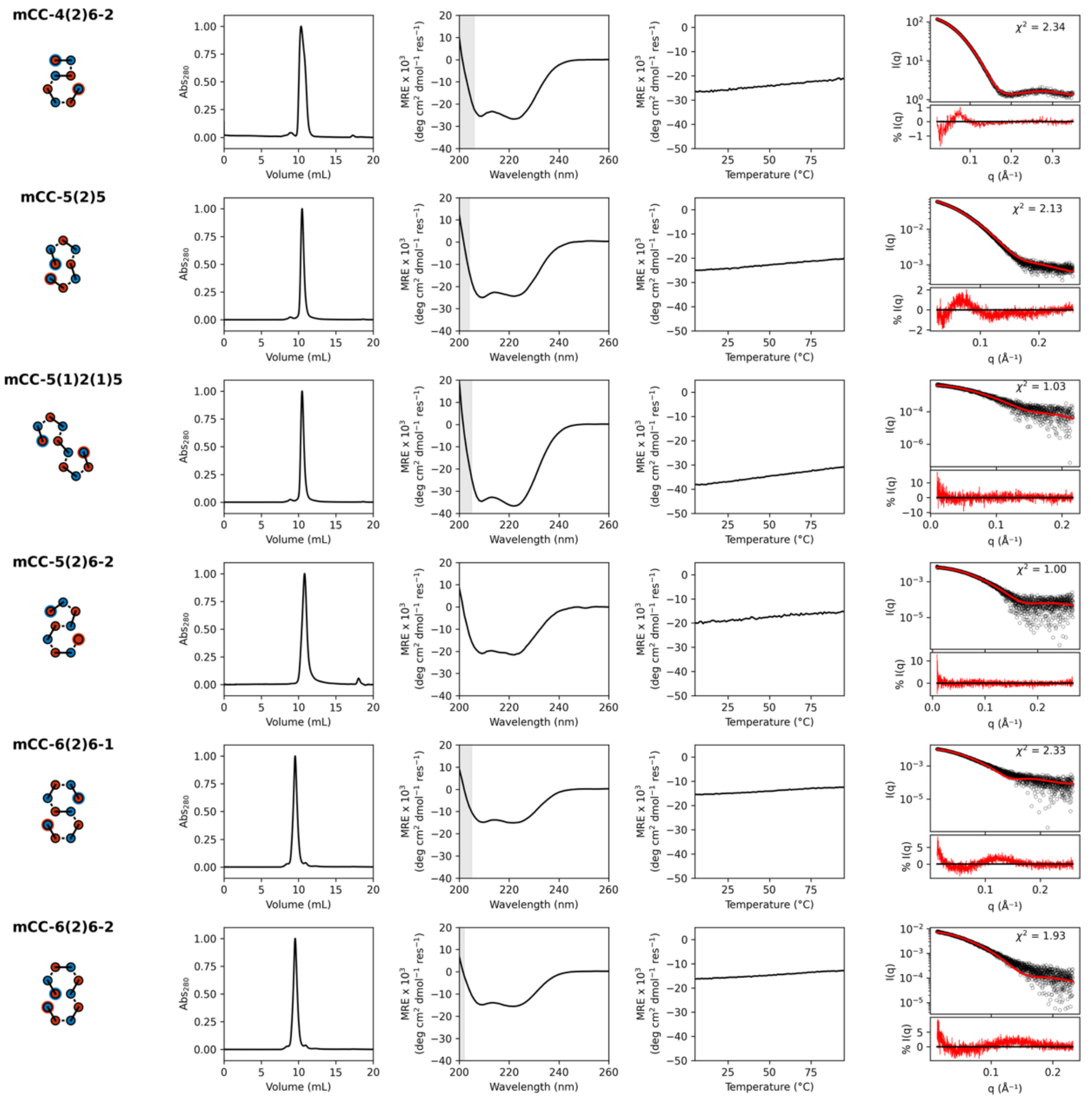


**Figure S4. Biophysical characterization of mCCs representing diverse architectures**. Topology diagrams showing helical interconnectivity. **(B)** Normalized analytical size exclusion chromatograms. Conditions: Superdex™ 75 Increase 10/300 GL, 50 mM phosphate buffer pH 7.4, 150mM NaCl **(C)** CD spectra recorded at 5 °C. Filled grey regions denote hight tension voltage ≥ 600 V. MRE, mean residue ellipticity (deg cm$^2$ dmol$^{-1}$ res$^{-1}$). Conditions: 5-10 µM protein, 50 mM sodium phosphate and 150 mM NaCl (pH 7.4). **(D)** Temperature dependent CD signal, monitored at 222 nm. Scans were collected heating to (solid lines) and cooling from (dashed lines) 95 °C. MRE, mean residue ellipticity (deg cm$^2$ dmol$^{-1}$ res$^{-1}$). **Conditions:** 10 µM protein, 50 mM sodium phosphate and 150 mM NaCl (pH 7.4). (E) Small-angle X-ray scattering profiles fitted to the AF2 models returning using MultiFoXS[13]. Residuals are shown as percentage of maximum scattering intensity.

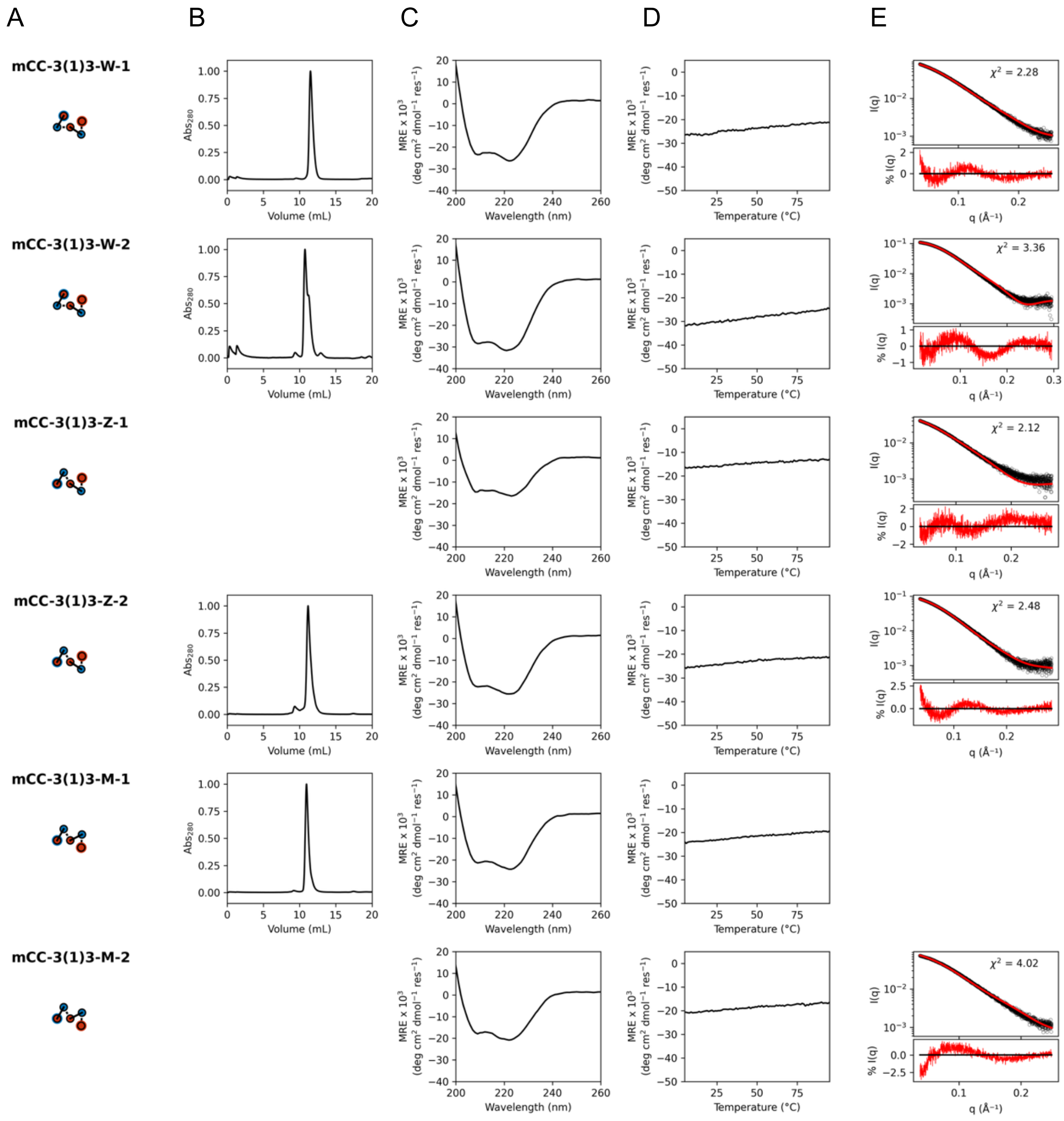


**Figure S5. Biophysical characterization of mCCs representing 3(1)3 architectures**. Topology diagrams showing helical interconnectivity. **(B)** Normalized analytical size exclusion chromatograms. Conditions: Superdex™ 75 Increase 10/300 GL, 50 mM phosphate buffer pH 7.4, 150mM NaCl **(C)** CD spectra recorded at 5 °C. Filled grey regions denote hight tension voltage ≥ 600 V. MRE, mean residue ellipticity (deg cm2 dmol-1 res-1). Conditions: 5-10 μM protein, 50 mM sodium phosphate and 150 mM NaCl (pH 7.4). **(D)** Temperature dependent CD signal, monitored at 222 nm. Scans were collected heating to (solid lines) and cooling from (dashed lines) 95 °C. MRE, mean residue ellipticity (deg $cm^2$ $dmol^{-1}$ $res^{-1}$). **Conditions:** 10 μM protein, 50 mM sodium phosphate and 150 mM NaCl (pH 7.4). (E) Small-angle X-ray scattering profiles fitted to the AF2 models returning using MultiFoXS[13]. Residuals are shown as percentage of maximum scattering intensity.

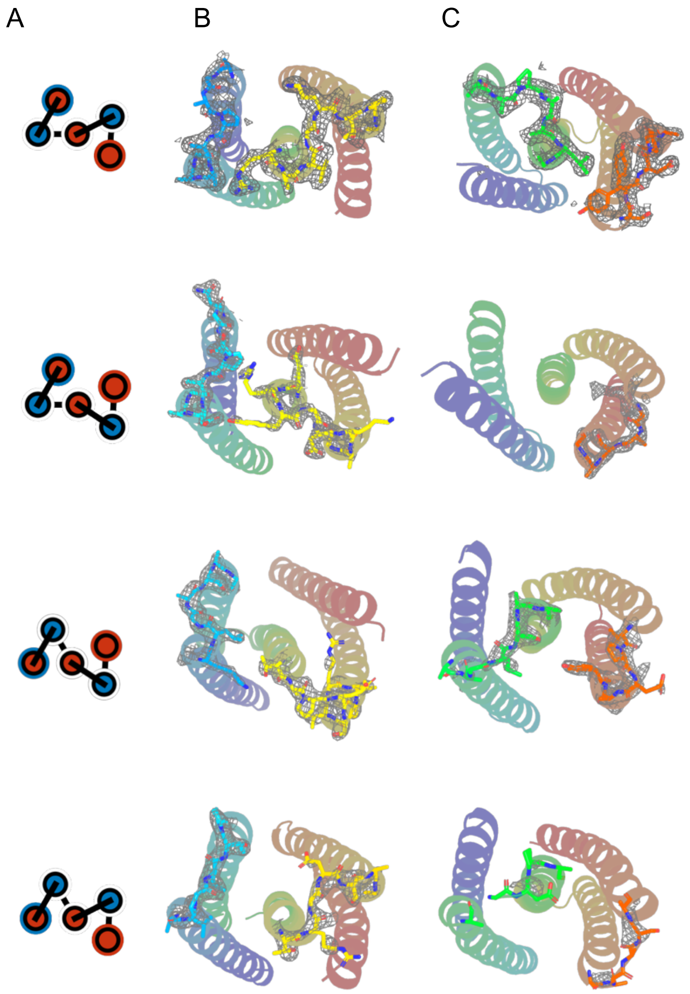


**Figure S6. Electron density surrounding the loop regions of four X-ray crystal structures of 3(1)3.** (**A**) Topology diagrams for mCC-3(1)3-S, mCC-3(1)3-W-1, mCC-3(1)3-Z-1 and, CC-3(1)3-M-1, top to bottom. (**B**) Electron density around first and third loop. (**C**) Electron density around second and fourth loop. Each structure is coloured by chain using a chainbow palette; refined loop residues are shown as sticks and the electron density is contoured at 1.0σ. Data collection and refinement statistics are provided in Table S5; PDB IDs: 31WJ, 31WK, 31WL and 31WN.

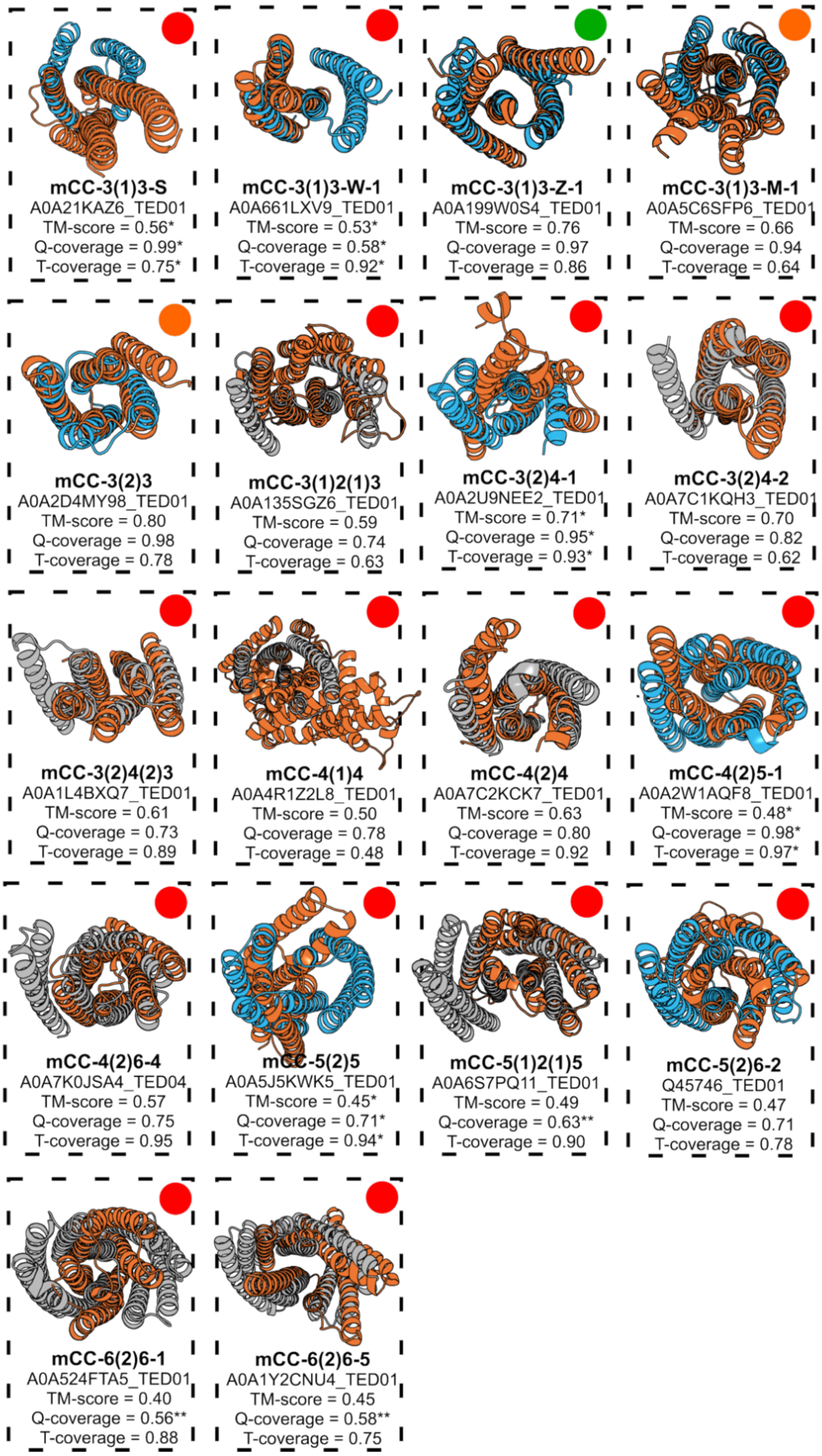


**Figure S7. Top structural matches for each designed mCC based on TM-align and FoldSeek.** For each of the 18 designed mCCs, the top-scoring TED domain identified through the FoldSeek + TM-align workflow (orange) is shown superimposed on the mCC X-ray crystal structure (blue) or AF2 model (grey). Circle markers indicate the confidence class based on similarity: green = clear structural match; orange = borderline or ambiguous similarity; red = no clear match. TM-scores correspond to the query-normalized values as calculated by TM-align except when noted otherwise, query (Q) and target (T) coverage values are also reported. *TM-align could not find a succesful superimposition,TM-score and coverages as reported by FoldSeek, **highest query coverage was below 0.7.

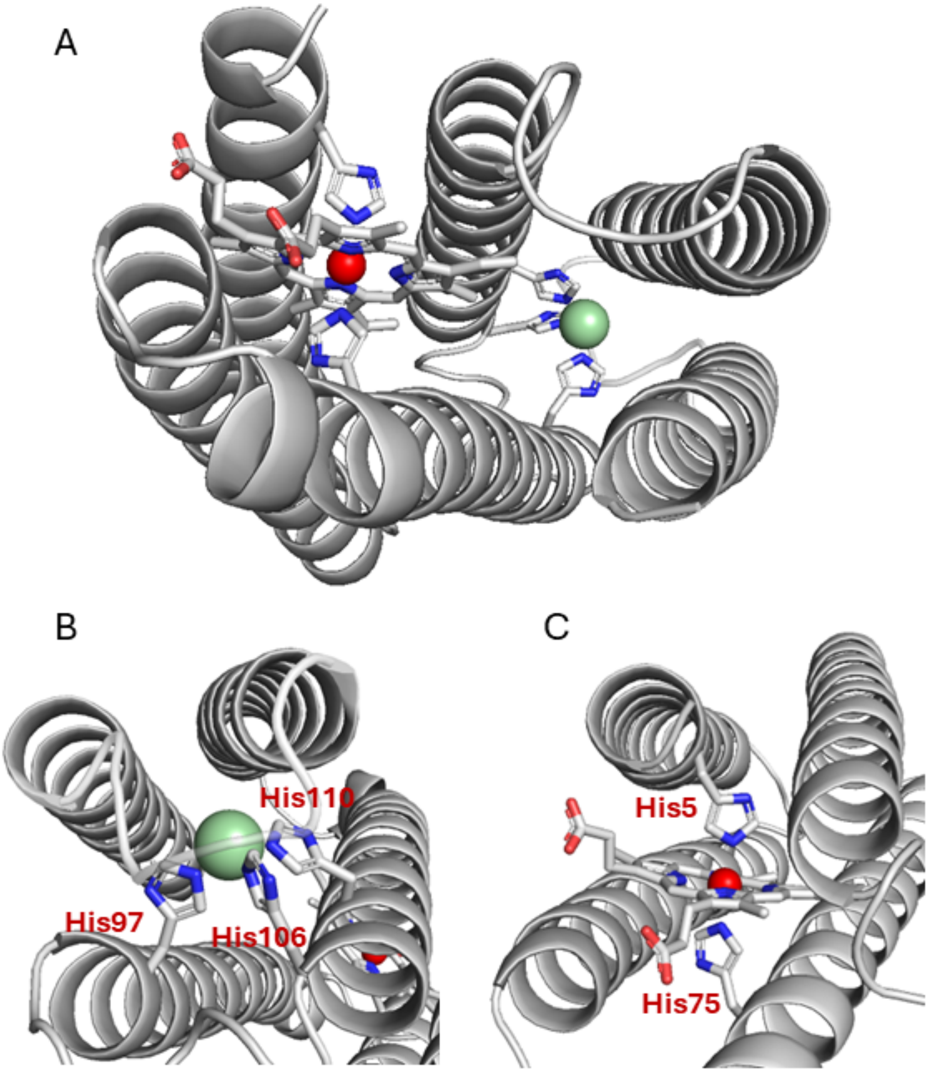


**Figure S8. AlphaFold3-predicted structure of the double binder mCC-4(2)4-ZN-HEM variant in the presence of haem and $Zn^{2+}$.** (A) Overall predicted structure showing both ligands bound to the protein scaffold and the designed His residues involved in haem and $Zn^{2+}$ coordination. The model shows high overall confidence, with a global pLDDT of 88.5, pTM of 0.88 and iPTM of 0.85. (B) Detail of the $Zn^{2+}$-binding site, showing the three designed His residues, His97, His106 and His110, arranged around the $Zn^{2+}$ ion in a geometry compatible with tetrahedral coordination. The $Zn^{2+}$ ion shows a pLDDT value of 83.1, with a protein–$Zn^{2+}$ chain-pair iPTM of 0.93 and a minimum pairwise PAE of 1.43 Å. The three His residues in the $Zn^{2+}$-binding site also show good prediction confidence, with an average whole-residue pLDDT of 82.8. (C) Detail of the haem-binding site, showing the two designed His residues coordinating the haem Fe atom. The Fe–N(His) distances are 2.0–2.1 Å for the two axial His ligands, consistent with the expected distances for bis-His coordinated haem proteins. The haem molecule shows a pLDDT value of 84.6, with a protein–haem chain-pair iPTM of 0.85 and minimum pairwise PAE values of 1.11–1.49 Å. The designed haem-binding site residues show high prediction confidence, with an average whole-residue pLDDT of 95.1. Overall, the AF3 model shows high global confidence and strong support for the predicted placement of both haem and $Zn^{2+}$ relative to the protein scaffold and the designed binding sites.

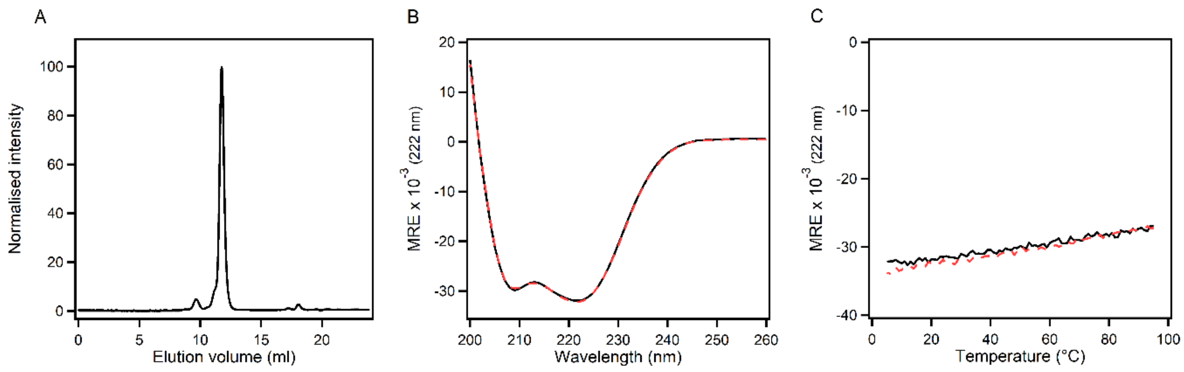


**Figure S9. Biophysical characterization of mCC-4(2)4-ZN-HEM. (A)** Normalized analytical size exclusion chromatograms. Conditions: Superdex™ 75 Increase 10/300 GL, 50 mM HEPES pH 7.0, 150mM NaCl. **(B)** CD spectra recorded at 5 °C. MRE, mean residue ellipticity (deg $cm^2$ $dmol^{-1}$ $res^{-1}$). Conditions: 10 μM protein, 20 mM Phosphate buffer pH 7.4, 150mM NaCl. Solid black line shows CD spectra before melting and dashed red line shows CD spectra after heating up to 95° C and cooling down back to 5° C **(C)** Temperature dependent CD signal, monitored at 222 nm. Scans were collected heating to (solid black line) and cooling from (dashed red line) 95° C. Conditions**:** phosphate buffer pH 7.4, 150mM NaCl.

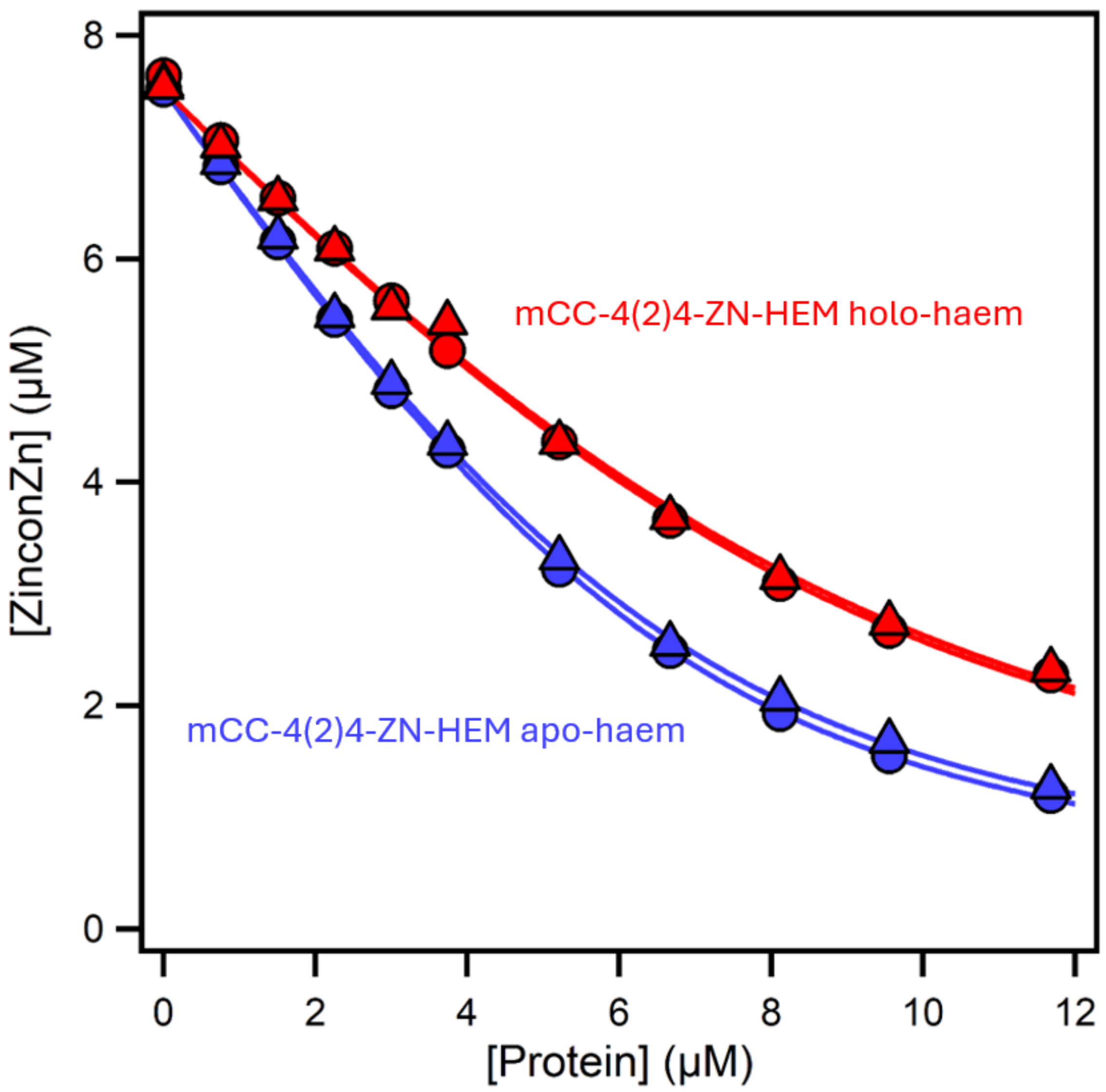


**Figure S10. Zincon competition assays to determine $Zn^{2+}$ binding affinity.** $Zn^{2+}$ binding to the designed dual binder mCC-4(2)4-ZN-HEM was measured by titrating protein into a pre-formed Zincon–$Zn^{2+}$ complex. Competition between the protein and Zincon for $Zn^{2+}$ binding resulted in dissociation of the Zincon–$Zn^{2+}$ complex, which was monitored by absorbance. Blue data points show duplicate titrations of the apo-haem state, whereas red data points show duplicate titrations of the holo-haem state. The data were fitted using a one-site competitive binding model (see supplementary materials). Individual fits for the apo-haem state gave $K_D$ values of 260 ± 10 nM and 280 ± 20 nM, with 1.38 ± 0.01 and 1.37 ± 0.01 effective binding sites, respectively. Individual fits for the holo-haem state gave $K_D$ values of 400 ± 50 nM and 410 ± 80 nM, with 1.06 ± 0.05 and 1.05 ± 0.07 effective binding sites, respectively.

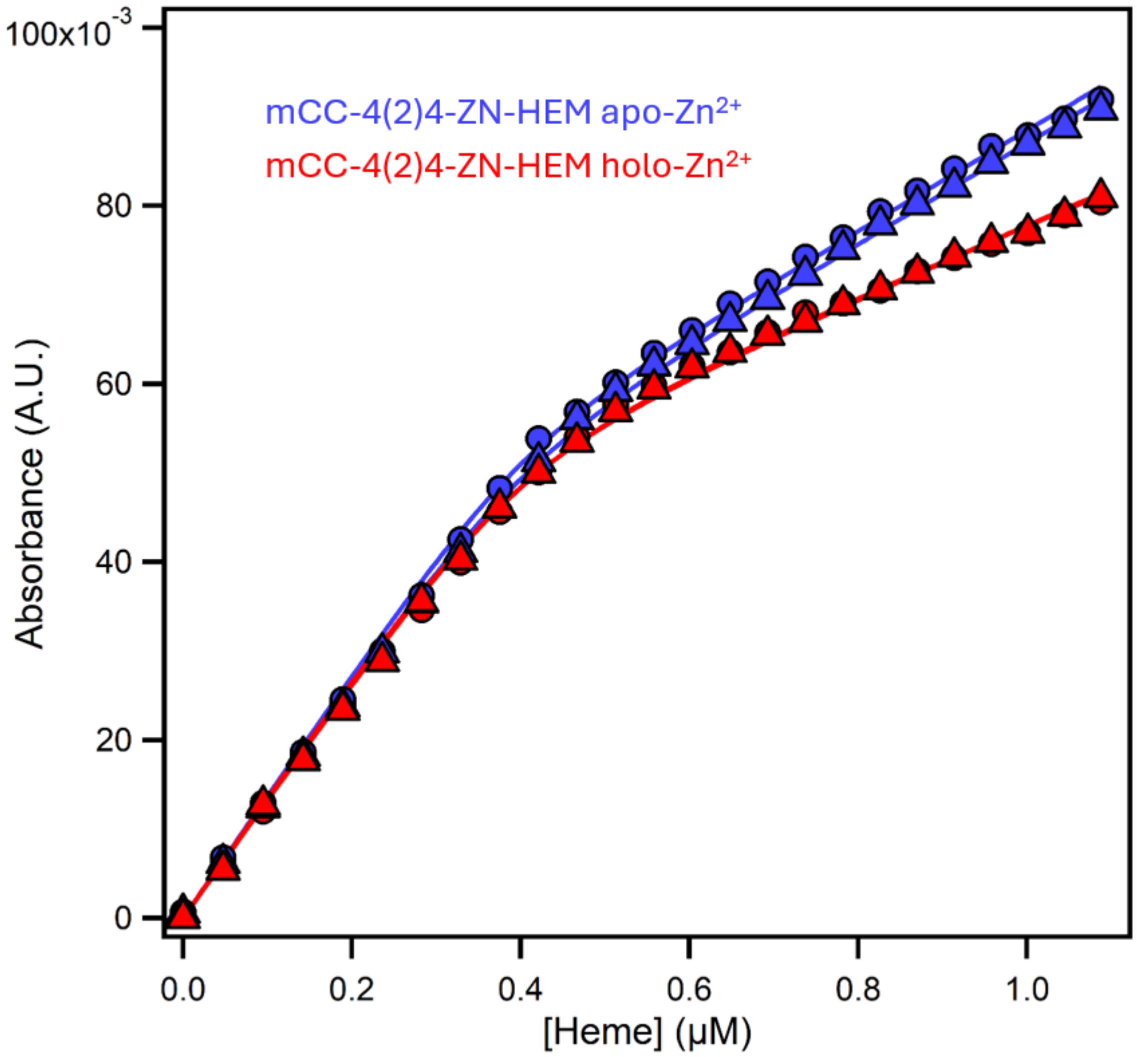


**Figure S11. Haem-binding titrations of mCC-4(2)4-ZN-HEM in the absence and presence of $Zn^{2+}$.** Haem binding was monitored by titrating haem into mCC-4(2)4-ZN-HEM (0.4 μM) and following the absorbance change at 413 nm, corresponding to the Soret maximum of the bound haem. Blue data points show duplicate titrations of the $Zn^{2+}$apo state, whereas red data points show duplicate titrations of the $Zn^{2+}$ holo state. The data were fitted using a one-site binding model. Individual fits for the $Zn^{2+}$ apo state gave apparent $K_D$ values of 12.3 ± 7.6 nM and 9.6 ± 6.9 nM, whereas fits for the $Zn^{2+}$-bound state gave apparent $K_D$ values of 11.3 ± 6.8 nM and 17.3 ± 10.9 nM. In all titrations, the binding transition occurred close to a 1:1 haem ratio, consistent with a single high-affinity haem-binding site. Because the fitted $K_D$ values are substantially lower than the protein concentration used in the assay, these values should be interpreted as apparent estimates rather than precise equilibrium constants[14]. Nevertheless, the transition near stoichiometric haem binding supports tight haem association in both the $Zn^{2+}$-free and $Zn^{2+}$-bound states.

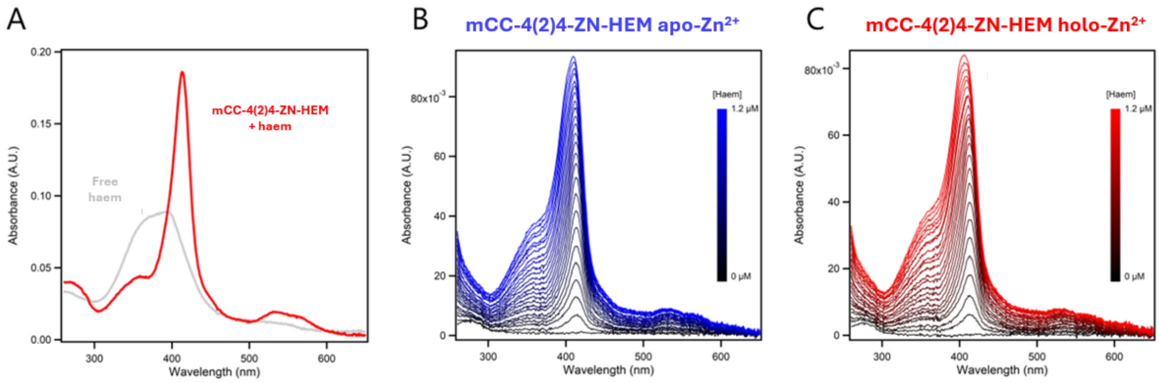


**Figure S12**. **UV–visible characterisation of haem binding by mCC-4(2)4-ZN-HEM.** (A) UV–visible absorbance spectrum of mCC-4(2)4-ZN-HEM in the presence of haem (red), compared with free haem in solution (grey). Both samples contained 1.5 µM haem in 50 mM HEPES, 150 mM NaCl, pH 7.0. Haem binding to mCC-4(2)4-ZN-HEM resulted in a sharp Soret band and resolved α/β bands, consistent with a protein-bound bis-His-coordinated haem species. (B,C) UV–visible absorbance spectra corresponding to the haem titrations used to determine apparent haem-binding affinities for the $Zn^{2+}$-free state (B, apo-$Zn^{2+}$) and $Zn^{2+}$-bound state (C, holo-$Zn^{2+}$) of mCC-4(2)4-ZN-HEM. Haem was titrated into 0.4 µM protein, and spectra were recorded after each addition. The colour gradients indicate increasing haem concentration from 0 to 1.2 µM.

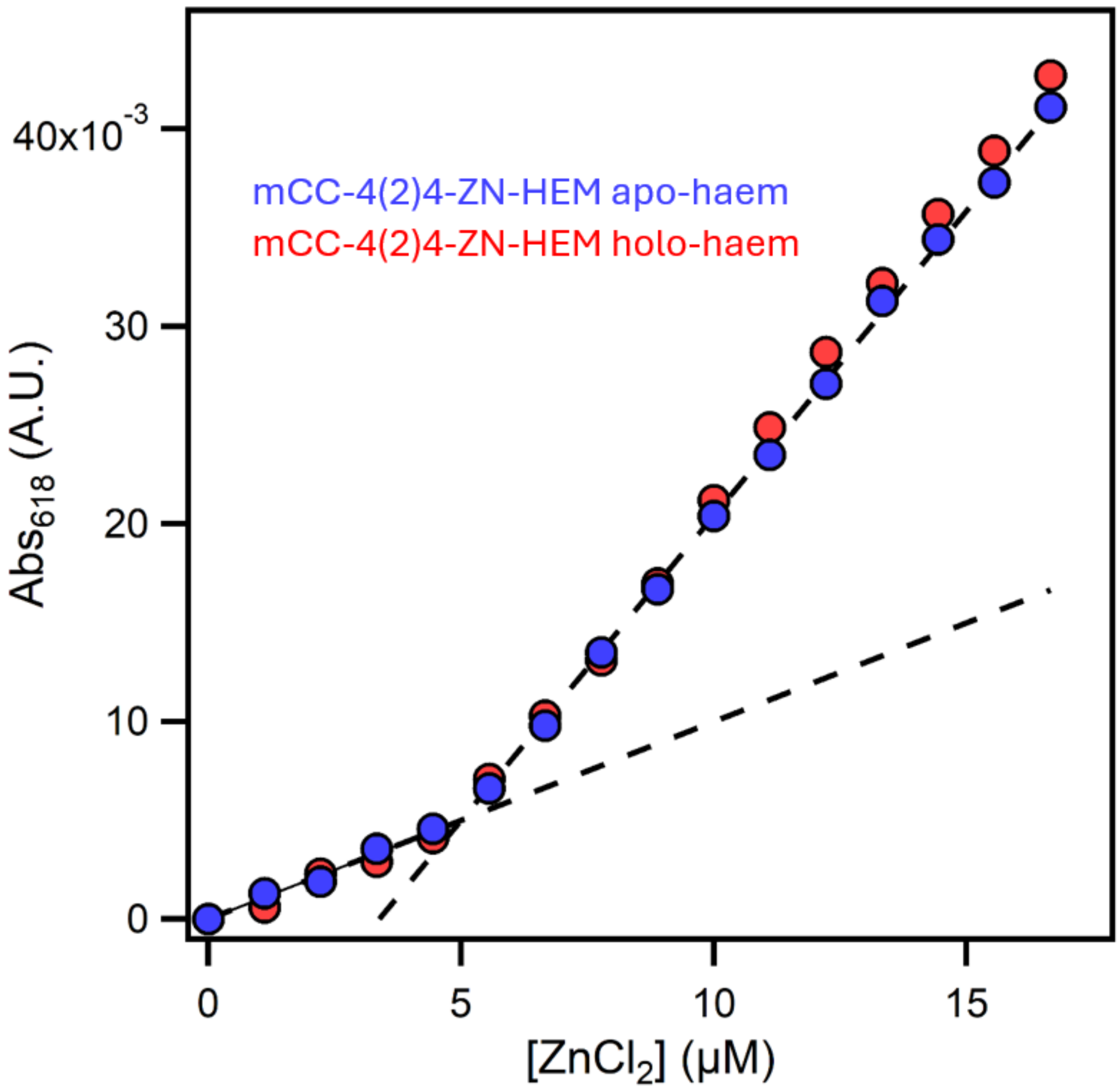


**Figure S13. Determination of $Zn^{2+}$-binding stoichiometry by Zincon competition assay**. $Zn^{2+}$-binding stoichiometry of mCC-4(2)4-ZN-HEM was assessed by titrating $ZnCl_2$ into samples containing 5 μM protein and 10 μM Zincon in 50 mM HEPES, 150 mM NaCl, pH 7.0. Formation of the Zincon–$Zn^{2+}$ complex was monitored by absorbance at 618 nm. Blue data points correspond to the haem-free state of mCC-4(2)4-ZN-HEM, whereas red data points correspond to the haem-bound state. In both cases, the titration showed a single transition at approximately 5 μM $ZnCl_2$, matching the protein concentration and indicating one high-affinity $Zn^{2+}$-binding site per protein. The similar stoichiometry observed in the haem-free and haem-bound states supports the orthogonality of the designed $Zn^{2+}$- and haem-binding sites, with no apparent cross-binding or interference between the two sites under the conditions tested.

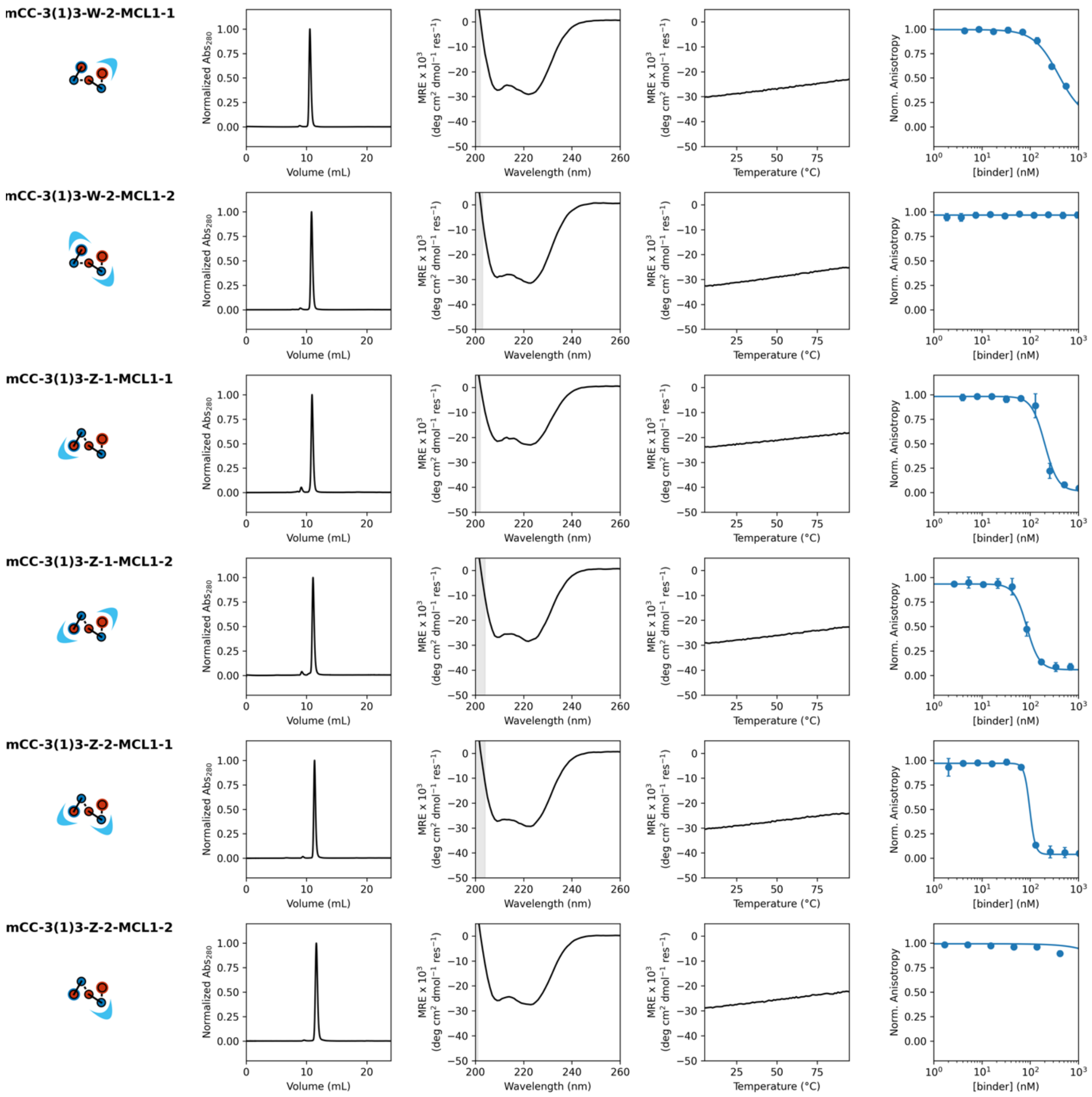


**Figure S14. Biophysical characterization of the MCL-1 binders**. Topology diagrams showing helical interconnectivity. **(B)** Normalized analytical size exclusion chromatograms. Conditions: Superdex™ 75 Increase 10/300 GL, 20 mM Tris pH 7.5, 150mM NaCl. **(C)** CD spectra recorded at 5 °C. Filled grey regions denote hight tension voltage ≥ 600 V. MRE, mean residue ellipticity (deg $cm^2$ $dmol^{-1}$ $res^{-1}$). Conditions: 5-10 µM protein, 20 mM Tris pH 7.5, 150mM NaCl. **(D)** Temperature dependent CD signal, monitored at 222 nm. Scans were collected heating to (solid lines) and cooling from (dashed lines) 95 °C. MRE, mean residue ellipticity (deg $cm^2$ $dmol^{-1}$ $res^{-1}$). **Conditions:** 20 mM Tris pH 7.5, 150mM NaCl. **(E)** MCL-1 binding assay. **Conditions:** 150 nM MCL-1, 25 nM reporter, varying concentration of binders, 20 mM Tris pH 7.5, 150mM NaCl. Returned $IC_{50}$ (top to bottom): 385±25 nM, 5000±265 nM, 200±16 nM, 85±4 nM, 98±3 nM, no binding.

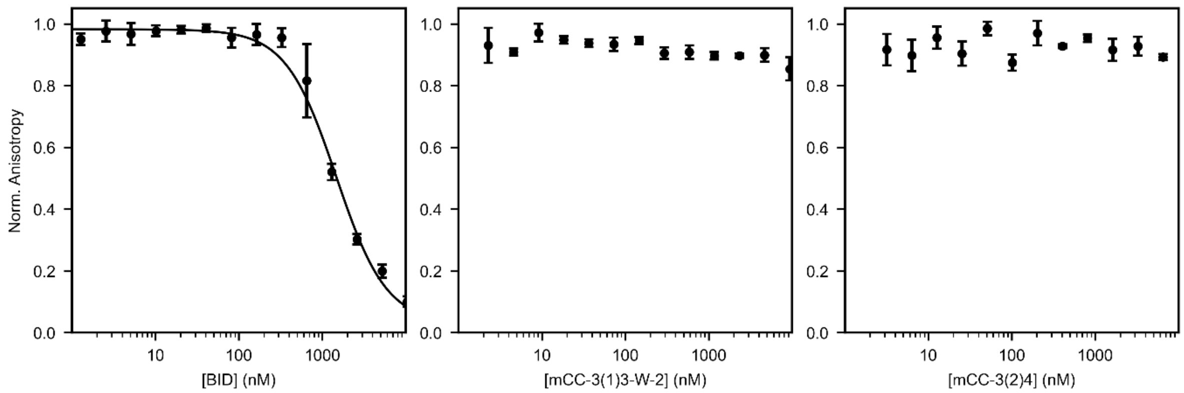


**Figure S15. Fluorescence anisotropy of BID competitor peptide and non-grafted controls.** Normalized anisotropy of competitor with titration of BID, non-decorated 3(1)3 and 3(2)4 controls respectively. **Conditions:** 150 nM MCL1, 25 nM reporter, varying concentration of binders, 20 mM Tris pH 7.5, 150mM NaCl. Returned $IC_{50}$ for BID is 1470 ± 110 nM. Controls show no binding.

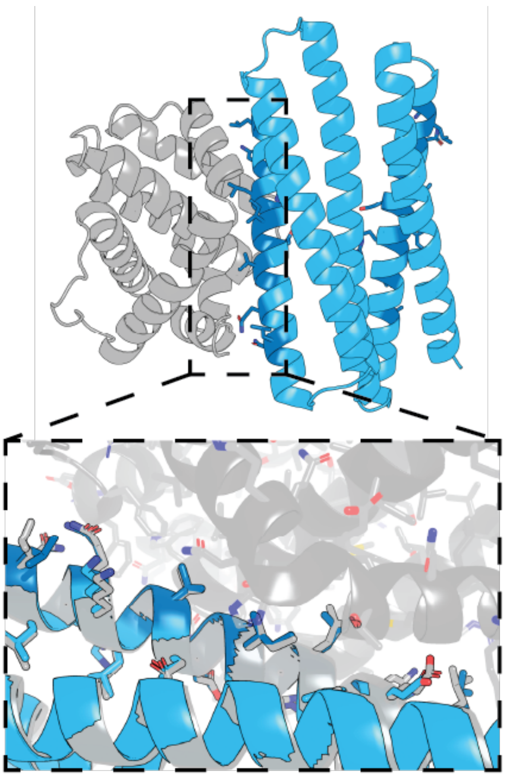

**Figure S16. X-ray crystal structure of mCC-3(1)3-Z-2-MCL1-1 (cyan) in complex with one MCL-1 (grey).** The zoom shows the binding-site residues of the experimental structure (blue) and AlphaFold2 model (grey) as sticks. PDB ID is 31WU.

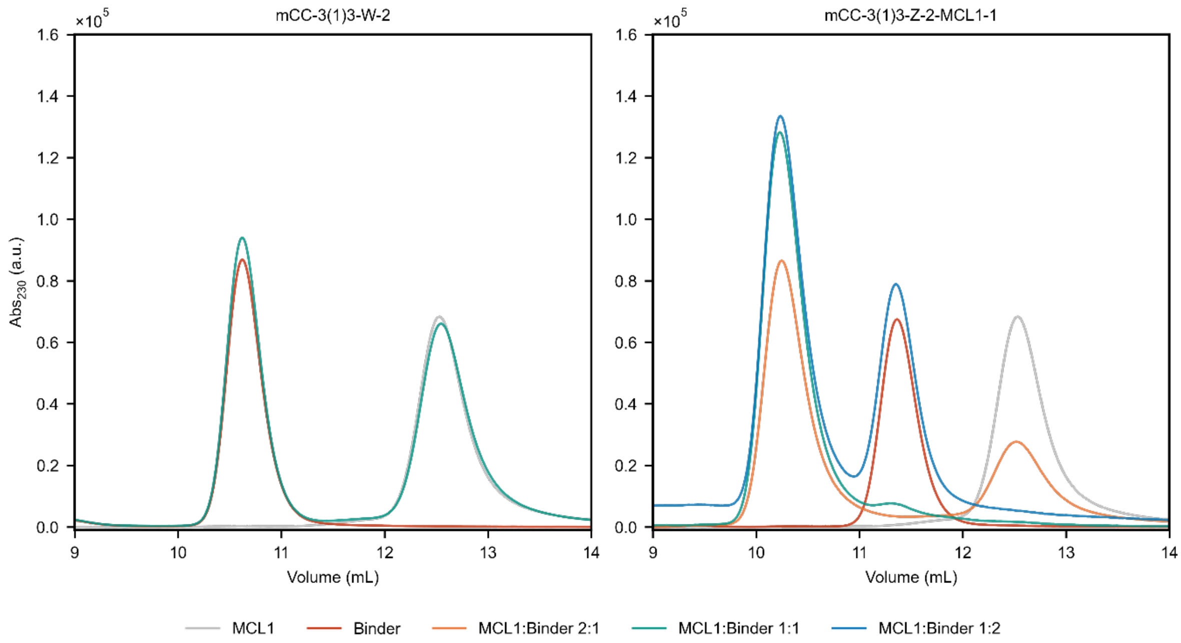


**Figure S17. Size exclusion binding assay.** Absorbance measured at 230 nm for mixture of MCL-1 and mCC-3(1)3-W-2 (non-binding control) and mCC-3(1)3-Z-2-MCL1-1 respectively. **Conditions:** 20 mM Tris pH 7.5 150 mM NaCl, MCL-1 fixed at 30 µM, binder concentration variable ratio (shown in legend), Superdex™ 75 Increase 10/300 GL run at 1 ml/min. Control shows no binding, mCC-3(1)3-Z-2-MCL1-1 shows binding a single copy of MCL-1.

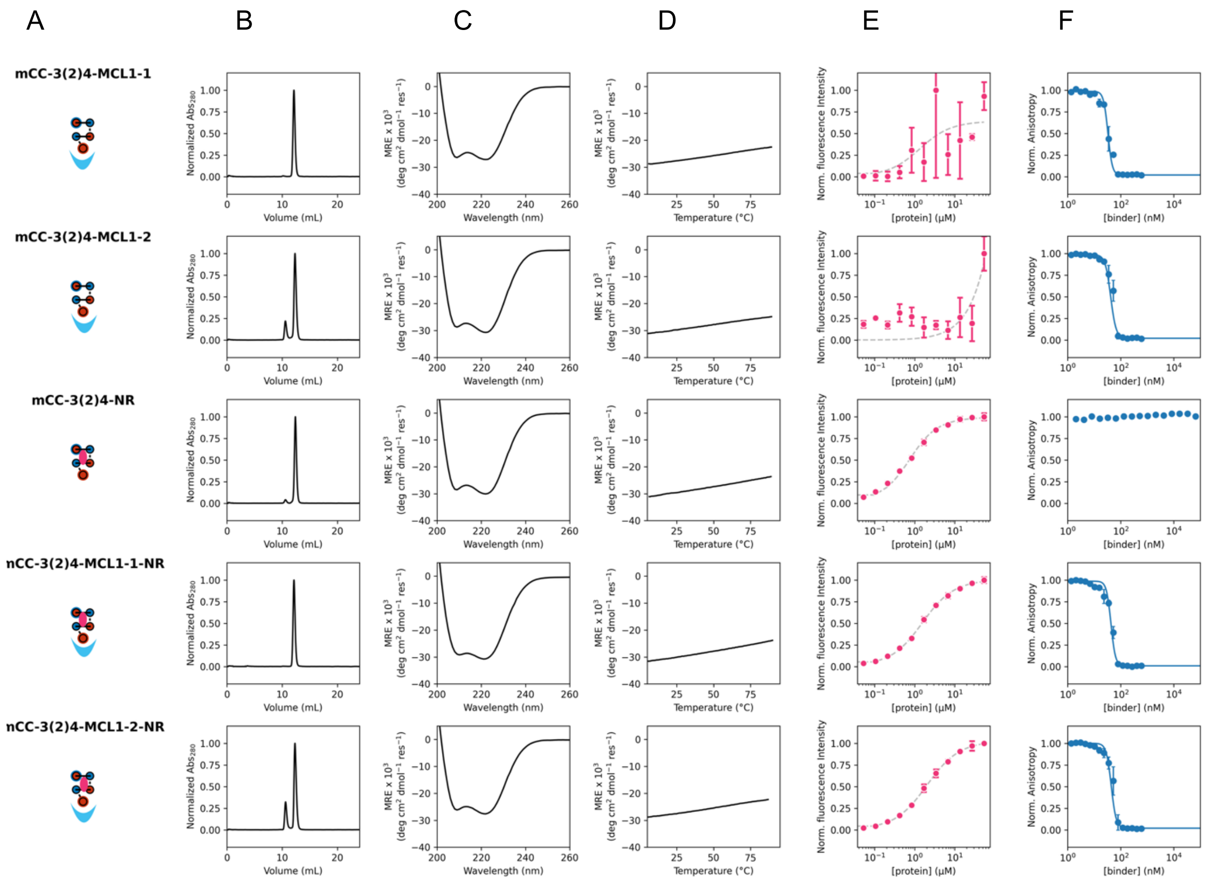


**Figure S18. Biophysical characterization of MCL1 and Nile Red binders**. (**A**) Topology diagrams showing helical interconnectivity. **(B)** Normalized analytical size exclusion chromatograms. Conditions: Superdex™ 75 Increase 10/300 GL, 50 mM phosphate buffer pH 7.4, 150mM NaCl **(C)** CD spectra recorded at 5 °C. Filled grey regions denote hight tension voltage ≥ 600 V. MRE, mean residue ellipticity (deg $cm^2$ $dmol^{-1}$ $res^{-1}$). Conditions: 5-10 μM protein, 50 mM sodium phosphate and 150 mM NaCl (pH 7.4). **(D)** Temperature dependent CD signal, monitored at 222 nm. Scans were collected heating to (solid lines) and cooling from (dashed lines) 95 °C. MRE, mean residue ellipticity (deg $cm^2$ $dmol^{-1}$ $res^{-1}$). **Conditions:** 10 μM protein, 50 mM sodium phosphate and 150 mM NaCl (pH 7.4). **(E)** Nile Red binding curves measured by fluorescence intensity. **Conditions:** 54-0 μM protein, 0.5 μM ligand, 50 mM sodium phosphate and 150 mM NaCl (pH 7.4). Returned $K_D$ (top to bottom): no binding, no binding, 0.48±0.04 μM, 1.3±0.1 μM, 1.8±0.1 μM (DPH. **F)** MCL-1 binding assay. **Conditions:** 150 nM MCL-1, 25 nM reporter, varying concentration of binders, 20 mM Tris pH 7.5, 150mM NaCl. Returned $IC_{50}$ (top to bottom): 33±2 nM, 42±2 nM, no binding, 43±2 nM, 44±4 nM.

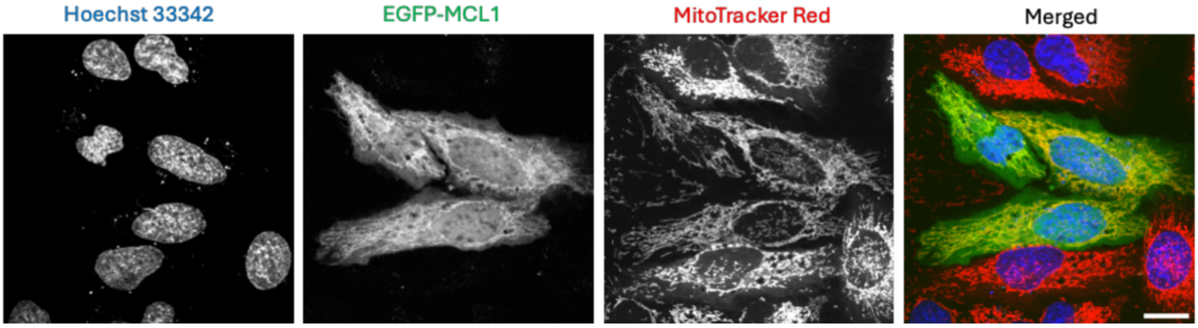


**Figure S19. Overexpressed MCL1 colocalizes with the mitochondrial network, as evidenced by coincident fluorescence of the mitochondrial stain (MitoTracker Red) and EGFP-labelled MCL1.**

#### References

1 Albanese, K. I. *et al.* Rationally seeded computational protein design of α-helical barrels. *Nat Chem Biol* **20**, 991–999 (2024). https://doi.org/10.1038/s41589-024-01642-0

2 Naudin, E. A. *et al.* From peptides to proteins: coiled-coil tetramers to single-chain 4-helix bundles. *Chem Sci* **13**, 11330-11340 (2022). https://doi.org/10.1039/d2sc04479j

3 Dawson, W. M. *et al.* Coiled coils 9-to-5: rational design of α-helical barrels with tunable oligomeric states. *Chem Sci* **12**, 6923-6928 (2021). https://doi.org/10.1039/d1sc00460c

4 Thomson, A. R. *et al.* Computational design of water-soluble α-helical barrels. *Science* **346**, 485-488 (2014). https://doi.org/10.1126/science.1257452

5 Zhou, J. F. & Grigoryan, G. Rapid search for tertiary fragments reveals protein sequence-structure relationships. *Protein Sci* **24**, 508-524 (2015). https://doi.org/10.1002/pro.2610

6 Dauparas, J. *et al.* Robust deep learning-based protein sequence design using ProteinMPNN. *Science* **378**, 49-55 (2022). https://doi.org/10.1126/science.add2187

7 Leng, X. *et al.* De novo designed 3-helix bundle peptides and proteins with controlled topology and stability. *Chem Sci* **16**, 18632-18641 (2025). https://doi.org/10.1039/D5SC05576H

8 Jeong, W. J., Ha, S. & Song, W. J. Accurate computational design of artificial metalloproteins using Metal-Installer. *Chem-Us* **11**, 102644 (2025). https://doi.org/10.1016/j.chempr.2025.102644

9 Ennist, N. M. *et al.* De novo protein design of photochemical reaction centers. *Nat Commun* **13**, 4937 (2022). https://doi.org/10.1038/s41467-022-32710-5

10 Mylemans, B. *et al.* *De novo* designed bifunctional proteins for targeted protein degradation. *bioRxiv*, 2025.2012.2022.695915 (2025). https://doi.org/10.64898/2025.12.22.695915

11 Acevedo-Jake, A. M. *et al.* Grafted Coiled-Coil Peptides as Multivalent Scaffolds for Protein Recognition. *Acs Chem Biol* **20**, 1309-1318 (2025). https://doi.org/10.1021/acschembio.5c00137

12 Petrenas, R. *et al.* Rapid Assessment of Size, Shape, and Chemical Complementarity of Ligands for Computational Protein Design. *bioRxiv*, 2025.2006.2030.662286 (2025). https://doi.org/10.1101/2025.06.30.662286

13 Schneidman-Duhovny, D., Hammel, M., Tainer, J. A. & Sali, A. FoXS, FoXSDock and MultiFoXS: Single-state and multi-state structural modeling of proteins and their complexes based on SAXS profiles. *Nucleic Acids Res* **44**, W424-W429 (2016). https://doi.org/10.1093/nar/gkw389

14 Kalvet, I. *et al.* Design of Heme Enzymes with a Tunable Substrate Binding Pocket Adjacent to an Open Metal Coordination Site. *J Am Chem Soc* **145**, 14307-14315 (2023). https://doi.org/10.1021/jacs.3c02742
